# Quantitative Measurement of Secretory Protein Mistargeting by Proximity Labeling and Parallel Reaction Monitoring

**DOI:** 10.1101/2023.07.19.549095

**Authors:** Ziqi Lyu, Joseph C. Genereux

## Abstract

Proximity labeling is a powerful approach for characterizing subcellular proteomes. We recently demonstrated that proximity labeling can be used to identify mistrafficking of secretory proteins, such as occurs during pre-emptive quality control (pre-QC) following endoplasmic reticulum (ER) stress. This assay depends on protein quantification by immunoblotting and densitometry, which is only semi-quantitative and suffers from poor sensitivity. Here, we integrate parallel reaction monitoring mass spectrometry to enable a more quantitative platform for ER import. PRM as opposed to densitometry improves quantification of transthyretin mistargeting while also achieving at least a ten-fold gain in sensitivity. The multiplexing of PRM also enabled us to evaluate a series of normalization approaches, revealing that normalization to auto-labeled APEX2 peroxidase is necessary to account for drug treatment-dependent changes in labeling efficiency. We apply this approach to systematically characterize the relationship between chemical ER stressors and ER pre-QC induction in HEK293T cells. Using dual-FLAG-tagged transthyretin (^FLAG^TTR) as a model secretory protein, we find that Brefeldin A treatment as well as ER calcium depletion cause pre-QC, while tunicamycin and dithiothreitol do not, indicating ER stress alone is not sufficient. This finding contrasts with the canonical model of pre-QC induction, and establishes the utility of our platform.

**TOC graph:** 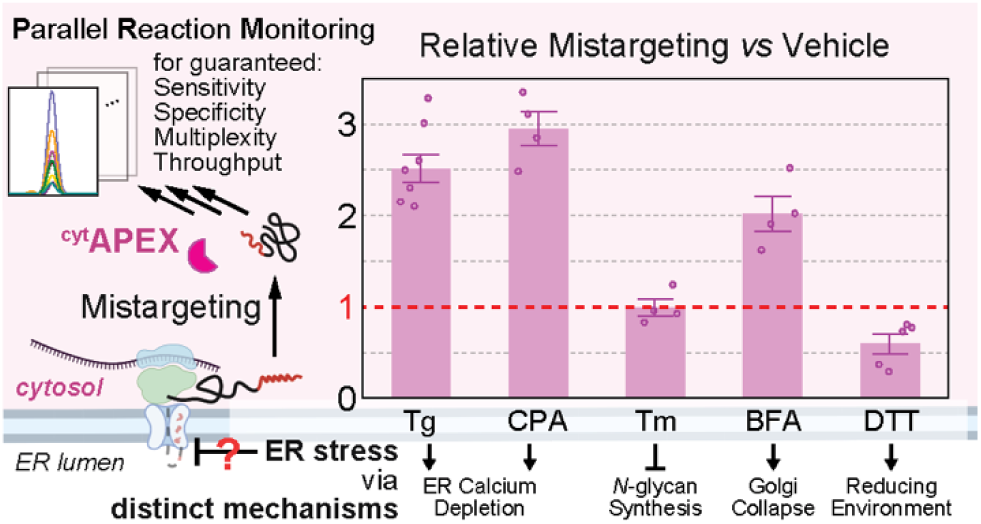

## INTRODUCTION

Eukaryotic cells depend upon the secretory pathway to properly traffic about one-third of their proteome^1^, including nearly all secreted and plasma membrane proteins. As the first compartment of the secretory pathway, the endoplasmic reticulum (ER) maintains a calcium-rich environment for calcium binding^2^, an oxidative environment for disulfide bond formation^3^, and possesses a unique set of enzymes and chaperones for glycoprotein biogenesis and quality control^4^. Secretory proteins have evolved to rely on this unique folding environment, and hence if mistargeted, these proteins present a threat to the cytosolic proteostasis^5^. Multiple checkpoints and quality control steps ensure high fidelity of translocation of secretory proteins^6^. In the presence of ER stress, translocation for some secretory proteins is attenuated, leading to their cytosolic mislocalization. These mistargeted proteins are primarily directed towards degradation^7–10^. This process is termed ER pre-emptive quality control^11,12^ (sometimes denoted ER pQC; we use ER pre-QC instead, to avoid confusion with generic protein quality control PQC^13–15^). Because current techniques for measuring protein mislocalization are onerous, semi-quantitative, or *in vitro*^16,17^, we do not yet have clear understanding of the substrates and biochemical mechanisms of ER pre-QC^18^, nor which stresses activate it.

Proximity labeling has emerged as a technique of choice for characterizing subcellular proteomes^19^ and protein trafficking^20–24^. We recently demonstrated that proximity labeling is an effective method for identifying mistargeting of secretory proteins^25,26^. In this approach, N-terminal FLAG-tagged APEX2 with a nuclear export signal (NES)^27,28^ is expressed and localized in the cytosol (^cyt^APEX). Upon initiation of labeling reactions with a 1-min H_2_O_2_ pulse, cytoplasmic proteins are biotin-phenol (BP)-labeled, and these BP-labeled proteins can be affinity purified. Secretory proteins that mistarget and accumulate in the cytosol are labeled and purified as well, and the relative amount of mistargeted protein can be determined by immunoblotting (IB). While this assay allows easy measurement of protein mistargeting under stress, the use of IB introduces several limitations. The relatively limited sensitivity of IB necessitates the use of several million cells for each drug treatment condition. Proximity labeling can hide epitopes, particularly on the most popular affinity tags, such as the FLAG tag DYKDDDDK and the hemagglutinin tag YPYDVPDYA.^29,25^ In addition to the availability, specificity, and sensitivity of antibodies, good quality of gel electrophoresis and electrotransfer must be assured for reliable quantification. If assayed proteins fall in a wide range of molecular weights, for the sake of resolution, more gels of different acrylamide/bis-acrylamide compositions should be used. If some assayed proteins share similar molecular weights and the same host animal of the available antibody, another gel or stripping must be considered. Only a few antibodies can be employed per gel, limiting the potential for multiplexed detection. Finally, IB is a time-consuming process.

Targeted mass spectrometry methods like parallel reaction monitoring^30^ (PRM) avoid these problems and have some advantages over IB^31^. Peptides can be chosen to avoid those sensitive to peroxidase labeling. During data acquisition, these pre-defined peptides are isolated according to their mass-to-charge (*m*/*z*) ratios with a pre-set isolation window. Isolated precursor peptide ions are fragmented to generate product ions and all resulting product ions are analyzed in *parallel* with a mass analyzer that allows MS^2^ full scan (*e.g.* Orbitrap, time-of-flight^32^ or linear ion trap^33^). Just as antibodies must be carefully validated for their specificity for the protein of interest, PRM transitions must be carefully validated for specificity^34^. This is particularly easy for PRM, as multiple product ions of each isolated precursor are analyzed. If fragment ions (conventionally at least three^35^) associated with each precursor peptide maintain similar chromatographic profiles, constant proportion of fragment ion intensity and the same retention time, they can be considered as *bona fide* product ions for later quantification. In addition to the high sensitivity and specificity of (tandem) mass spectrometry, PRM also benefits from liquid chromatography in that it allows multiplexing of hundreds of peptides and inferably hundreds of proteins with ease^32^. Quantification of more than 1000 peptides from a single run has now been achieved, using internal standard-triggered scheduling with modern instrumentation.^36^

Herein, we integrate PRM mass spectrometry with our assay to quantify mistargeting of the model secretory protein transthyretin (TTR) in response to ER stress by distinct mechanisms (**Figure 1**). Using one-order-of-magnitude-less sample, we obtain the same quantification results as are seen by IB^25^. We compare multiple normalization factors and demonstrate the necessity to have a control for proximity labeling efficiency. For drug treatments that do not change proximity labeling efficiency, most normalization factors yield the same result. For treatment that changes labeling efficiency, normalization to auto-labeled APEX2 peptides may be the most accurate method. With the PRM assay and proper data normalization, we establish that not all ER stressors induce ER pre-QC in HEK293T cells. Rather, only Brefeldin A (BFA) and sarcoplasmic/endoplasmic reticulum calcium ATPase (SERCA) inhibitors thapsigargin (Tg) and cyclopiazonic acids (CPA) induce ER pre-QC. While tunicamycin (Tm) or 1,4-dithiothreitol (DTT) induce ER stress, they do not increase relative ^FLAG^TTR mistargeting in the cytosol. Hence, we show that PRM-based quantification of secretory protein mistargeting can be used to determine the factors responsible for pre-QC in living cells.

**Figure 1.**
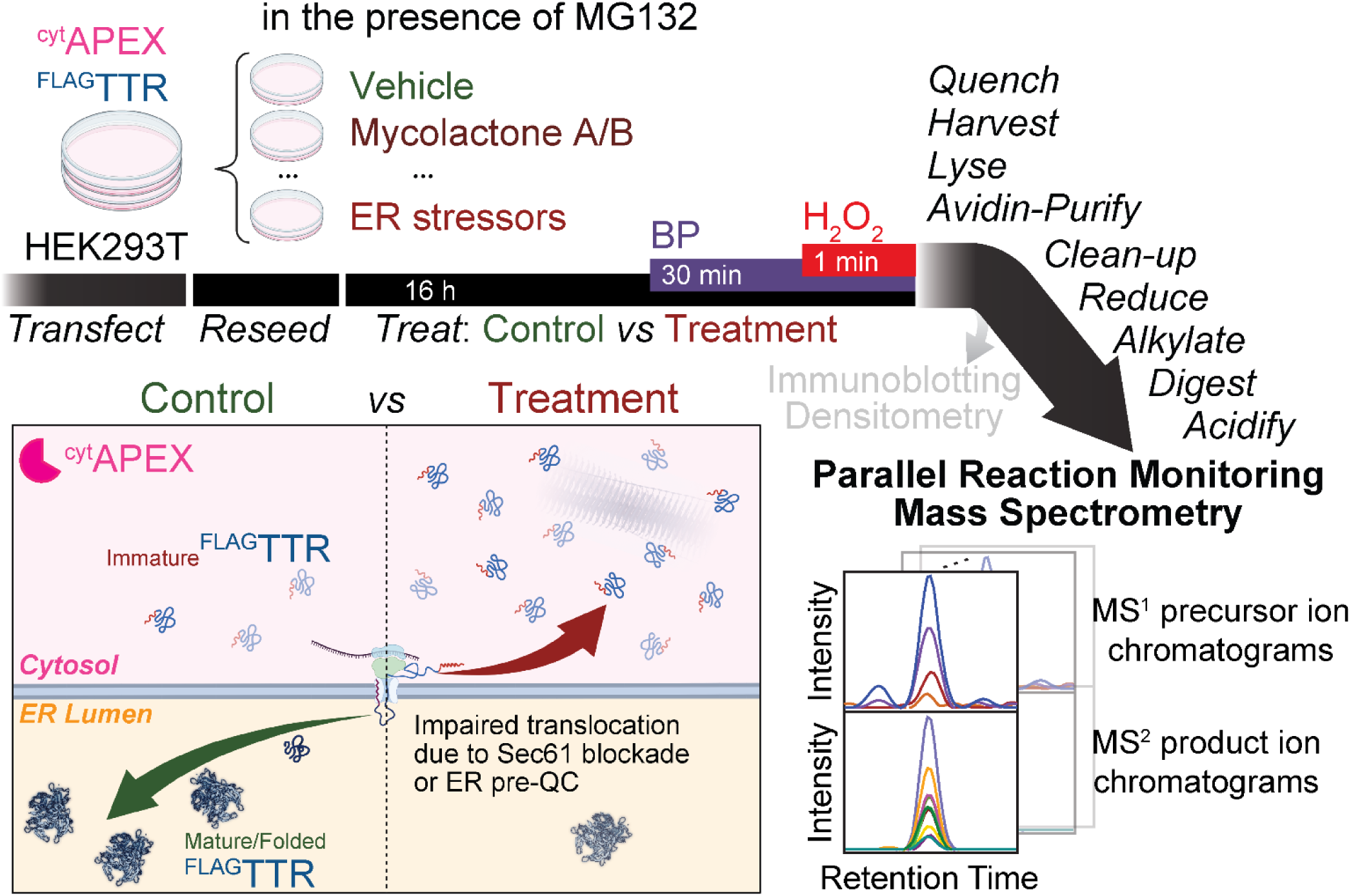
Proposed experimental workflow of this assay. ^cyt^APEX and ^FLAG^TTR are transiently transfected into HEK293T cells via calcium phosphate transfection. Cells are reseeded for later drug treatment (16 h). We expect increased differential ^FLAG^TTR mistargeting to be observed under the condition of Sec61 blockade (during mycolactone A/B treatment) or ER pre-QC induction (during ER stress). 30 min before the H_2_O_2_ pulse, biotin-phenol (BP) is added. The 1-min BP-labeling reaction is quenched by washing cells on ice with quencher solution 3 times. Cells are then harvested and lysed; cell lysates are brought to the same mass concentration and subjected to affinity purification with avidin agarose beads. Instead of loading eluate samples for SDS-PAGE and IB, we pellet avidin-enriched proteins by MeOH/CHCl_3_ precipitation and get the protein pellets washed with MeOH to clean-up, resuspended, reduced, carbamidomethylated and trypsin-digested, followed by acidification and clarification by hard spinning. Eluate digests are then analyzed by parallel reaction monitoring mass spectrometry. Displayed at the right bottom corner are schematic MS^1^ precursor ion chromatograms and MS^2^ product ion chromatograms of a targeted peptide. Areas under chromatograms are used for quantification.

## EXPERIMENTAL SECTION

The ambient temperature in our lab is 17–21°C. Buffer components and other biochemical reagents were all purchased from Fisher, VWR, or Millipore Sigma. Nanopure water and sterilized consumables were used for all biochemical experiments. Cell culture media and cell detachment solution (0.25% trypsin, 0.1% EDTA, w/v; Corning) are pre-warmed before use in a 37°C aluminum bead bath. The heat block for denaturing protein samples is set to 100 °C.

### Human tissue culture

HEK293T cells (American Type Culture Collection) were cultured in Dulbecco’s modified Eagle’s medium (DMEM, Corning) supplemented with 10% (v/v) fetal bovine serum (FBS, Seradigm), 2 mM L-glutamine (Corning), and penicillin (100 IU mL^−1^)-streptomycin (100 μg mL^−1^, Corning), and used within 30 passages after thawing.

### Transfection and reseeding

Desired exogenous plasmid DNAs were introduced into HEK293T cells via calcium phosphate transfection, with media changed 12−16 hours post-transfection. At least one hour after media change, cells were reseeded into poly D-lysine-treated 6-cm dishes to ensure cellular retention during later treatments and washing. Poly D-lysine treatment was performed by coating plates with poly D-lysine hydrobromide (0.1 mg mL^−1^ in H_2_O from lyophilized powder, Sigma-Aldrich) for 15 min, then washing twice with Dulbecco’s phosphate-buffered saline (DPBS, 1×, HyClone, GE) before adding cell culture media.

### Drug treatment and proximity labeling

After cellular attachment, transfected cells were treated with corresponding drugs by changing media, as summarized in **Table S1**. On the second day post-transfection, dimethyl sulfoxide (DMSO, as vehicle, tissue culture grade, Corning) or biotin-phenol (BP, 500 μM, from 0.5 M stock in DMSO, prepared in lab as described^26^) were added to the cells through conditioned media containing residual drugs from initial treatment. Cells were incubated at 37°C for 30 min. 1 M sodium (+)-L-ascorbate (in H_2_O, as 100× stock, Sigma), 0.5 M Trolox (in DMSO, as 100× stock, Acros) and 100 mM H_2_O_2_ (in 1× DPBS, as 100× stock, from 9.8 M, Fisher) were freshly prepared during or before the 30-min BP incubation. 1× quencher solution was made by diluting 1 M sodium azide (NaN_3_, in H_2_O, as 100× stock, Fisher) to 10 mM, 1 M ascorbate to 10 mM, 0.5 M Trolox to 5 mM, with 1× DPBS and chilled on ice.

After the 30-min BP incubation, 30 μL 100 mM H_2_O_2_ was added into each dish to a final concentration of 1 mM, and dishes were agitated immediately after addition. For DTT-treated cells, conditioned media containing DTT was aspirated before the replacement by pre-warmed DMEM containing 1 mM H_2_O_2_. Exactly 1 min after the H_2_O_2_ delivery, media were aspirated, and cells were washed three times with 3 mL ice-cold 1× quencher solution and kept on ice. Quenched cells were then harvested in 1 mL 1× quencher solution by scraping and pelleted at 4°C, 700× *g* for 5 min. Cell pellets that were not immediately processed were stored in the freezer (≤ −60°C).

### Cell lysis

Freshly harvested or thawed cell pellets were lysed on ice for at least 10 min with 1× quenchers (10 mM NaN_3_, 10 mM sodium (+)-L-ascorbate, 5 mM Trolox) and protease inhibitors cocktail (PIC, Roche) in radioimmunoprecipitation assay (RIPA) buffer (50 mM Tris pH 7.5, 150 mM NaCl, 1% (w/v) Triton X-100, 0.5% (w/v) sodium deoxycholate, 0.1% (w/v) SDS). Lysates were clarified by centrifugation at 21,100× g for 15 min at 4°C. Soluble protein concentration was determined by colorimetric assay (Bio-Rad) using Agilent Cary 60 UV-Vis spectrophotometer, and lysate concentration was normalized to the lowest sample. SDS-PAGE samples were prepared in reducing Laemmli buffer (mix with 6× stock, 12% SDS, 0.01% bromophenol blue, Acros or Fisher, 47% (w/v) glycerol, Fisher, 60 mM Tris pH 6.8; DTT was freshly added immediately before use) followed by 10-min boiling. Samples with TTR were boiled for 20 min to break up aggregated material.^25,26^

### Avidin purification

BP-labeled proteins were affinity purified from mass-balanced lysates with RIPA-rinsed avidin agarose beads (Pierce, 30 μL slurry per sample) and rotated overnight at 4°C. Beads were then washed twice with RIPA, once with 1 M KCl/0.1% (w/v) Triton X-100 in H_2_O, once with 0.1 M Na_2_CO_3_/0.1% (w/v) Triton X-100 in H_2_O, once with 2 M urea/0.1% (w/v) Triton X-100 in 10 mM Tris pH 8.0, and twice with RIPA to decrease non-specific binding. BP-labeled proteins were eluted in denaturing elution buffer (12% (w/v) SDS, 0.01% (w/v) bromophenol blue, 7.8% (w/v) glycerol, 10 mM Tris pH 6.8, stored at ambient temperature; 2 mM D-(+)-biotin and 20 mM DTT added to the elution buffer freshly before use) by boiling for 10 min. Collected eluates were boiled for another 10 min.

### Gel electrophoresis, immunoblotting, and electrochemiluminescence

SDS-PAGE was performed on 12% Tris-glycine gels (from 30% (w/v) acrylamide/bis-acrylamide, 37.5:1, w/w, Bio-Rad, or from acrylamide powder, Sigma and bis-acrylamide powder, Bio-Rad). Approximately 40 μg protein was loaded in input gels; 20 μL (60% v/v of elution buffer used) eluate was loaded in eluate gels. Proteins were transferred to nitrocellulose membrane (Bio-Rad) by semi-dry transfer (Turboblot, Bio-Rad). After visualization of total protein by Ponceau S (0.1% (w/v) in 5% (v/v) acetic acid (AcOH)/H_2_O, from powder, Acros) to confirm loading and transfer, membranes were blocked with 5% (w/v) non-fat dried milk (Walmart) in Tris-buffered saline (TBS, 10 mM Tris pH 7.0, 150 mM NaCl) 40 to 60 min at ambient temperature or overnight at 4°C. Rinsed membranes were incubated in primary antibody solution (primary antibody reserved in 5% bovine serum albumin, BSA, Sigma, 0.1% (w/v) NaN_3_ in TBS) for ≥ 2 h at ambient temperature or overnight at 4°C, rinsed well with TBS with 0.1% Tween 20 (Fisher, TBST), incubated in secondary antibody solution (50 ng mL^−1^ in 5% (w/v) non-fat milk/TBS) 20 to 30 min at ambient temperature. Blots were rinsed three times with TBST, once with TBS, and once with H_2_O, followed by imaging on a LI-COR Fc Odyssey imager and analyzed with Image Studio Lite software (LI-COR). Quantification was done using background-subtracted densitometric data of each band of interest.

Antibodies for IB include: polyclonal rabbit anti-GRP78/BiP (1:1000, from 86 μg per 150 μL stock, Proteintech), anti-HSP70/HSPA1A (1:5000, from 24.0 μg per 150 μL stock, Proteintech), anti-human prealbumin/TTR (1:1000, 2.0 g L^−1^, Dako), anti-DNAJB11/ERdj3 (1:1000, from 27 μg per 150 μL stock), Proteintech), anti-GAPDH (1:1000, 42 mg mL^−1^, Cell Signaling Technology or 1:1000, 600 μg mL^−1^, Proteintech) followed by secondary goat anti-rabbit antibody (IRDye 800 CW, 1:10000–20000, from 0.5 mg mL^−1^, LI-COR). Monoclonal mouse anti-KDEL (1:500, 1 mg mL^−1^, Enzo), M2 anti-FLAG tag (1:1000, from 1 mg mL^−1^, Sigma), anti-β-actin (1:5000, from 86 or 150 μg (150 μL)^−1^, Proteintech), and anti-α-tubulin (1:5000, from 260 μg (150 μL)^−1^, Proteintech) followed by secondary goat anti-mouse antibody (IRDye 680 RD, 1:10000–20000, from 0.5 mg mL^−1^, LI-COR).

For electrochemiluminescence, nitro-cellulose membranes were incubated in HRP-conjugated streptavidin (Thermo, 1.25 mg mL^−1^, 1:5000 dilution in 1% milk/TBST) for 4 h at room temperature, followed by washing three times with TBST, once with TBS, and once with H_2_O. Membranes were then drained and placed on the image tray. ECL substrate and peroxide (Cytiva) were mixed, applied to the entire membrane and drained before acquisition on the LI-COR Fc Odyssey imager.

### Mass spectrometry

Only MS grade organic solvents were used during sample preparation, except chloroform (CHCl_3_, certified ACS). Buffer A is 0.1% formic acid in 5% acetonitrile (ACN)/H_2_O, v/v. Buffer B is 0.1% formic acid in 80% ACN/H_2_O, v/v.

*Sample clean-up:* Samples (100 μg lysates or 40 μL eluates) were transferred to low-protein-binding microcentrifuge tubes and brought up to 100 μL with H_2_O and mixed well by vortex mixer at slowest mode. 400 μL MeOH, 100 μL CHCl_3_ and 300 μL H_2_O were added sequentially, with gentle vortex mixing after each addition. After centrifugation twice at 12,500× *g* for 5 min, protein pellets form between the interface of the two liquid phases. Majority of the upper liquid layer was removed carefully by aspiration. The remnant was cleaned by adding 400 μL MeOH, vortex mixing, hard spinning for at least 15 min and supernatant aspiration, repeated ≥ 3 times. Protein pellets were dried in air.

*Sample preparation from protein resuspension using Rapigest:* Dried protein pellets were resuspended in 3 μL 1% Rapigest in H_2_O, followed by addition of 47 μL 100 mM HEPES, pH 8.0. Proteins were then reduced by 10 mM tris(2-carboxyethyl)phosphine (TCEP, Millipore Sigma) for 30 min at 37°C, alkylated by 5 mM iodoacetamide (Millipore Sigma) for 30 min in dark at ambient temperature and digested by trypsin (Thermo Fisher Scientific, final concentration 0.01 μg μL^−1^) overnight (16–24 hours) at 37°C with 600-rpm agitation. Tryptic digestion was quenched by adding formic acid (Acros) to pH 2.0. Acidified samples were heated at 37°C for 1 hour and hard spun for 30 min to precipitate Rapigest decomposition products. Clarified samples were transferred to new low-protein-binding tubes. This process of heating and hard spinning was repeated twice. Samples were stored in freezer ≤ –50°C until analysis.

*Sample preparation from protein resuspension using pH-buffered 9 M urea:* Dried protein pellets were resuspended in 9 M urea in 25 mM NH_4_HCO_3_, pH 7.8 (or 50 mM Tris, pH 8.0). Proteins were then reduced by 10 mM TCEP in 200 mM NH_4_HCO_3_, pH 7.8 (or 50 mM Tris pH 8.0) for 30 min at 37°C, alkylated by 10 mM iodoacetamide in 25 mM NH_4_HCO_3_, pH 7.8 (or 50 mM Tris pH 8.0) for 30 min in dark at room temperature. Samples were diluted with 25 mM NH_4_HCO_3_, pH 7.8 (or 50 mM Tris pH 8.0) to ≤ 2 M urea and brought to 1 mM CaCl_2_, before digested by trypsin (Thermo Fisher Scientific, final concentration 0.01 μg μL^−1^) overnight (16–24 hours) at 37°C with agitation. For sample volume greater than 60 μL, we used an orbital shaker at 37°C and 600 rpm. For samples of 20 μL, we placed those samples inside a 37°C incubator, with 300 rpm agitation, to avoid water evaporation and condensation at the EP tube cap. Tryptic digestion was quenched by adding formic acid (Acros) to pH 2.0 and digests stored in freezer (≤ –50°C) until analysis.

*Column Preparation:* Monophasic C18 loading columns were prepared by polymerizing a Kasil 1624 (next advance) frit into a 150-µm-inner-diameter fused silica capillary (Agilent) and then packing with 2.5-cm-long reversed-phase 5 µm Aqua C18 resin (125 Å, Phenomenex). C18 loading columns were washed with MeOH and Buffer A prior to analysis. Analytical columns were prepared by pulling 100 μm diameter fused silica columns (Agilent) with a P-2000 laser tip puller (Sutter Instrument Co., Novato, CA), followed by packing with at least 15-cm reversed-phase 3 µm Aqua C18 resin (Phenomenex).

*Parallel Reaction Monitoring Mass Spectrometry:* 2 μL digest was analyzed using nLC-1000 (Thermo) with a 100-min ACN gradient (5 min from 1% B to 6% B, 15 min to 12% B, 25 min to 18% B, 35 min to 33% B, 5 min to 100% B, 5 min at 100% B, 5 min to 1% B, 5 min at 1%, 100 min in total; 500 nL/min flow rate). Eluted peptides were ionized by electrospray (3.0 kV) and scanned from 110 to 2000 m/z in the Orbitrap with resolution 30000 in MS^1^ at scheduled 10-min-long window. Targeted precursors were isolated (isolation window 2.0 *m*/*z*) and fragmented by collision-induced dissociation (CID, normalized collision energy NCE 35%, activation time 10 ms) in the ion trap, and detected in the orbitrap with a resolution of 7500. Raw data were imported into and analyzed with Skyline^37^. Peak boundaries for integration were manually inspected and adjusted if necessary to include the entire peak. Where indicated, normalization was performed by dividing raw TTR peptide peak areas by either the TIC or raw peptide peak areas of the indicated normalization factor. The MS proteomics data and associated results files have been deposited to the Panorama Repository^38^ and are available at https://panoramaweb.org/GenereuxLab_MistargetingAssay.url

*Data Dependent Acquisition:* 15 μL avidin-purification digest from HEK293T cell expressing eGFP.N2 was analyzed using the same interface and LC gradient as in PRM method. Eluted peptides were ionized by electrospray (3.0 kV) and scanned from 110 to 2000 m/z in the Orbitrap with resolution 30000 in data dependent acquisition mode. The top ten peaks from each full scan were fragmented by higher energy C-trap dissociation (HCD) using a normalized collision energy of 38%, a 100 ms activation time, and a resolution of 7500. Dynamic exclusion parameters were 1 repeat count, 30 ms repeat duration, 500 exclusion list size, 60 s exclusion duration, and 1.50 Da exclusion width. MS^1^ and MS^2^ spectra were searched with MSFragger (with FragPipe^39,40^) against a combined database of Uniprot human proteome database (downloaded with FragPipe, 2021-07-09), ^cyt^APEX, ^ER^HRP and chicken avidin, and reverse sequences for each entry as the decoy set, with common contaminants (*e.g*. keratin, porcine trypsin, *etc.*). Closed searches were allowed for static modification of cysteine residues (57.02146 Da, carbamidomethylation), variable modification of methionine (15.9949 Da, oxidation), N-terminal free amino group (42.0106 Da, acetylation) and tyrosine residues (361.14601 Da, +BP), half tryptic peptidolysis specificity, and mass tolerance of 20 ppm for precursor mass and 20 ppm for product ion masses. Spectral matches were assembled and filtered with a false discovery rate (FDR) of 0.01.

### Statistics

For quantification of IB or PRM experiments of same types of conditions, we normalized individual densitometric signal or MS^1^ TIC-normalized peak area by the sum of all conditions. For the comparison across different PRM experiments with distinct drug treatment conditions, we divided individual raw peak area or global standard-normalized peak area datum to that of (MG132 and Veh.) sample (fold change) across 10 experiments. To be conservative, these fold changes were subjected to two-tailed heteroscedastic *t*-test in Excel, with Bonferroni correction (6 comparisons).

## RESULTS AND DISCUSSION

ER pre-QC has been described as a general protective mechanism of the ER in the presence of ER stress, and it is presumably regulated by activation of the ER unfolded protein response (UPR).^11,18,41^ This model suggests that all ER stressors should induce ER pre-QC to a similar extent. We used proximity labeling combined with immunoblotting to determine whether ER stress, independently of the mechanism by which it is activated, always mistargets ^FLAG^TTR (a known pre-QC substrate^12,25^) into the cytosol. In addition to Tg, which induces ER stress through depletion of ER Ca^2+^, we considered the well-studied small molecule ER stressors tunicamycin (Tm), 2-deoxy-D-glucose (2-DG), Brefeldin A (BFA) and 1,4-dithiothreitol (DTT). Tm inhibits GlcNAc-1-phosphate transferase, blocking the first step of *N*-glycosylation.^42^ Tm treatment leads to glycoprotein misfolding inside the ER and activation of ER UPR, and it is also a reported ER pre-QC inducer in HepG2 cells^12^. Different from Tm, 2-DG inhibits *N*-glycosylation due to its aberrant incorporation into the *N*-glycan, in place of mannose.^43^ BFA leads to *cis*-Golgi cisternae collapse into the ER and a complete loss of ER-to-Golgi transport and canonical protein secretion.^44–46^ DTT is a cell-penetrable reductant that triggers ER stress by preventing disulfide bond formation inside the ER.

### Immunoblotting provides inadequate sensitivity for quantifying mistargeted protein

We treated HEK293T cells co-expressing ^cyt^APEX and ^FLAG^TTR with chemical ER stressors (Tg, Tm, BFA, DTT, or 2-DG) in the presence of MG132 and determined the relative mistargeted (cytosolic) ^FLAG^TTR under each condition using proximity labeling and IB. Only Tg increases ^FLAG^TTR mistargeting relative to vehicle treatment (**Figure S1**, avidin-purification IB: TTR). BFA and 2-DG did not induce as much BiP expression (a UPR target) under these conditions as Tg, Tm, or DTT (**Figure S1**, whole cell lysate IB: KDEL, lanes 3–5 *vs* 2). Hence, we performed titrations to determine optimized conditions for UPR induction (**Figure S2**). 2-DG did not effectively induce BiP upregulation at any concentration in these cells, leading us to exclude this stressor in future experiments. We also observed that DTT, 2-DG, and higher concentrations of BFA inhibited total peroxidase labeling yield. For BFA, we chose the minimum concentration that still yields maximum BiP expression. For DTT-treated cells, ^cyt^APEX labeling can be rescued by aspirating DTT-containing media and replacement of fresh media containing 1 mM H_2_O_2_ (**Figures S3**, whole cell lysate, ECL: biotin). We repeated the treatments with the optimized conditions for each stressor, but still found that only Tg-treated cells display increased ^FLAG^TTR mistargeting in the cytosol (**Figure S3**, avidin purification, IB: TTR). It is difficult to evaluate the extent to which conditions affect TTR mistargeting, however, because mistargeted populations are small and IB bands after proximity labeling are often faint and difficult to quantify (**Figure S1, S3**). Firm conclusions would require substantial material scale-up, and there are many chemical and genetic ER stressors worth considering, especially if pre-QC activation is dependent on how stress is induced. This limitation made us consider using a more sensitive platform for quantifying mistargeted proteins.

### Development of the PRM assay

Peroxidase proximity labeling has been previously integrated with mass spectrometry, with quantification by SILAC^28,47–50^, TMT^51–53^, or label-free methods including MRM and PRM^54–59^. We decided to replace IB with PRM as our detection methods, as shown in **Figure 1**. In addition to ^FLAG^TTR, we included peptides from the labeling peroxidase ^cyt^APEX. Immunodetection of ^cyt^APEX in avidin-enriched samples is difficult, since common epitopes including the FLAG tag are disrupted by proximity labeling, presumably due to direct modification. We considered endogenously biotin-binding proteins in mitochondrial matrix^52,60–62^ as potential global standards for data normalization. These include PC, PCCA, MCCC1 and ACACB. We also targeted common loading control proteins β-actin, α-tubulin and GAPDH. Stress-inducible chaperones HSPA1A (nucleocytosolic) and BiP (*alias* HSPA5/GRP78, primarily ER luminal) were also included.

The workflow of seeking target peptides is summarized in **Figure 2**. In general, we required that peptides (i) be 8-25 amino acids long, (ii) do not contain labile modification sites, (iii) do not contain ragged ends of tryptic digestion,^63^ and (iv) be available in NIST peptide tandem mass spectra library^64^. For ^cyt^APEX peptides and some of chicken avidin peptides, we turned to Prosit^65^ to predict their CID fragmentation patterns at NCE 35%. Uniqueness was examined by either using a background proteome in the Skyline software^37^, or by uploading the candidate precursor list into Nextprot^66^. For actin and tubulin, uniqueness was required at the class-level and not the family-level. Retention times for candidate peptides from ^FLAG^TTR, ^cyt^APEX, actin, tubulin, GAPDH, HSPA1A and BiP were determined by unscheduled analysis of lysate digest, followed by scheduled analysis to evaluate peak shapes. For ^cyt^APEX, actin and tubulin peptides, precursor ion chromatograms are used for quantification because little interference is observed. Mitochondrial matrix biotin carboxylases are at low levels in HEK293T lysate and chicken avidin should not be present in HEK293T lysate. Hence, their peptides were first evaluated from an avidin-purification sample digest in data dependent acquisition mode with the same LC gradient. Only PC peptides, as opposed to those from PCCA, MCCC1 and ACACB, were used because they are the most abundant among the mitochondrial biotin carboxylases. Peptides with proper retention time, high intensity, good peak shape, little transition interference were confirmed with scheduled PRM. We also removed peptides that were confirmed to be deamidation-prone.

**Figure 2.**
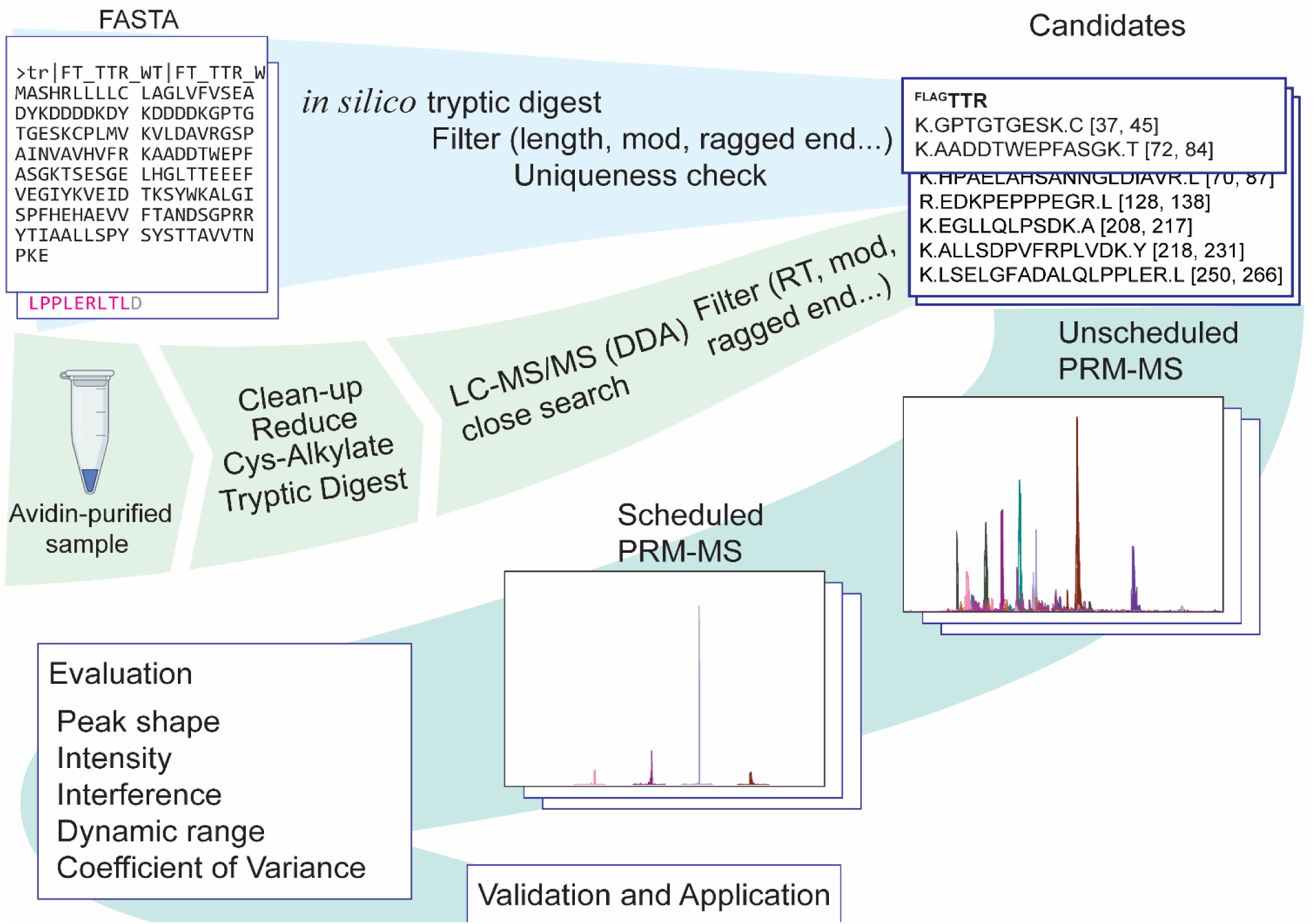
Workflow of choosing peptides for PRM assay. For the generation of the initial target list, protein sequences were subjected to in silico tryptic digestion (C-term to K/R, not before P) and filtered in Skyline. For proteins that are at low abundance in lysate, namely mitochondrial biotin carboxylases, avidin-enriched samples were prepared for liquid chromatography - tandem mass spectrometry (LC-MS/MS) data dependent acquisition (DDA) mode. Raw data was searched with MSFragger^39,40^. Fully tryptic precursor ions with decent intensity, proper retention time, and no sub-stochiometric modification sites and ragged ends are kept. Filtered candidates were confirmed with unscheduled PRM-MS first, retention times of those were confirmed with scheduled PRM-MS. Peak shapes and other properties of selected precursors were evaluated, before or during application.

The targeted peptides in this study are summarized in **Table 1**, with coefficients of variance (*CV*s) of 8 technical replicates listed as well. *CV*s calculated from raw peak areas are below 20%, with a median of 9.1%. If raw peak areas are normalized by total ion current (TIC), *CV*s of assayed peptides do not exceed 14%, with a median of 5.8% (**Figure S4**). The MS^2^ spectra and product ion chromatograms of targeted peptides are provide in **Figure 5**.

**Table 1.**
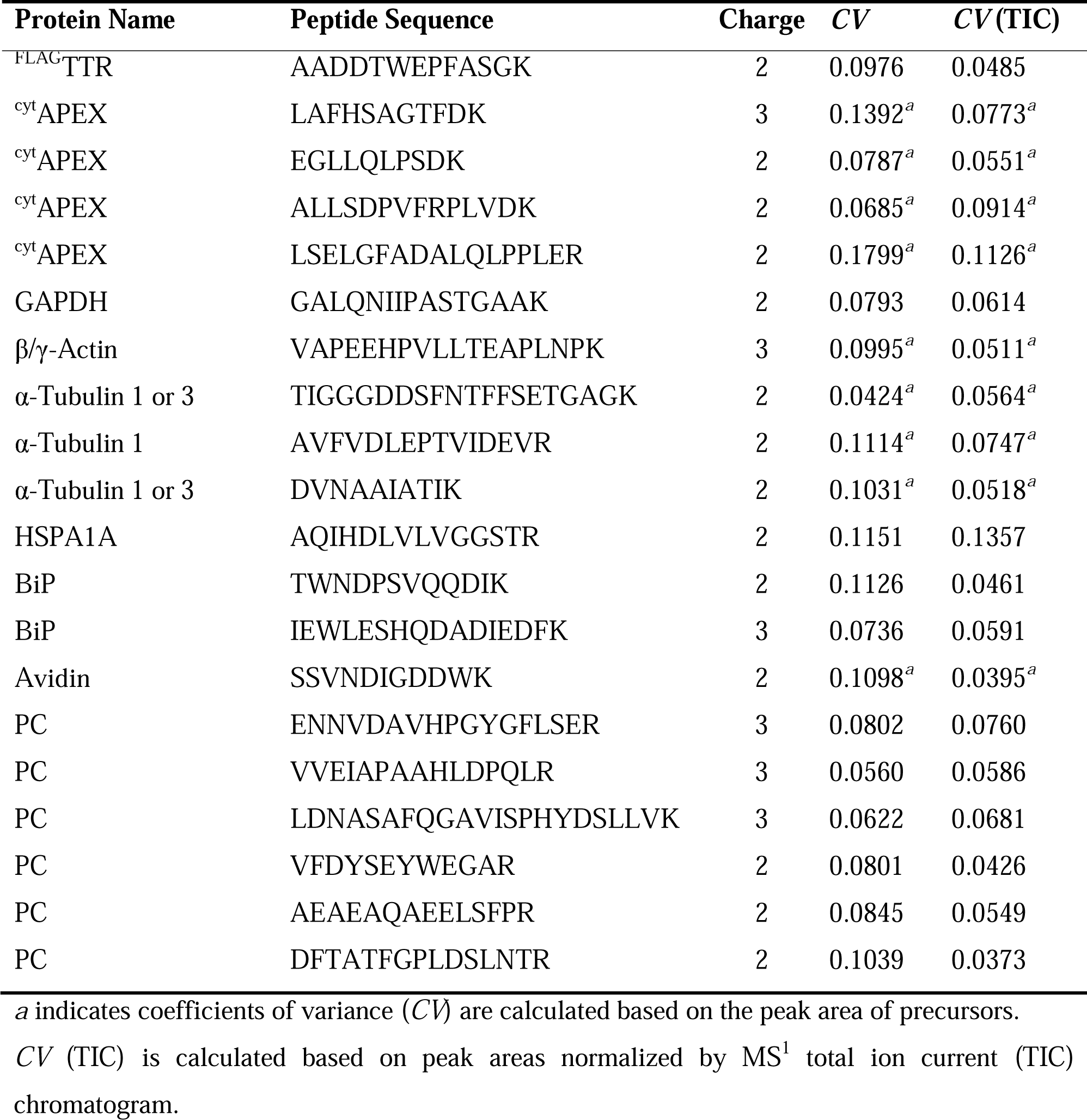
Assayed proteins, peptides, and coefficients of variance (*n* = 8).

### Validation of proximity-labeling PRM Assay

We tested whether IB and PRM yield comparable results with our assay. To perform the comparison, we used drug treatment conditions that provide a large dynamic range of ^FLAG^TTR mistargeting yields. Mycolactone A/B (ML) is an inhibitor of Sec61-mediated co-translational translocation for secreted and type-I and type-II transmembrane proteins^67–69^. It should completely arrest ^FLAG^TTR translocation during the time-course of the experiment. When used in combination with the proteasome inhibitor MG132, ML is expected to give us the most ^FLAG^TTR accumulation inside the cytosol. Tg is a noncompetitive SERCA inhibitor that rapidly induces severe ER stress^70^, and is a known inducer of ER pre-QC^11,12,9^. From our previous study by IB, combined Tg and MG132 treatment triggers around a 3-fold increase in ^FLAG^TTR mistargeting compared to MG132 treatment alone, and 6-fold increase compared to the basal condition.^25^

HEK293T cells co-expressing ^FLAG^TTR and ^cyt^APEX were treated with vehicle, ML or Tg for 16 hours in the absence or presence of 16-h MG132 treatment, before BP-labeling and quenching. Eluate samples (avidin-purifications) were split in half. One half was separated by SDS-PAGE followed by IB (**Figure 3a**), while the other half was prepared for bottom-up proteomics and analyzed by PRM. The amount of eluate digest we injected is equivalent to *one tenth* the amount we recovered from avidin beads. IB is quantified by densitometry, and PRM by raw peak areas of the TTR peptide AADDTWEGFASGK^2+^. Each sample was normalized to the total intensity across conditions for a given replicate.^71^ Tg induces a 2.5-fold increase over vehicle in ^FLAG^TTR mistargeting when co-treated with MG132 (**Figure 3b,c**), which agrees with what we have measured by IB in our previous study^25^. The results from the PRM assay are similar, with Pearson’s *R*^2^ ∼0.99 for both replicates. These experiments demonstrate that PRM yields similar results to IB, but with 10-fold less sample consumption.

**Figure 3.**
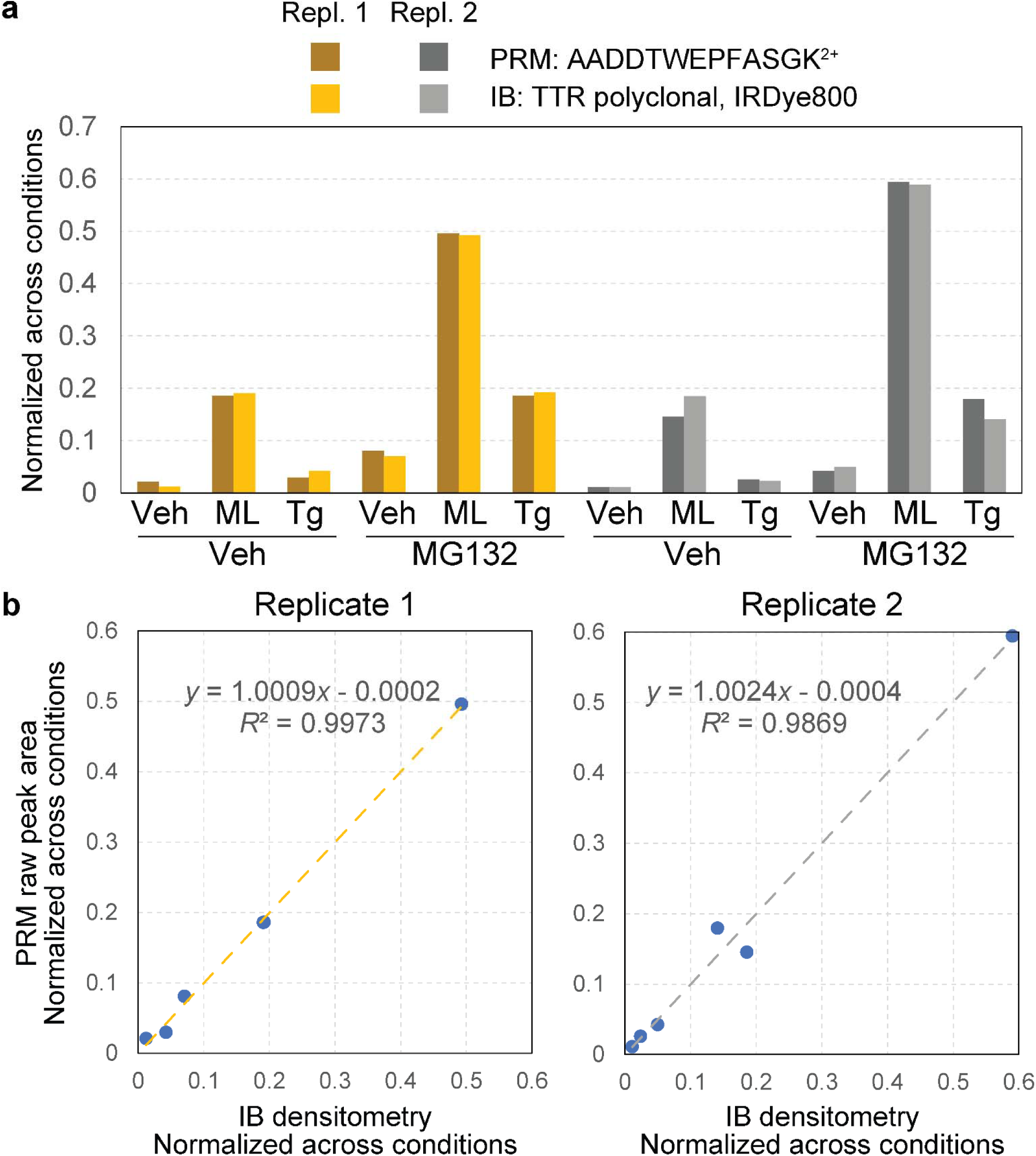
PRM and IB similarly quantify relative ^FLAG^TTR mistargeting. **a)** Quantification of ^FLAG^TTR from avidin-purified samples by IB and PRM. *y*-Axis represents signals normalized across all six conditions within a replicate. Normalization is performed by dividing signal from one condition by tallied signals across all six conditions in a replicate. For IB, it is local background-subtracted band densitometry from each condition normalized to the sum across all conditions; for PRM, it is raw peak area from each condition normalized to the sum across all conditions. See Figure **S6** for full blots of both replicates. **c**) Plot of PRM quantification against IB is displayed to show correlation between the two methods, within the same scale. Linear regression equation and Pearson’s *R*^2^ displayed as well.

### ER pre-QC induction is indeed ER stressor-dependent

With this PRM-coupled mistargeting assay, we revisited the small molecule ER stressors Tg, Tm, BFA and DTT (raw peak areas in **Table S2**). Having observed that not all ER stressors increase mistargeting of ^FLAG^TTR in HEK293T cells, we also included two other molecules that impact ER calcium homeostasis. Cyclopiazonic acid (CPA) is another SERCA inhibitor, but differs from thapsigargin in that it is a competitive inhibitor, less potent, and inhibits SERCA reversibly^72^. Diltiazem (Dil.) is a calcium channel blocker that is used to maintain ER calcium level by preventing Ca^2+^ leakage. Dil. does not induce ER stress, but does influence the ER protein homeostasis through elevated activity of ER calcium binding proteins^73,74^. We confirmed that 100 µM CPA induces ER stress similarly to 50 nM Tg on the basis of BiP upregulation following a 16-h treatment (**Figure S7a**, IB:KDEL, lanes 3,4 *vs* 1,2 and 7,8 *vs* 5,6).

We also took advantage of the inherent multiplexing of PRM to consider normalization. Appropriate normalization to control for loading, sample handling, and ionization efficiency is necessary for most biological mass spectrometry techniques. However, biased normalization methods can introduce artifacts into interpretation of results. A straightforward normalization factor is the area under the entire MS^1^ TIC chromatogram^75^. This factor should control for injection efficiency, loss of material during sample preparation, differences in the recovery of cells or protein, errors in protein quantification prior to avidin purification, and to the extent that the signal is dominated by ^cyt^APEX-labeled proteins, the labeling activity of ^cyt^APEX in a given experiment. Mitochondrial carboxylases such as PC, which are endogenously biotinylated, have been used for normalization in proximity labeling experiments when the peroxidases is localized elsewhere than the mitochondria^52,60–62^, controlling for the total amount protein loaded onto (strept)avidin beads. These proteins as normalization factors are valid if the assumptions of consistent expression level and consistent proximity labeling activity are maintained across conditions. Unlike normalization against TIC and PC, normalization against abundant proteins that share a compartment with the peroxidase can control against changes in BP-labeling efficiency.^76^ We considered β-actin, α-tubulin and GAPDH. We found that α-Tubulin 1B levels are affected by cellular stress (*e.g.* **Figure S6a,b,** Lysate IB: α-tubulin) and poor chromatographic performance in our gradient for GAPDH peptides, and hence focused on β/γ-actin as a proxy for protein load and BP-labeling yield by ^cyt^APEX. We also considered normalization against the heme peroxidase ^cyt^APEX itself, under the expectation that ^cyt^APEX auto-labeling is a proxy for total biotinylation yield.

We compared relative ^FLAG^TTR mistargeting across drug treatments and different normalization schemes (**Figure 4**). Similar results are seen for most treatments. The SERCA inhibitors Tg and CPA induce pre-QC to similar extents. Tm and Dil do not induce pre-QC. The observed relative mistargeting of ^FLAG^TTR following ML treatment varies between normalization methods. While DTT lowers the apparent mistargeted TTR load with each normalization, the extent of this decrease varies from 86% with PC normalization (**Figure 4b**) to 40% with β-actin or ^cyt^APEX normalization (**Figure 4c,d**). BFA shows the largest disparity, doubling ^FLAG^TTR mistargeting with β-actin or ^cyt^APEX normalization, moderately increasing (14% increase) with TIC normalization, or having no effect with PC normalization.

**Figure 4.**
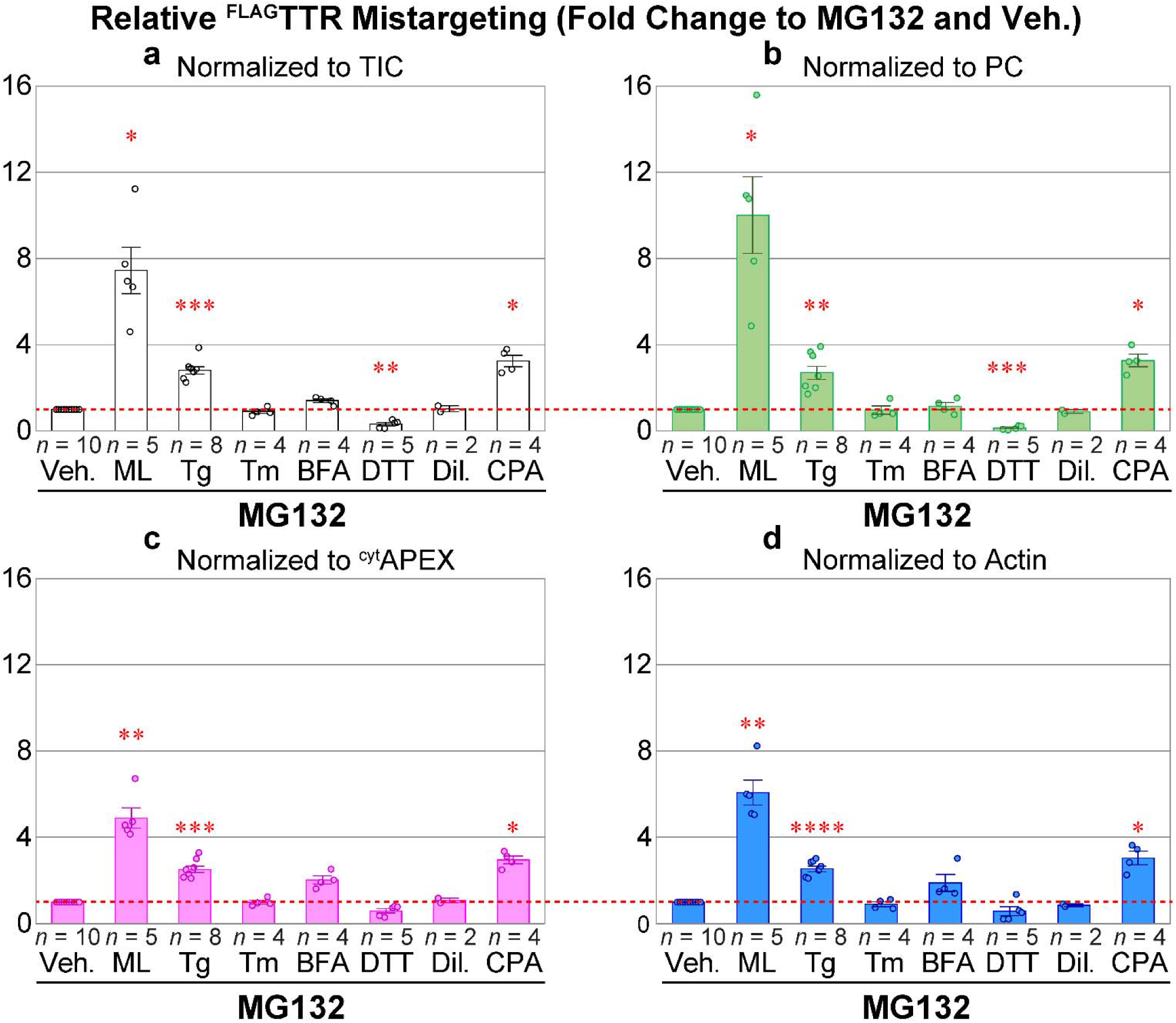
PRM quantification of apparent ^FLAG^TTR mistargeting as results of various drug treatments, in the presence of MG132 (16 h). **a**) MS^1^ TIC-, **b**) PC-, **c**) APEX2- and **d**) actin-normalized PRM peak area fold change, compared to 1 μM MG132 and vehicle. Veh., 0.1% DMSO; ML, 25 nM, mycolactone A/B; Tg, 50 nM thapsigargin; Tm, 200 nM tunicamycin; BFA, 400 ng mL^−1^ Brefeldin A; DTT, 3 mM (*d*,*l*-)1,4-dithiothreitol; Dil., 30 μM diltiazem; CPA, 100 μM cyclopiazonic acid. Representative full blots of lysates in one experiment can be found in **Figure S7b**. Error bars represent standard errors of the mean. Sample sizes are displayed above drug annotations. Fold change of 1 is marked as a red dashed line. Two-tailed heteroscedastic *t*-tests were performed for mistargeting fold change by each normalization factor, with Bonferroni correction (6 comparisons *vs* Veh., excluding Dil.). Adjusted *p*-value ≥ 0.05 or not compared, not annotated; 0.01 ≤ adj. *p* < 0.05, *; 0.001 ≤ adj. *p* < 0.01, **; 0.0001 ≤ adj. *p* < 0.01, ***; adj. *p* < 0.0001, ****.

### Determination of Appropriate Normalization

To explain the disagreement among normalization methods for ML-, DTT- and BFA-treated cells, we considered how each of these will be affected by changes to labeling efficiency. PC recovery from avidin purification will solely reflect the total amount of lysate added to the beads. It will be insensitive to changes in peroxidase labeling. Recovery of biotinylated ^cyt^APEX or β-actin, by contrast, will reflect both the amount of cells harvested as well as the peroxidase labeling efficiency. Normalization against MS^1^ TIC will also partially account for differences in labeling efficiency, however several other factors will affect the TIC signal (**Figure 5a**). These include carboxylases such as PC, which reflect total protein inputs, but also common contaminants (keratin, trypsin, etc.) and avidin that can be leached from the beads in a strongly condition- and lot-dependent manner^77^.

**Figure 5.**
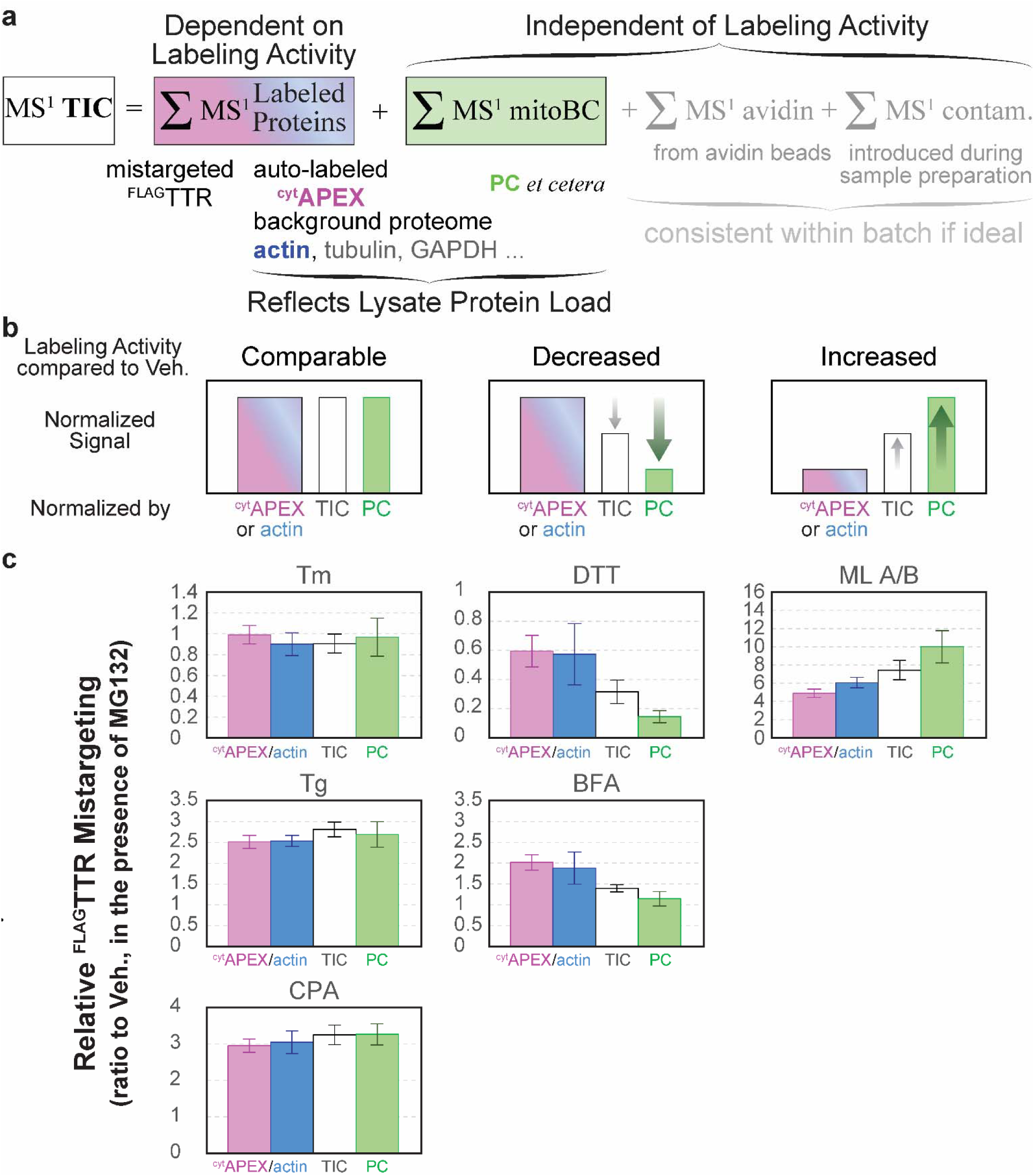
Composition of MS^1^ TIC chromatogram and the effect of labeling activity on the pattern of normalized data, organized per drug treatment. **a**) MS^1^ TIC consists of the MS^1^ chromatograms of peptides from BP-labeled proteins, endogenous biotin carboxylases in mitochondrial matrix (mitoBC, independent of labeling), avidin etched from beads during elution and denaturation (independent of labeling), and other contaminants (independent of labeling). **b**) Prediction of how relative amount of BP-labeling yield and PC level impacts normalized ^FLAG^TTR mistargeting. For treatments that neither alter cytoplasmic proteome too much, nor change the relative levels of ^cyt^APEX, actin and PC, normalization to ^cyt^APEX, actin, TIC and PC gives similar results. For conditions that decrease the labeling activity, fold change normalized to ^cyt^APEX or actin will be higher than that by PC. For conditions that increase the relative level of labeling, fold change normalized to ^cyt^APEX or actin will be lower than that by PC. **c**) Head-to-head comparison between apparent ^FLAG^TTR mistargeting normalized by different factors, per drug treatment. These drug treatment conditions are categorized based on the pattern in panel **b**. All drug treatment conditions include 16-h 1 μM MG132. Error bars represent the standard errors of the mean.

Prior to the affinity purification step, we bring clarified lysates to the same protein concentration, based on Bradford assay, and load the same mass of protein to avidin-agarose beads. If the drug treatments, as compared to vehicle, do not change the profile of labeling-independent biotin carboxylases, background cytoplasmic proteome and the labeling activity of ^cyt^APEX, normalization to different global standards should yield similar results (**Figure 5b** left, “comparable labeling activity”). If ^cyt^APEX labeling activity somehow decreases under some drug treatment conditions, but the total amount of biotin carboxylases and cytoplasmic background remain consistent, the portion of enriched cytoplasmic protein (*e.g.* ^cyt^APEX and β-actin) is expected to decrease accordingly with the labeling activity decrease. This may result in a relatively higher proportion of PC and other labeling-independent components. Eventually, if data are divided by the level of ^cyt^APEX or actin, the normalized value will be higher than that normalized by PC. On the contrary, data divided by PC level will be smaller than that by ^cyt^APEX or actin. MS^1^ TIC can represent labeling activity to some extent, but it is also convoluted by labeling-independent components, thus mistargeting normalized by TIC is expected to be in the middle of the two extremes (**Figure 5b** center, “decreased labeling activity”). And *vice versa*, if ^cyt^APEX labeling activity increases upon some treatments compared to the control condition, an opposite trend will be expected (**Figure 5b** right, “increased labeling activity”): the proportion of avidin-purified cytoplasmic proteins (*e.g.* ^cyt^APEX or actin) will be relatively higher, resulting in a lower apparent mistargeting after normalization, while the proportion of labeling-independent PC will be lower, leading to a higher apparent mistargeting after normalization. Again, TIC normalization should yield an intermediate result. We find that our data nicely matches this model (**Figure 5C**), indicating that the divergence between TIC- and PC-normalization and ^cyt^APEX-normalization can be entirely ascribed to drug treatment dependent variation in peroxidase labeling efficiency. We saw in our immunoblotting experiments (**Figure S1**) that BFA and DTT decrease labeling efficiency, and altered our protocol to mitigate this interference. Nevertheless, it is clear that these treatments even under optimized conditions affect peroxidase labeling enough to influence the quantitative accuracy of the data. Given that DTT is a potent reductant, it is not surprising that it inhibits oxidative labeling. The cause of inhibition during BFA treatment is unclear. It could be that changes in glutathione redox state also mediate the increased labeling following ML treatment, as a recent study shows that ML depletes cellular glutathione in myeloid leukemia cells KBM-7.78 Whatever the basis of the change in labeling efficiency, by using PRM, we can normalize against ^cyt^APEX auto-labeling and remove this confounding factor to find the most accurate quantification of mistargeting (**Figure 4C**)

## CONCLUSIONS

In this study, we coupled PRM mass spectrometry with our protein mistargeting assay, enabling us to control for labeling efficiency and quantitatively compare the extent to which several ER stressors induce TTR mistargeting through ER pre-QC. Our results indicate that UPR activation alone is not sufficient to induce ER pre-QC in HEK293T cells. A previous study of prion protein (PrP) mistargeting in HeLa cells found that Tg and DTT, but not BFA, treatment induce pre-QC of PrP^79^. Another study found that both Tg and Tm induce ER pre-QC in HepG2 cells^12^. Our PRM-coupled mistargeting assay will now enable the effects of ER stress inducers and other proteostasis regulators to be scanned across multiple cell types, towards establishing the generality of ER pre-QC induction by ER stress. This platform could also be used to evaluate other pre-QC substrates beyond TTR, or to determine which signaling factors participate in mediating pre-QC. More broadly, we have demonstrated that the multiplexing capacity of PRM can be leveraged to ensure appropriate normalization when using in situ peroxidase labeling

## ASSOCIATED CONTENT

### Supporting Information

Supporting information (pdf) includes Table S1, description of drug treatment conditions, Figures including whole blots, extracted ion chromatograms, and mass spectra.

**Table S2** raw and normalized PRM peak areas, quantification and statistics (XLSX)

## AUTHOR INFORMATION

### Corresponding Author

**Joseph C. Genereux** – *Department of Chemistry, University of California, Riverside, California 92521, United States*; orcid.org/0000-0002-5093-7710; phone: 1-951-827-3759;

### Author

***Ziqi Lyu*** *– Department of Chemistry, University of California, Riverside, California 92521, United States*.

### Author Contributions

The manuscript was written through contributions of all authors. All authors have given approval to the final version of the manuscript.

### Notes

The authors declare no competing financial interest.

## ACKNOWLEDGMENT

We gratefully acknowledge Y. Kishi for the generous gift of mycolactone A/B. Support was provided by the University of California and NIH R01GM134125. We acknowledge the Analytical Chemistry Instrumentation Facility instrumentation, supported by the UC-Riverside Chemistry ACIF fund. Figures were created with Biorender.com.

## REFERENCE

1. Juszkiewicz, S.; Hegde, R. S. Quality Control of Orphaned Proteins. Molecular Cell 2018, 71 (3), 443–457. https://doi.org/10.1016/j.molcel.2018.07.001.

2. Carreras-Sureda, A.; Pihán, P.; Hetz, C. Calcium Signaling at the Endoplasmic Reticulum: Fine-Tuning Stress Responses. Cell Calcium 2018, 70, 24–31. https://doi.org/10.1016/j.ceca.2017.08.004.

3. Walczak, C. P.; Bernardi, K. M.; Tsai, B. Endoplasmic Reticulum-Dependent Redox Reactions Control Endoplasmic Reticulum-Associated Degradation and Pathogen Entry. Antioxidants & Redox Signaling 2012, 16 (8), 809–818. https://doi.org/10.1089/ars.2011.4425.

4. Kozlov, G.; Gehring, K. Calnexin Cycle – Structural Features of the ER Chaperone System. FEBS J. 2020, 287 (20), 4322–4340. https://doi.org/10.1111/febs.15330.

5. Rane, N. S.; Kang, S.-W.; Chakrabarti, O.; Feigenbaum, L.; Hegde, R. S. Reduced Translocation of Nascent Prion Protein During ER Stress Contributes to Neurodegeneration. Developmental Cell 2008, 15 (3), 359–370. https://doi.org/10.1016/j.devcel.2008.06.015.

6. Zhang, X.; Shan, S. O. Fidelity of Cotranslational Protein Targeting by the Signal Recognition Particle. Annu Rev Biophys 2014, 43, 381–408.

7. Rodrigo-Brenni, M. C.; Gutierrez, E.; Hegde, R. S. Cytosolic Quality Control of Mislocalized Proteins Requires RNF126 Recruitment to Bag6. Mol Cell 2014, 55 (2), 227–237.

8. Hessa, T.; Sharma, A.; Mariappan, M.; Eshleman, H. D.; Gutierrez, E.; Hegde, R. S. Protein Targeting and Degradation Are Coupled for Elimination of Mislocalized Proteins. Nature 2011, 475 (7356), 394–397.

9. Kadowaki, H.; Satrimafitrah, P.; Takami, Y.; Nishitoh, H. Molecular Mechanism of ER Stress-Induced Pre-Emptive Quality Control Involving Association of the Translocon, Derlin-1, and HRD1. Sci Rep 2018, 8 (1), 1–11. https://doi.org/10.1038/s41598-018-25724-x.

10. Braunstein, I.; Zach, L.; Allan, S.; Kalies, K. U.; Stanhill, A. Proteasomal Degradation of Preemptive Quality Control (PQC) Substrates Is Mediated by an AIRAPL-P97 Complex. Mol Biol Cell 2015, 26 (21), 3719–3727.

11. Kang, S. W.; Rane, N. S.; Kim, S. J.; Garrison, J. L.; Taunton, J.; Hegde, R. S. Substrate-Specific Translocational Attenuation during ER Stress Defines a Pre-Emptive Quality Control Pathway. Cell 2006, 127 (5), 999–1013.

12. Kadowaki, H.; Nagai, A.; Maruyama, T.; Takami, Y.; Satrimafitrah, P.; Kato, H.; Honda, A.; Hatta, T.; Natsume, T.; Sato, T.; Kai, H.; Ichijo, H.; Nishitoh, H. Pre-Emptive Quality Control Protects the ER from Protein Overload via the Proximity of ERAD Components and SRP. Cell Reports 2015, 13 (5), 944–956. https://doi.org/10.1016/j.celrep.2015.09.047.

13. McCaffrey, K.; Braakman, I. Protein Quality Control at the Endoplasmic Reticulum. Essays in Biochemistry 2016, 60 (2), 227–235. https://doi.org/10.1042/EBC20160003.

14. Arrieta, A.; Blackwood, E. A.; Glembotski, C. C. ER Protein Quality Control and the Unfolded Protein Response in the Heart. In Coordinating Organismal Physiology Through the Unfolded Protein Response; Wiseman, R. L., Haynes, C. M., Eds.; Current Topics in Microbiology and Immunology; Springer International Publishing: Cham, 2017; Vol. 414, pp 193–213. https://doi.org/10.1007/82_2017_54.

15. Schwabl, S.; Teis, D. Protein Quality Control at the Golgi. Current Opinion in Cell Biology 2022, 75, 102074. https://doi.org/10.1016/j.ceb.2022.02.008.

16. Lyu, Z.; Genereux, J. C. Methodologies for Measuring Protein Trafficking across Cellular Membranes. ChemPlusChem 2021, 86 (10), 1397–1415. https://doi.org/10.1002/cplu.202100304.

17. Sharma, A.; Mariappan, M.; Appathurai, S.; Hegde, R. S. In Vitro Dissection of Protein Translocation into the Mammalian Endoplasmic Reticulum. In Protein Secretion: Methods and Protocols; Economou, A., Economou, A., Eds.; Totowa, NJ, 2010; pp 339–363.

18. Kadowaki, H.; Nishitoh, H. Endoplasmic Reticulum Quality Control by Garbage Disposal. FEBS J 2019, 286 (2), 232–240. https://doi.org/10.1111/febs.14589.

19. Bosch, J. A.; Chen, C.-L.; Perrimon, N. Proximity-Dependent Labeling Methods for Proteomic Profiling in Living Cells: An Update. WIREs Developmental Biology 2021, 10 (1), e392. https://doi.org/10.1002/wdev.392.

20. Droujinine, I. A.; Meyer, A. S.; Wang, D.; Udeshi, N. D.; Hu, Y.; Rocco, D.; McMahon, J. A.; Yang, R.; Guo, J.; Mu, L.; Carey, D. K.; Svinkina, T.; Zeng, R.; Branon, T.; Tabatabai, A.; Bosch, J. A.; Asara, J. M.; Ting, A. Y.; Carr, S. A.; McMahon, A. P.; Perrimon, N. Proteomics of Protein Trafficking by in Vivo Tissue-Specific Labeling. Nature Communications 2021, 12 (1), 2382. https://doi.org/10.1038/s41467-021-22599-x.

21. Liu, J.; Jang, J. Y.; Pirooznia, M.; Liu, S.; Finkel, T. The Secretome Mouse Provides a Genetic Platform to Delineate Tissue-Specific in Vivo Secretion. Proc. Natl. Acad. Sci. U.S.A. 2021, 118 (3), e2005134118. https://doi.org/10.1073/pnas.2005134118.

22. Kim, K.; Park, I.; Kim, J.; Kang, M.-G.; Choi, W. G.; Shin, H.; Kim, J.-S.; Rhee, H.-W.; Suh, J. M. Dynamic Tracking and Identification of Tissue-Specific Secretory Proteins in the Circulation of Live Mice. Nat Commun 2021, 12 (1), 1–9. https://doi.org/10.1038/s41467-021-25546-y.

23. Lee, S.-Y.; Cheah, J. S.; Zhao, B.; Xu, C.; Roh, H.; Kim, C. K.; Cho, K. F.; Udeshi, N. D.; Carr, S. A.; Ting, A. Y. Engineered Allostery in Light-Regulated LOV-Turbo Enables Precise Spatiotemporal Control of Proximity Labeling in Living Cells. Nat Methods 2023, 20 (6), 908– 917. https://doi.org/10.1038/s41592-023-01880-5.

24. Qin, W.; Cheah, J. S.; Xu, C.; Messing, J.; Freibaum, B. D.; Boeynaems, S.; Taylor, J. P.; Udeshi, N. D.; Carr, S. A.; Ting, A. Y. Dynamic Mapping of Proteome Trafficking within and between Living Cells by TransitID. Cell 2023, S0092867423005962. https://doi.org/10.1016/j.cell.2023.05.044.

25. Lyu, Z.; Sycks, M. M.; Espinoza, M. F.; Nguyen, K. K.; Montoya, M. R.; Galapate, C. M.; Mei, L.; Genereux, J. C. Monitoring Protein Import into the Endoplasmic Reticulum in Living Cells with Proximity Labeling. ACS Chem. Biol. 2022, 17 (7), 1963–1977. https://doi.org/10.1021/acschembio.2c00405.

26. Espinoza, M. F.; Nguyen, K. K.; Sycks, M. M.; Lyu, Z.; Quanrud, G. M.; Montoya, M. R.; Genereux, J. C. Heat Shock Protein Hspa13 Regulates Endoplasmic Reticulum and Cytosolic Proteostasis through Modulation of Protein Translocation. Journal of Biological Chemistry 2022, 298 (12), 102597. https://doi.org/10.1016/j.jbc.2022.102597.

27. Lam, S. S.; Martell, J. D.; Kamer, K. J.; Deerinck, T. J.; Ellisman, M. H.; Mootha, V. K.; Ting, A. Y. Directed Evolution of APEX2 for Electron Microscopy and Proximity Labeling. Nat Methods 2015, 12 (1), 51–54.

28. Lee, S. Y.; Kang, M. G.; Park, J. S.; Lee, G.; Ting, A. Y.; Rhee, H. W. APEX Fingerprinting Reveals the Subcellular Localization of Proteins of Interest. Cell Rep 2016, 15 (8), 1837–1847.

29. Martell, J. D.; Deerinck, T. J.; Lam, S. S.; Ellisman, M. H.; Ting, A. Y. Electron Microscopy Using the Genetically Encoded APEX2 Tag in Cultured Mammalian Cells. Nat Protoc 2017, 12 (9), 1792–1816. https://doi.org/10.1038/nprot.2017.065.

30. Peterson, A. C.; Russell, J. D.; Bailey, D. J.; Westphall, M. S.; Coon, J. J. Parallel Reaction Monitoring for High Resolution and High Mass Accuracy Quantitative, Targeted Proteomics. Molecular & Cellular Proteomics 2012, 11 (11), 1475–1488. https://doi.org/10.1074/mcp.O112.020131.

31. Aebersold, R.; Burlingame, A. L.; Bradshaw, R. A. Western Blots versus Selected Reaction Monitoring Assays: Time to Turn the Tables? Molecular & Cellular Proteomics 2013, 12 (9), 2381–2382. https://doi.org/10.1074/mcp.E113.031658.

32. Van Bentum, M.; Selbach, M. An Introduction to Advanced Targeted Acquisition Methods. Molecular & Cellular Proteomics 2021, 20, 100165. https://doi.org/10.1016/j.mcpro.2021.100165.

33. Heil, L. R.; Remes, P. M.; MacCoss, M. J. Comparison of Unit Resolution Versus High-Resolution Accurate Mass for Parallel Reaction Monitoring. J. Proteome Res. 2021, 20 (9), 4435– 4442. https://doi.org/10.1021/acs.jproteome.1c00377.

34. Pino, L. K.; Searle, B. C.; Bollinger, J. G.; Nunn, B.; MacLean, B.; MacCoss, M. J. The Skyline Ecosystem: Informatics for Quantitative Mass Spectrometry Proteomics. Mass Spec Rev 2020, 39 (3), 229–244. https://doi.org/10.1002/mas.21540.

35. Carr, S. A.; Abbatiello, S. E.; Ackermann, B. L.; Borchers, C.; Domon, B.; Deutsch, E. W.; Grant, R. P.; Hoofnagle, A. N.; Hüttenhain, R.; Koomen, J. M.; Liebler, D. C.; Liu, T.; MacLean, B.; Mani, D.; Mansfield, E.; Neubert, H.; Paulovich, A. G.; Reiter, L.; Vitek, O.; Aebersold, R.; Anderson, L.; Bethem, R.; Blonder, J.; Boja, E.; Botelho, J.; Boyne, M.; Bradshaw, R. A.; Burlingame, A. L.; Chan, D.; Keshishian, H.; Kuhn, E.; Kinsinger, C.; Lee, J. S. H.; Lee, S.-W.; Moritz, R.; Oses-Prieto, J.; Rifai, N.; Ritchie, J.; Rodriguez, H.; Srinivas, P. R.; Townsend, R. R.; Van Eyk, J.; Whiteley, G.; Wiita, A.; Weintraub, S. Targeted Peptide Measurements in Biology and Medicine: Best Practices for Mass Spectrometry-Based Assay Development Using a Fit-for-Purpose Approach. Molecular & Cellular Proteomics 2014, 13 (3), 907–917. https://doi.org/10.1074/mcp.M113.036095.

36. Kennedy, J. J.; Whiteaker, J. R.; Ivey, R. G.; Burian, A.; Chowdhury, S.; Tsai, C.-F.; Liu, T.; Lin, C.; Murillo, O. D.; Lundeen, R. A.; Jones, L. A.; Gafken, P. R.; Longton, G.; Rodland, K. D.; Skates, S. J.; Landua, J.; Wang, P.; Lewis, M. T.; Paulovich, A. G. Internal Standard Triggered-Parallel Reaction Monitoring Mass Spectrometry Enables Multiplexed Quantification of Candidate Biomarkers in Plasma. Anal. Chem. 2022, 94 (27), 9540–9547. https://doi.org/10.1021/acs.analchem.1c04382.

37. MacLean, B.; Tomazela, D. M.; Shulman, N.; Chambers, M.; Finney, G. L.; Frewen, B.; Kern, R.; Tabb, D. L.; Liebler, D. C.; MacCoss, M. J. Skyline: An Open Source Document Editor for Creating and Analyzing Targeted Proteomics Experiments. Bioinformatics 2010, 26 (7), 966–968. https://doi.org/10.1093/bioinformatics/btq054.

38. Sharma, V.; Eckels, J.; Schilling, B.; Ludwig, C.; Jaffe, J. D.; MacCoss, M. J.; MacLean, B. Panorama Public: A Public Repository for Quantitative Data Sets Processed in Skyline. Molecular & Cellular Proteomics 2018, 17 (6), 1239–1244. https://doi.org/10.1074/mcp.RA117.000543.

39. Kong, A. T.; Leprevost, F. V.; Avtonomov, D. M.; Mellacheruvu, D.; Nesvizhskii, A. I. MSFragger: Ultrafast and Comprehensive Peptide Identification in Mass Spectrometry–Based Proteomics. Nat Methods 2017, 14 (5), 513–520. https://doi.org/10.1038/nmeth.4256.

40. Teo, G. C.; Polasky, D. A.; Yu, F.; Nesvizhskii, A. I. Fast Deisotoping Algorithm and Its Implementation in the MSFragger Search Engine. J. Proteome Res. 2021, 20 (1), 498–505. https://doi.org/10.1021/acs.jproteome.0c00544.

41. Legesse, A.; Kaushansky, N.; Braunstein, I.; Saad, H.; Lederkremer, G.; Navon, A.; Stanhill, A. The Role of RNF149 in the Pre-Emptive Quality Control Substrate Ubiquitination. Commun Biol 2023, 6 (1), 385. https://doi.org/10.1038/s42003-023-04763-9.

42. Yoo, J.; Mashalidis, E. H.; Kuk, A. C. Y.; Yamamoto, K.; Kaeser, B.; Ichikawa, S.; Lee, S.-Y. GlcNAc-1-P-Transferase–Tunicamycin Complex Structure Reveals Basis for Inhibition of N-Glycosylation. Nat Struct Mol Biol 2018, 25 (3), 217–224. https://doi.org/10.1038/s41594-018-0031-y.

43. Kurtoglu, M.; Gao, N.; Shang, J.; Maher, J. C.; Lehrman, M. A.; Wangpaichitr, M.; Savaraj, N.; Lane, A. N.; Lampidis, T. J. Under Normoxia, 2-Deoxy-D -Glucose Elicits Cell Death in Select Tumor Types Not by Inhibition of Glycolysis but by Interfering with N-Linked Glycosylation. Molecular Cancer Therapeutics 2007, 6 (11), 3049–3058. https://doi.org/10.1158/1535-7163.MCT-07-0310.

44. Klausner, R. D.; Donaldson, J. G.; Lippincott-Schwartz, J. Brefeldin A: Insights into the Control of Membrane Traffic and Organelle Structure. Journal of Cell Biology 1992, 116 (5), 1071–1080. https://doi.org/10.1083/jcb.116.5.1071.

45. Nebenführ, A.; Ritzenthaler, C.; Robinson, D. G. Brefeldin A: Deciphering an Enigmatic Inhibitor of Secretion. Plant Physiol 2002, 130 (3), 1102–1108. https://doi.org/10.1104/pp.011569.

46. Citterio, C.; Vichi, A.; Pacheco-Rodriguez, G.; Aponte, A. M.; Moss, J.; Vaughan, M. Unfolded Protein Response and Cell Death after Depletion of Brefeldin A-Inhibited Guanine Nucleotide-Exchange Protein GBF1. Proc. Natl. Acad. Sci. U.S.A. 2008, 105 (8), 2877–2882. https://doi.org/10.1073/pnas.0712224105.

47. Rhee, H. W.; Zou, P.; Udeshi, N. D.; Martell, J. D.; Mootha, V. K.; Carr, S. A.; Ting, A. Y. Proteomic Mapping of Mitochondria in Living Cells via Spatially Restricted Enzymatic Tagging. Science 2013, 339 (6125), 1328–1331.

48. James, C.; Müller, M.; Goldberg, M. W.; Lenz, C.; Urlaub, H.; Kehlenbach, R. H. Proteomic Mapping by Rapamycin-Dependent Targeting of APEX2 Identifies Binding Partners of VAPB at the Inner Nuclear Membrane. J Biol Chem 2019, 294 (44), 16241–16254. https://doi.org/10.1074/jbc.RA118.007283.

49. Li, Y.; Tian, C.; Liu, K.; Zhou, Y.; Yang, J.; Zou, P. A Clickable APEX Probe for Proximity-Dependent Proteomic Profiling in Yeast. Cell Chemical Biology 2020, 27 (7), 858–865.e8. https://doi.org/10.1016/j.chembiol.2020.05.006.

50. Bersuker, K.; Peterson, C. W. H.; To, M.; Sahl, S. J.; Savikhin, V.; Grossman, E. A.; Nomura, D. K.; Olzmann, J. A. A Proximity Labeling Strategy Provides Insights into the Composition and Dynamics of Lipid Droplet Proteomes. Developmental Cell 2018, 44 (1), 97–112.e7. https://doi.org/10.1016/j.devcel.2017.11.020.

51. Perez Verdaguer, M.; Zhang, T.; Surve, S.; Paulo, J. A.; Wallace, C.; Watkins, S. C.; Gygi, S. P.; Sorkin, A. Time-Resolved Proximity Labeling of Protein Networks Associated with Ligand-Activated EGFR. Cell Rep 2022, 39 (11), 110950. https://doi.org/10.1016/j.celrep.2022.110950.

52. Paek, J.; Kalocsay, M.; Staus, D. P.; Wingler, L.; Pascolutti, R.; Paulo, J. A.; Gygi, S. P.; Kruse, A. C. Multidimensional Tracking of GPCR Signaling via Peroxidase-Catalyzed Proximity Labeling. Cell 2017, 169 (2), 338–349.e11. https://doi.org/10.1016/j.cell.2017.03.028.

53. Chu, T.-T.; Tu, X.; Yang, K.; Wu, J.; Repa, J. J.; Yan, N. Tonic Prime-Boost of STING Signalling Mediates Niemann–Pick Disease Type C. Nature 2021, 596 (7873), 570–575. https://doi.org/10.1038/s41586-021-03762-2.

54. Lobingier, B. T.; Huttenhain, R.; Eichel, K.; Miller, K. B.; Ting, A. Y.; von Zastrow, M.; Krogan, N. J. An Approach to Spatiotemporally Resolve Protein Interaction Networks in Living Cells. Cell 2017, 169 (2), 350–360 e12.

55. Kong, Q.; Ke, M.; Weng, Y.; Qin, Y.; He, A.; Li, P.; Cai, Z.; Tian, R. Dynamic Phosphotyrosine-Dependent Signaling Profiling in Living Cells by Two-Dimensional Proximity Proteomics. J. Proteome Res. 2022, 21 (11), 2727–2735. https://doi.org/10.1021/acs.jproteome.2c00418.

56. Ke, M.; Yuan, X.; He, A.; Yu, P.; Chen, W.; Shi, Y.; Hunter, T.; Zou, P.; Tian, R. Spatiotemporal Profiling of Cytosolic Signaling Complexes in Living Cells by Selective Proximity Proteomics. Nat Commun 2021, 12 (1), 71. https://doi.org/10.1038/s41467-020-20367-x.

57. Saha, B.; Salemi, M.; Williams, G. L.; Oh, S.; Paffett, M. L.; Phinney, B.; Mandell, M. A. Interactomic Analysis Reveals a Homeostatic Role for the HIV Restriction Factor TRIM5α in Mitophagy. Cell Reports 2022, 39 (6), 110797. https://doi.org/10.1016/j.celrep.2022.110797.

58. Hobson, B. D.; Choi, S. J.; Mosharov, E. V.; Soni, R. K.; Sulzer, D.; Sims, P. A. Subcellular Proteomics of Dopamine Neurons in the Mouse Brain. eLife 2022, 11, e70921. https://doi.org/10.7554/eLife.70921.

59. Zhong, X.; Li, Q.; Polacco, B. J.; Patil, T.; DiBerto, J. F.; Vartak, R.; Xu, J.; Marley, A.; Foussard, H.; Roth, B. L.; Eckhardt, M.; Zastrow, M. V.; Krogan, N. J.; Hüttenhain, R. An Automated Proximity Proteomics Pipeline for Subcellular Proteome and Protein Interaction Mapping; preprint; Systems Biology, 2023. https://doi.org/10.1101/2023.04.11.536358.

60. Xiong, Z.; Lo, H. P.; McMahon, K.-A.; Martel, N.; Jones, A.; Hill, M. M.; Parton, R. G.; Hall, T. E. In Vivo Proteomic Mapping through GFP-Directed Proximity-Dependent Biotin Labelling in Zebrafish. eLife 2021, 10, e64631. https://doi.org/10.7554/eLife.64631.

61. Garcia, Y. A.; Velasquez, E. F.; Gao, L. W.; Gholkar, A. A.; Clutario, K. M.; Cheung, K.; Williams-Hamilton, T.; Whitelegge, J. P.; Torres, J. Z. Mapping Proximity Associations of Core Spindle Assembly Checkpoint Proteins. J. Proteome Res. 2021, 20 (7), 3414–3427. https://doi.org/10.1021/acs.jproteome.0c00941.

62. Frankenfield, A. M.; Fernandopulle, M. S.; Hasan, S.; Ward, M. E.; Hao, L. Development and Comparative Evaluation of Endolysosomal Proximity Labeling-Based Proteomic Methods in Human IPSC-Derived Neurons. Anal. Chem. 2020, 92 (23), 15437–15444. https://doi.org/10.1021/acs.analchem.0c03107.

63. Rauniyar, N. Parallel Reaction Monitoring: A Targeted Experiment Performed Using High Resolution and High Mass Accuracy Mass Spectrometry. IJMS 2015, 16 (12), 28566–28581. https://doi.org/10.3390/ijms161226120.

64. Stein, S. NIST Libraries of Peptide Fragmentation Mass Spectra, NIST Standard Reference Database 1 C, 2008. https://doi.org/10.18434/T4ZK5S.

65. Gessulat, S.; Schmidt, T.; Zolg, D. P.; Samaras, P.; Schnatbaum, K.; Zerweck, J.; Knaute, T.; Rechenberger, J.; Delanghe, B.; Huhmer, A.; Reimer, U.; Ehrlich, H.-C.; Aiche, S.; Kuster, B.; Wilhelm, M. Prosit: Proteome-Wide Prediction of Peptide Tandem Mass Spectra by Deep Learning. Nat Methods 2019, 16 (6), 509–518. https://doi.org/10.1038/s41592-019-0426-7.

66. Schaeffer, M.; Gateau, A.; Teixeira, D.; Michel, P.-A.; Zahn-Zabal, M.; Lane, L. The NeXtProt Peptide Uniqueness Checker: A Tool for the Proteomics Community. Bioinformatics 2017, 33 (21), 3471–3472. https://doi.org/10.1093/bioinformatics/btx318.

67. McKenna, M.; Simmonds, R. E.; High, S. Mechanistic Insights into the Inhibition of Sec61-Dependent Co- and Post-Translational Translocation by Mycolactone. Journal of Cell Science 2016, jcs.182352. https://doi.org/10.1242/jcs.182352.

68. McKenna, M.; Simmonds, R. E.; High, S. Mycolactone Reveals Substrate-Driven Complexity of Sec61-Dependent Transmembrane Protein Biogenesis. Journal of Cell Science 2017, jcs.198655. https://doi.org/10.1242/jcs.198655.

69. Morel, J.-D.; Paatero, A. O.; Wei, J.; Yewdell, J. W.; Guenin-Macé, L.; Van Haver, D.; Impens, F.; Pietrosemoli, N.; Paavilainen, V. O.; Demangel, C. Proteomics Reveals Scope of Mycolactone-Mediated Sec61 Blockade and Distinctive Stress Signature. Mol Cell Proteomics 2018, 17 (9), 1750–1765. https://doi.org/10.1074/mcp.RA118.000824.

70. Sehgal, P.; Szalai, P.; Olesen, C.; Praetorius, H. A.; Nissen, P.; Christensen, S. B.; Engedal, N.; Møller, J. V. Inhibition of the Sarco/Endoplasmic Reticulum (ER) Ca2+-ATPase by Thapsigargin Analogs Induces Cell Death via ER Ca2+ Depletion and the Unfolded Protein Response. Journal of Biological Chemistry 2017, 292 (48), 19656–19673. https://doi.org/10.1074/jbc.M117.796920.

71. Degasperi, A.; Birtwistle, M. R.; Volinsky, N.; Rauch, J.; Kolch, W.; Kholodenko, B. N. Evaluating Strategies to Normalise Biological Replicates of Western Blot Data. PLoS ONE 2014, 9 (1), e87293. https://doi.org/10.1371/journal.pone.0087293.

72. Moncoq, K.; Trieber, C. A.; Young, H. S. The Molecular Basis for Cyclopiazonic Acid Inhibition of the Sarcoplasmic Reticulum Calcium Pump. Journal of Biological Chemistry 2007, 282 (13), 9748–9757. https://doi.org/10.1074/jbc.M611653200.

73. Mu, T.-W.; Fowler, D. M.; Kelly, J. W. Partial Restoration of Mutant Enzyme Homeostasis in Three Distinct Lysosomal Storage Disease Cell Lines by Altering Calcium Homeostasis. PLoS Biol 2008, 6 (2), e26. https://doi.org/10.1371/journal.pbio.0060026.

74. Ong, D. S. T.; Mu, T.-W.; Palmer, A. E.; Kelly, J. W. Endoplasmic Reticulum Ca2+ Increases Enhance Mutant Glucocerebrosidase Proteostasis. Nat Chem Biol 2010, 6 (6), 424–432. https://doi.org/10.1038/nchembio.368.

75. Cox, J.; Hein, M. Y.; Luber, C. A.; Paron, I.; Nagaraj, N.; Mann, M. Accurate Proteome-Wide Label-Free Quantification by Delayed Normalization and Maximal Peptide Ratio Extraction, Termed MaxLFQ. Mol Cell Proteomics 2014, 13 (9), 2513–2526. https://doi.org/10.1074/mcp.M113.031591.

76. Lobingier, B. T.; Hüttenhain, R.; Eichel, K.; Miller, K. B.; Ting, A. Y.; Zastrow, M.; Krogan, N. J. An Approach to Spatiotemporally Resolve Protein Interaction Networks in Living Cells. Cell 2017, 169 (2), 350–360 312. https://doi.org/10.1016/j.cell.2017.03.022.

77. Berg Luecke, L.; Gundry, R. L. Assessment of Streptavidin Bead Binding Capacity to Improve Quality of Streptavidin-Based Enrichment Studies. J. Proteome Res. 2021, 20 (2), 1153– 1164. https://doi.org/10.1021/acs.jproteome.0c00772.

78. Förster, B.; Demangel, C.; Thye, T. Mycolactone Induces Cell Death by SETD1B-Dependent Degradation of Glutathione. PLoS Negl Trop Dis 2020, 14 (10), e0008709. https://doi.org/10.1371/journal.pntd.0008709.

79. Kang, S.-W.; Rane, N. S.; Kim, S. J.; Garrison, J. L.; Taunton, J.; Hegde, R. S. Substrate-Specific Translocational Attenuation during ER Stress Defines a Pre-Emptive Quality Control Pathway. Cell 2006, 127 (5), 999–1013. https://doi.org/10.1016/j.cell.2006.10.032.

