## Supplemental Figures for "Quantitative Measurement of Secretory Protein Mistargeting by Proximity Labeling and Parallel Reaction Monitoring"

| **Page** | **Contents** |
| --- | --- |
| **S-1** | **Table of Contents** |
| **S-2–3** | **Supplemental Figure S1** |
| **S-4** | **Supplemental Figure S2a** |
| **S-5** | **Supplemental Figure S2b** |
| **S-6–7** | **Supplemental Figure S3** |
| **S-8** | **Supplemental Figure S4** |
| **S-9–10** | **Supplemental Figure S5a** |
| **S-11–12** | **Supplemental Figure S5b** |
| **S-13–14** | **Supplemental Figure S5c** |
| **S-15** | **Supplemental Figure S5d** |
| **S-16** | **Supplemental Figure S5e** |
| **S-17** | **Supplemental Figure S5f** |
| **S-18** | **Supplemental Figure S5g** |
| **S-19** | **Supplemental Figure S5h** |
| **S-20–21** | **Supplemental Figure S5i** |
| **S-22–26** | **Supplemental Figure S6** |
| **S-27–29** | **Supplemental Figure S7** |
| **S-30** | **Supplemental Table S1** |

Supplemental **Table S2** is provided in a separate .xlsx spread sheet.

**Figure S1**

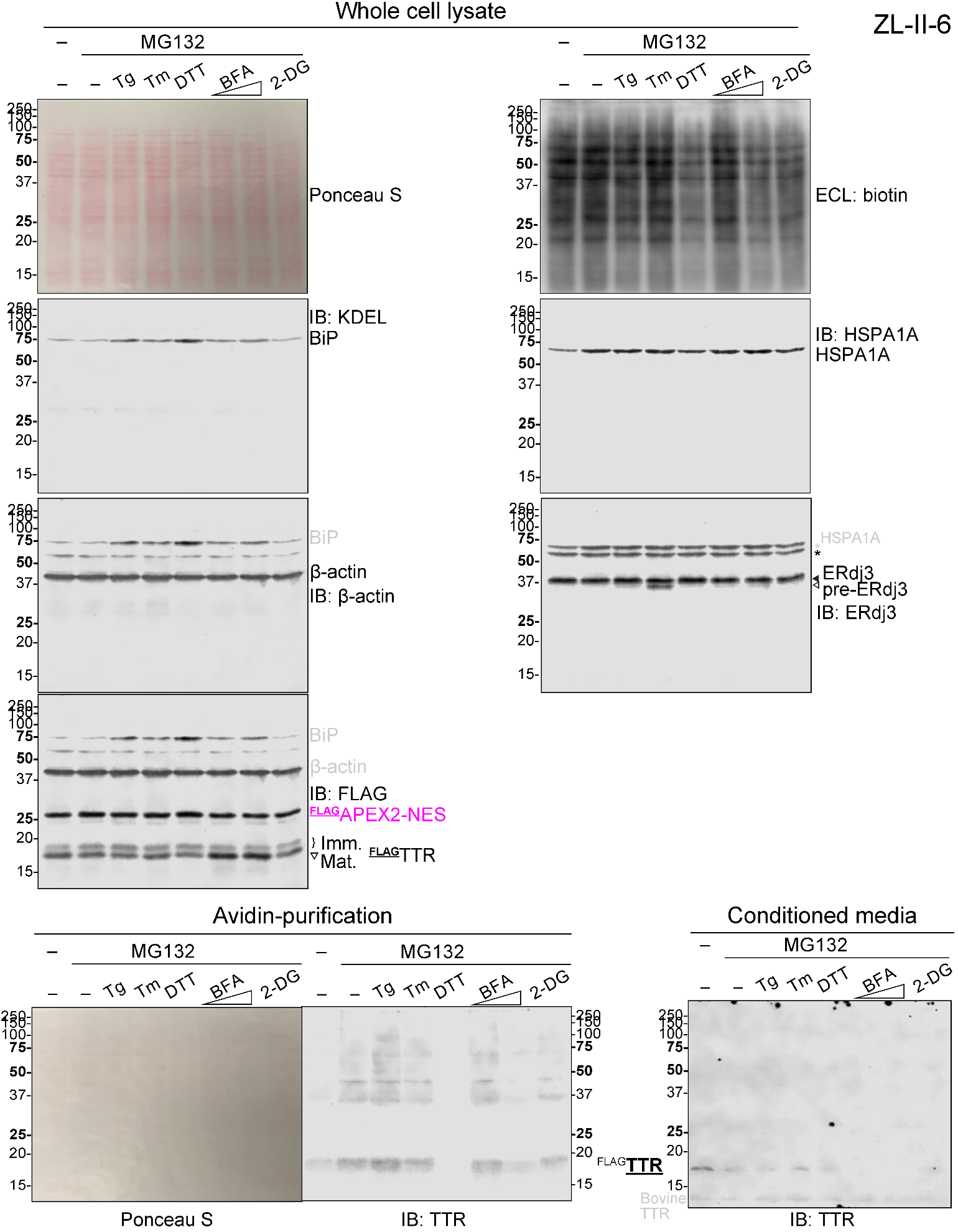

**Figure S1.** Measurement of ER stress-induced ^FLAG^TTR mistargeting in HEK293T cells by immunoblotting (IB), prior to parallel reaction monitoring (PRM) method development. Cells were treated with common ER stressors and proteasome inhibitor MG132 as indicated. Ponceau S stain, electrochemiluminescence (ECL) of biotin and IB full blots related are displayed. Dosages: Tg: 50 nM; Tm : 200 nM; DTT: 3 mM; BFA: 250 and 750 ng mL^–1^; 2-DG: 2 mM. 2 equiv. ^FLAG^TTR plasmid was used in this experiment. ER stress by Tg, Tm, DTT or BFA was confirmed by the upregulation of BiP (IB: KDEL). 2 mM 2-DG does not induce ER stress in HEK293T cells. The inhibition of *N*-glycan synthesis during Tm treatment was confirmed by the appearance of the pre-ERdj3 band (*N*-glycan naïve form) from IB (IB: ERdj3, lane Tm). The inhibition of secretion by BFA was indicated by increased steady state ^FLAG^TTR level in lysate or by decreased secreted ^FLAG^TTR in conditioned media (IB: TTR, lane BFA). 3 mM DTT, 750 ng mL^–1^ BFA and 2 mM 2-DG decrease proximity labeling efficiency on the basis of ECL.

**Figure S2a**

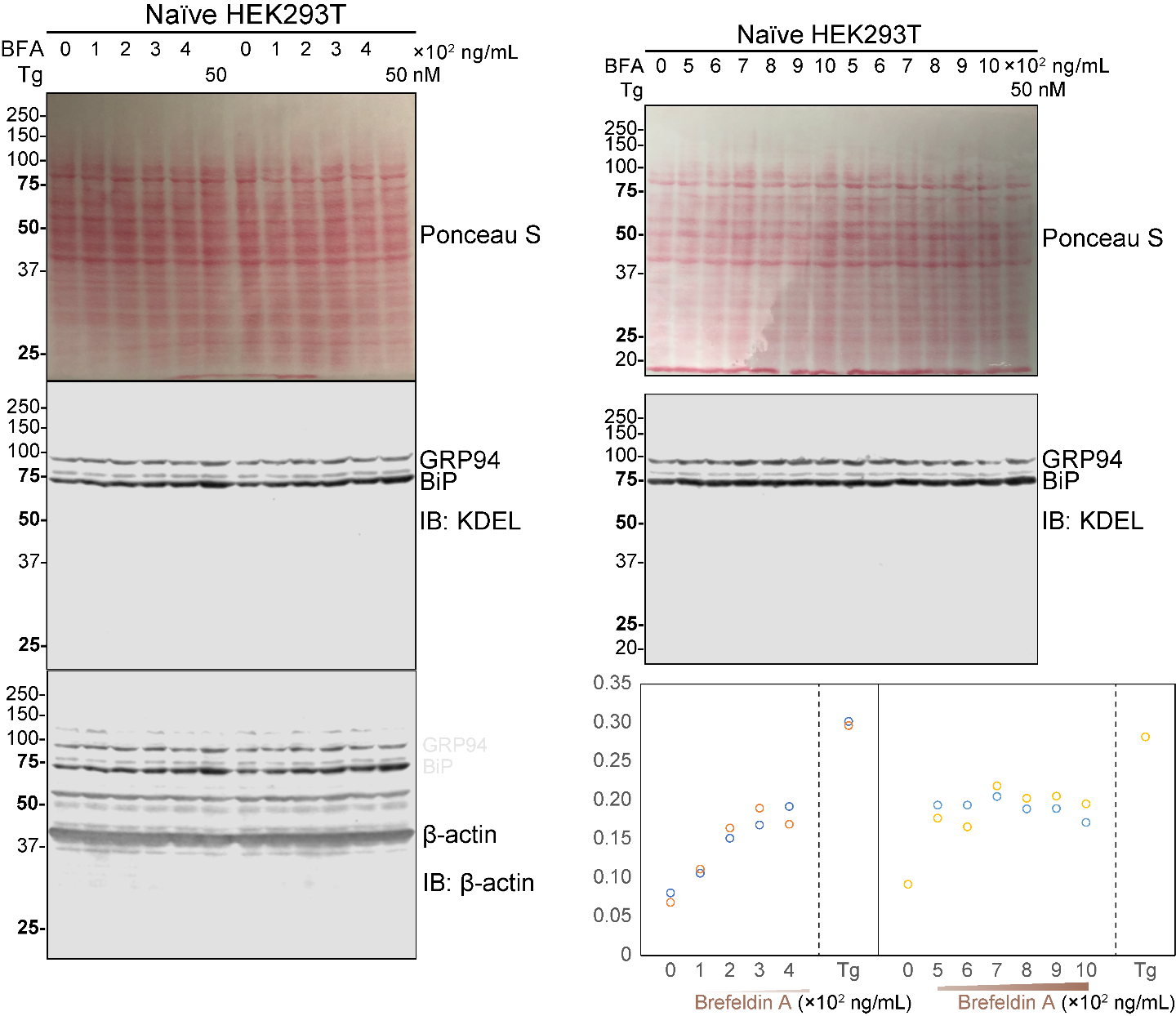

**Figure S2b**

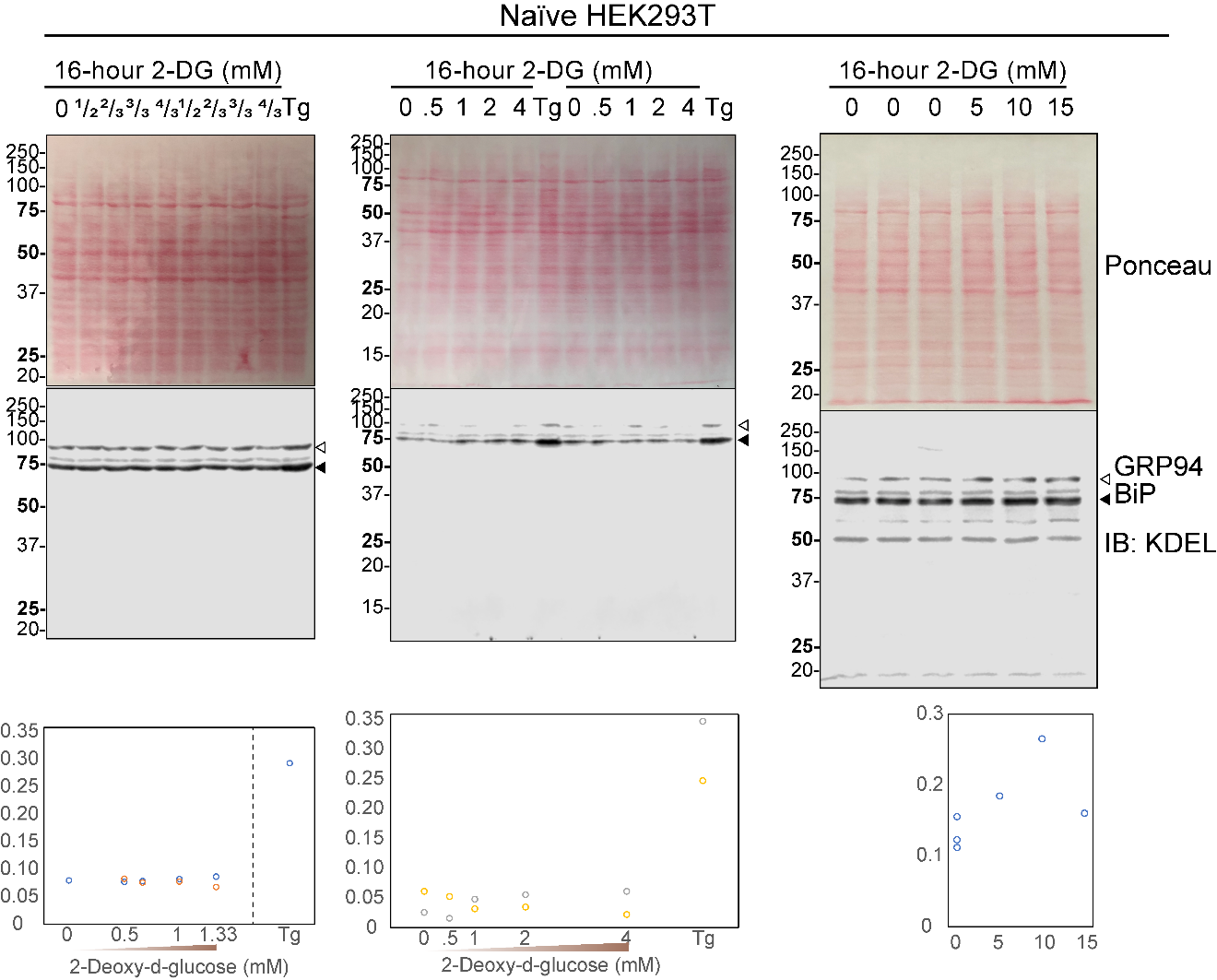

**Figure S2.** Full blots related to dosability experiments of selected ER stressors, aiming to induce maximum BiP upregulation. **a**) Dosability of BFA on BiP upregulation. Ponceau S stain and IB: β-actin serve as loading control. Normalization of BiP band densitometry was performed by dividing each condition by the sum of BiP densitometries across all conditions within each replicate. **b**) Dosability of 2-DG on BiP upregulation. Ponceau S stain serves as loading control. in the middle blot, naïve HEK293T lysate samples were analyzed in a 12% w/v resolving gel, instead of 10% w/v. Normalization of BiP band densitometry was performed by dividing each condition by the sum of BiP densitometries across all conditions within each replicate.

**Figure S3**

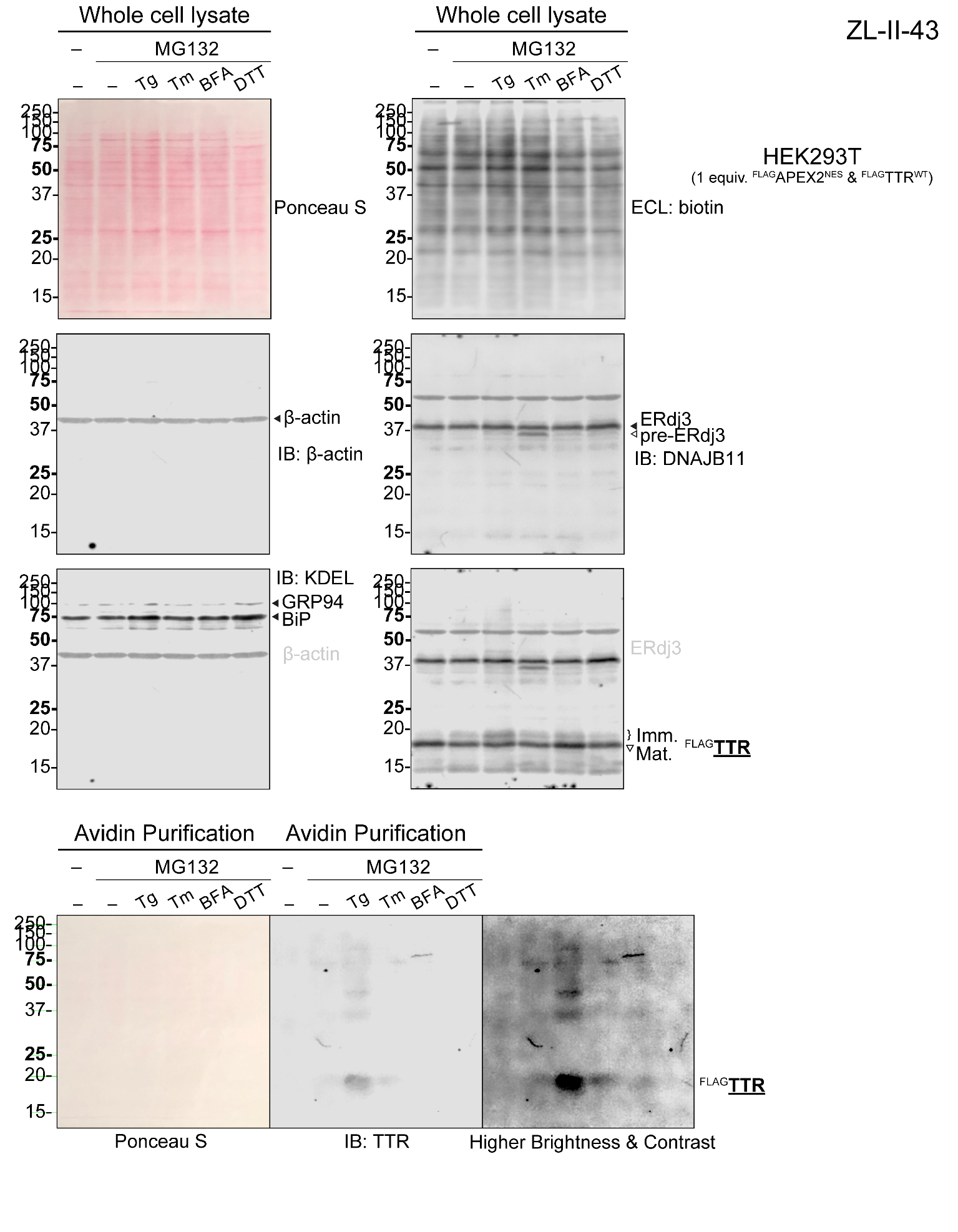

**Figure S3.** Measurement of ER stress-induced ^FLAG^TTR mistargeting in HEK293T cells by IB, after dosability titration and prior to PRM method development. Cells were treated with common ER stressors and proteasome inhibitor MG132 as indicated. Ponceau S stain, ECL of biotin and IB full blots related are displayed. Dosages: MG132: 1 μM; Tg: 50 nM; Tm : 200 nM; DTT: 3 mM; BFA: 300 ng mL^–1^. Normal equivalence ^FLAG^TTR plasmid was used in this experiment. ER stress was confirmed by the upregulation of BiP (IB: KDEL). The inhibition of *N*-glycan synthesis during Tm treatment was confirmed by the appearance of the pre-ERdj3 band (*N*-glycan naïve form) from IB (IB: ERdj3, lane Tm). The inhibition of secretion by BFA was indicated by increased steady state ^FLAG^TTR level in lysate (IB: TTR, lane BFA). By aspirating 3 mM DTT-containing media and replacement of fresh DMEM with 1 mM H_2_O_2_, proximity labeling efficiency is rescued.

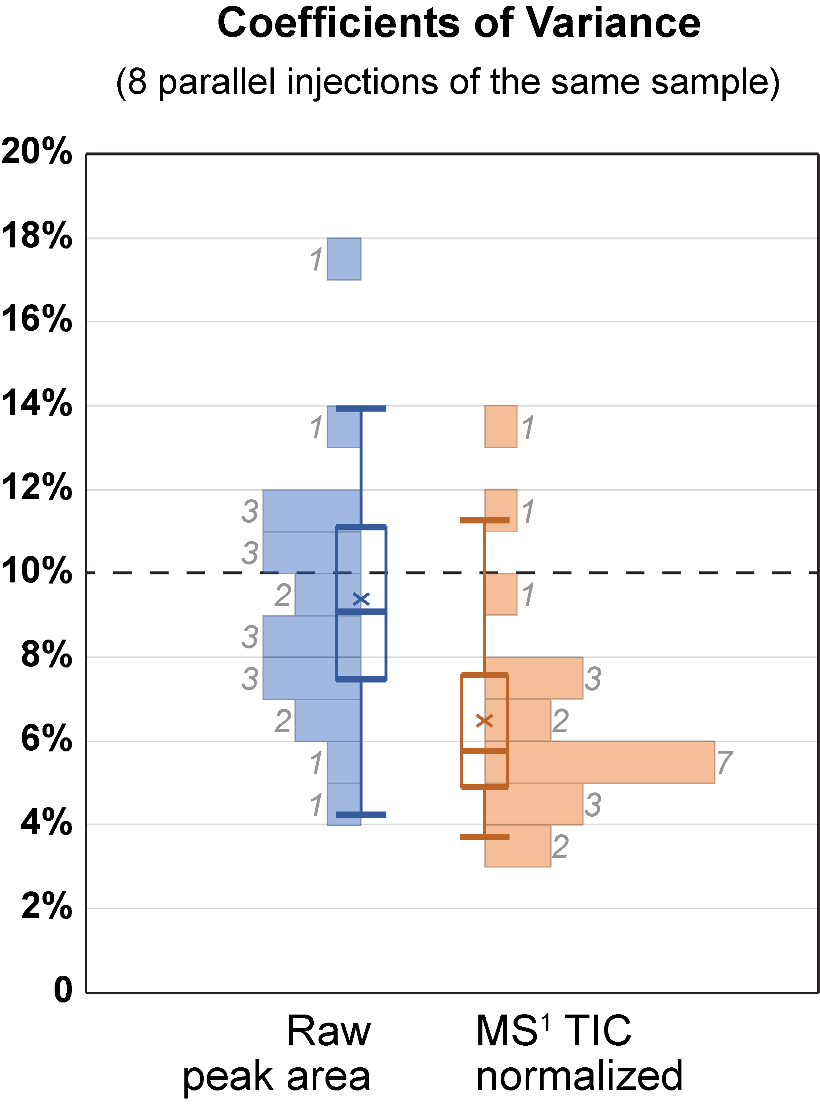

**Figure S4.** Coefficients of variance distribution of assayed peptides. Coefficients of variance (CVs) distribution of assayed peptides (raw peak areas or normalized by MS^1^ total ion current chromatograms) from 8 parallel injections. Box plots show whisker, quartiles, median and mean of CVs. Histograms have bin width of 1% CV, with frequencies falling in each bin labeled.

**Figure S5a** Targeted ^FLAG^TTR peptide

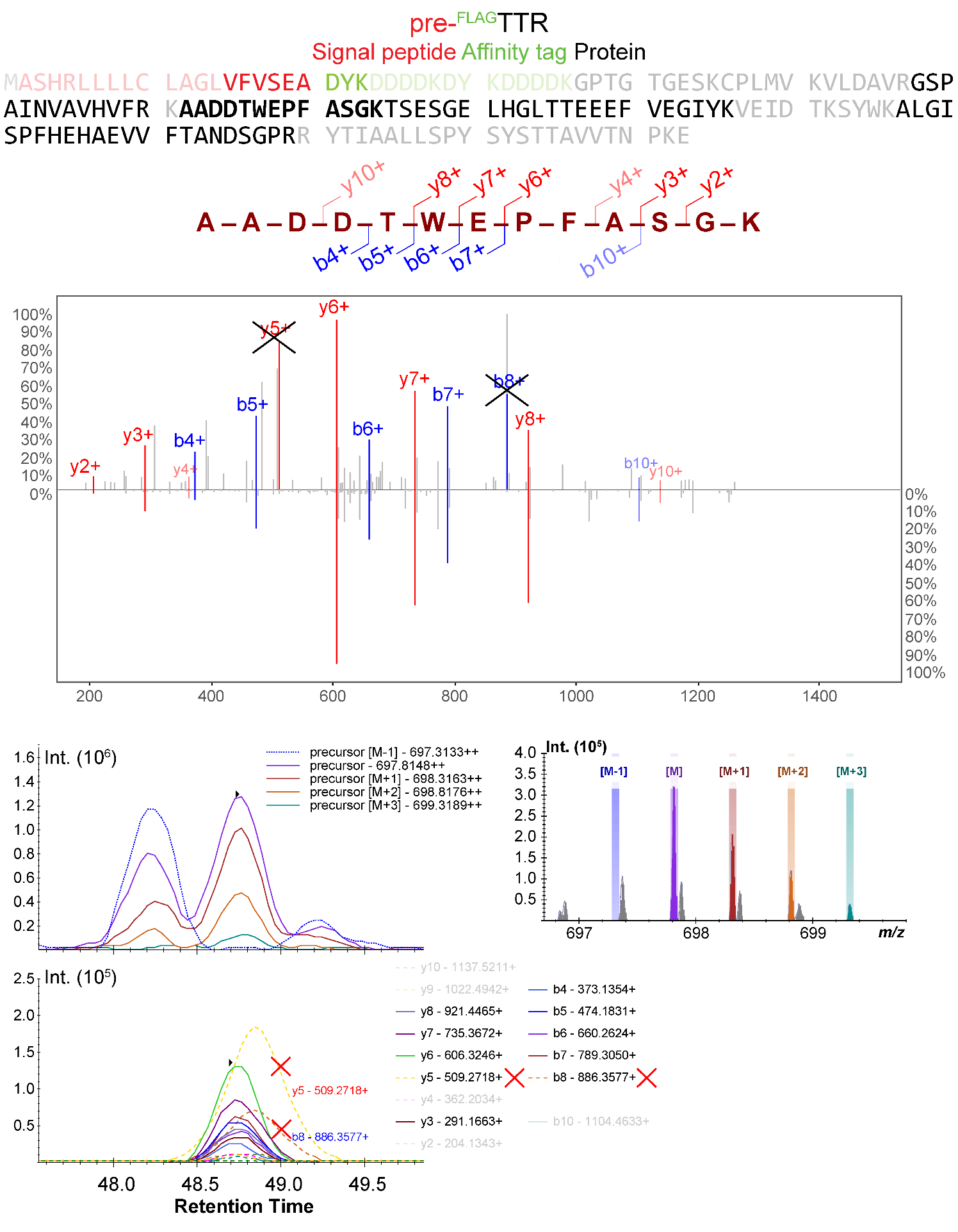

**Supplemental comment: Figure S5a** displays how the specificity of a peptide was evaluated. MS^2^ spectrum acquired in a PRM experiment was displayed in Lorikeet Spectrum viewer, with a default mass tolerance of 0.5 Th, and a minimum signal-to-noise ratio (S/N) of 0.8. Product ions in the representative MS^2^ with lower than 0.8 S/N are labeled in pale color and small font. CID spectra from NIST peptide library was mirrored at the bottom in the “butterfly mode” of Lorikeet spectrum viewer.

Other evidence includes: the chromatogram of precursor [M-1] must not align well with precursor series [M], [M+1], [M+2], *etc.* And the proportion among product ions should remain constant across different samples and injections. Product ions that do not align well with the precursor ions and the other product ions should be removed. For instance, the “y_5_^+^” and “b_8_^+^” in AADDTWEPFAGSK are excluded; they are also not shown in the reference spectrum. Production ions that interfere with background noise should also be removed.

**Figure S5b** Targeted ^cyt^APEX (^FLAG^APEX2^NES^) peptides, MS^2^ spectra (top) and Prosit predicted CID MS^2^ (bottom, NCE 35%)

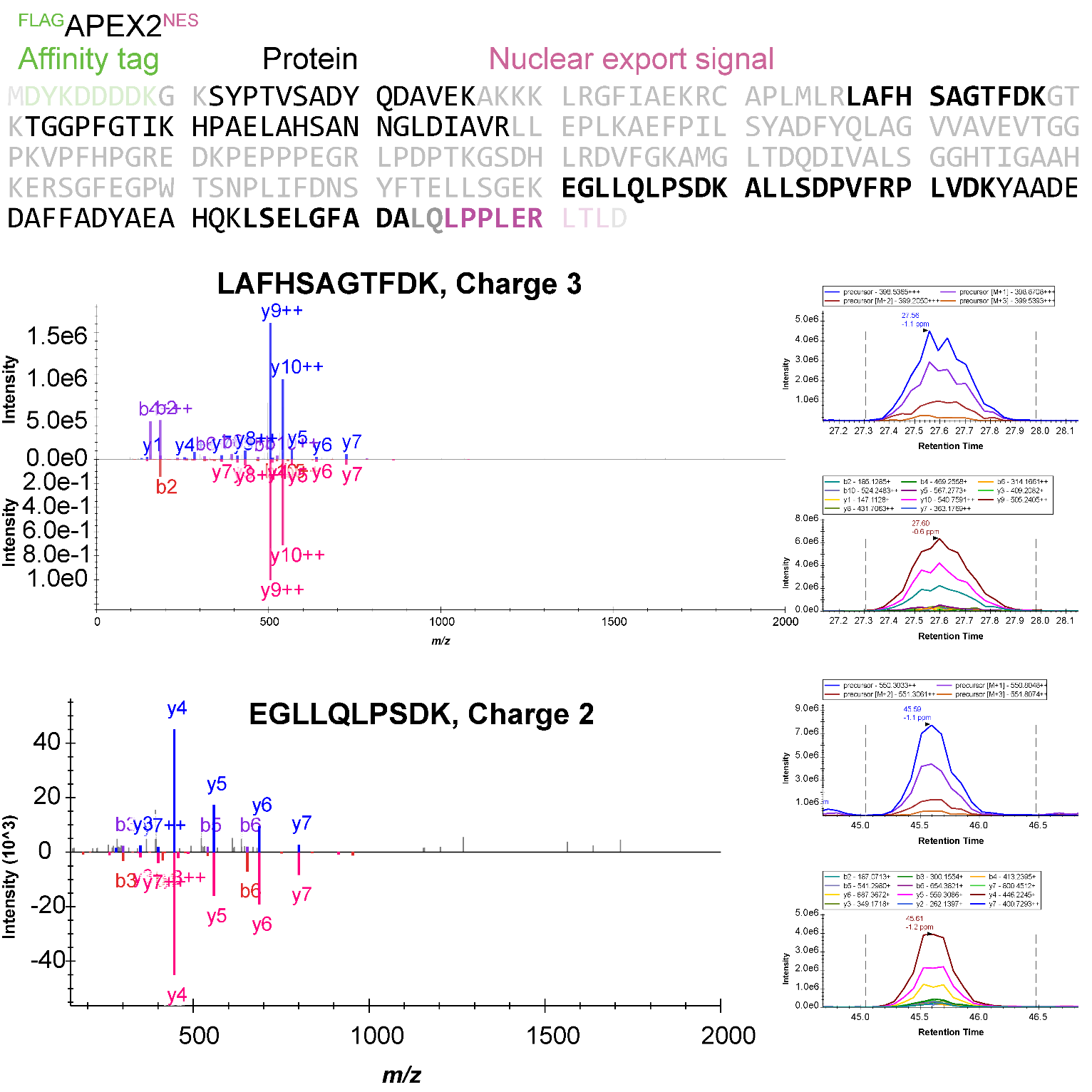

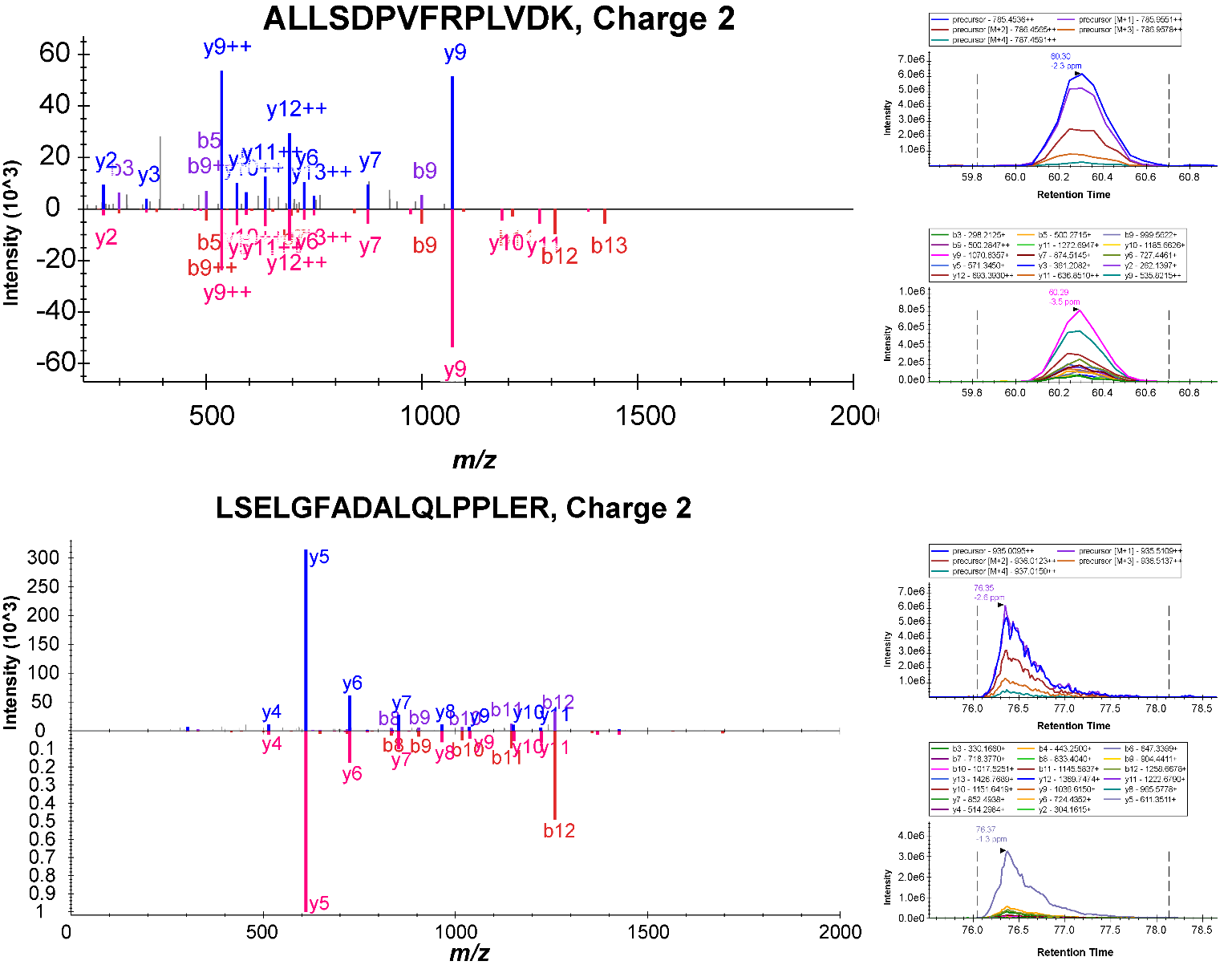

**Figure S5b** (continued, assayed peptide)

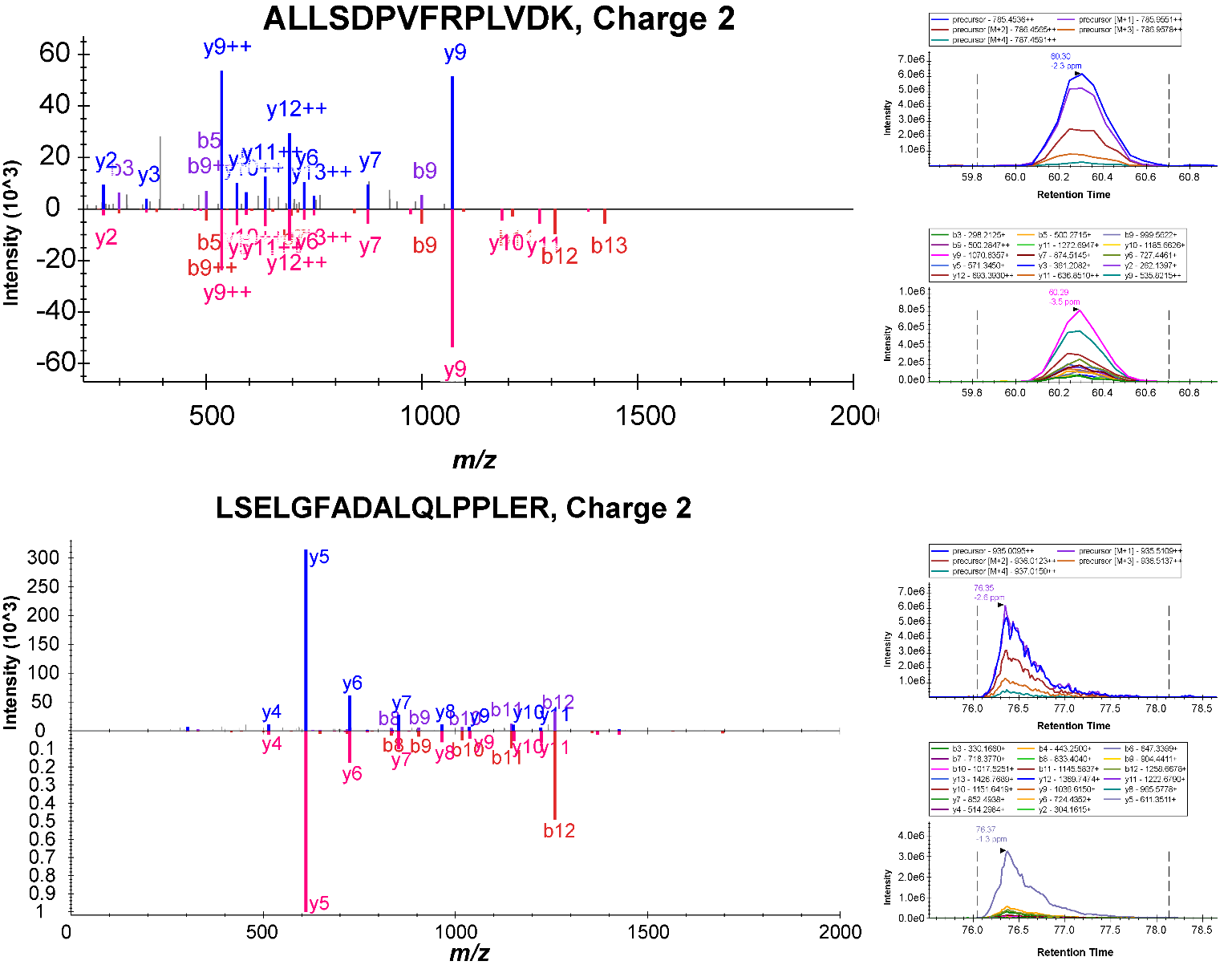

**Figure S5c** Targeted α-Tubulin (1B) peptides, MS^2^ spectra (top) and NIST CID MS^2^ (bottom)

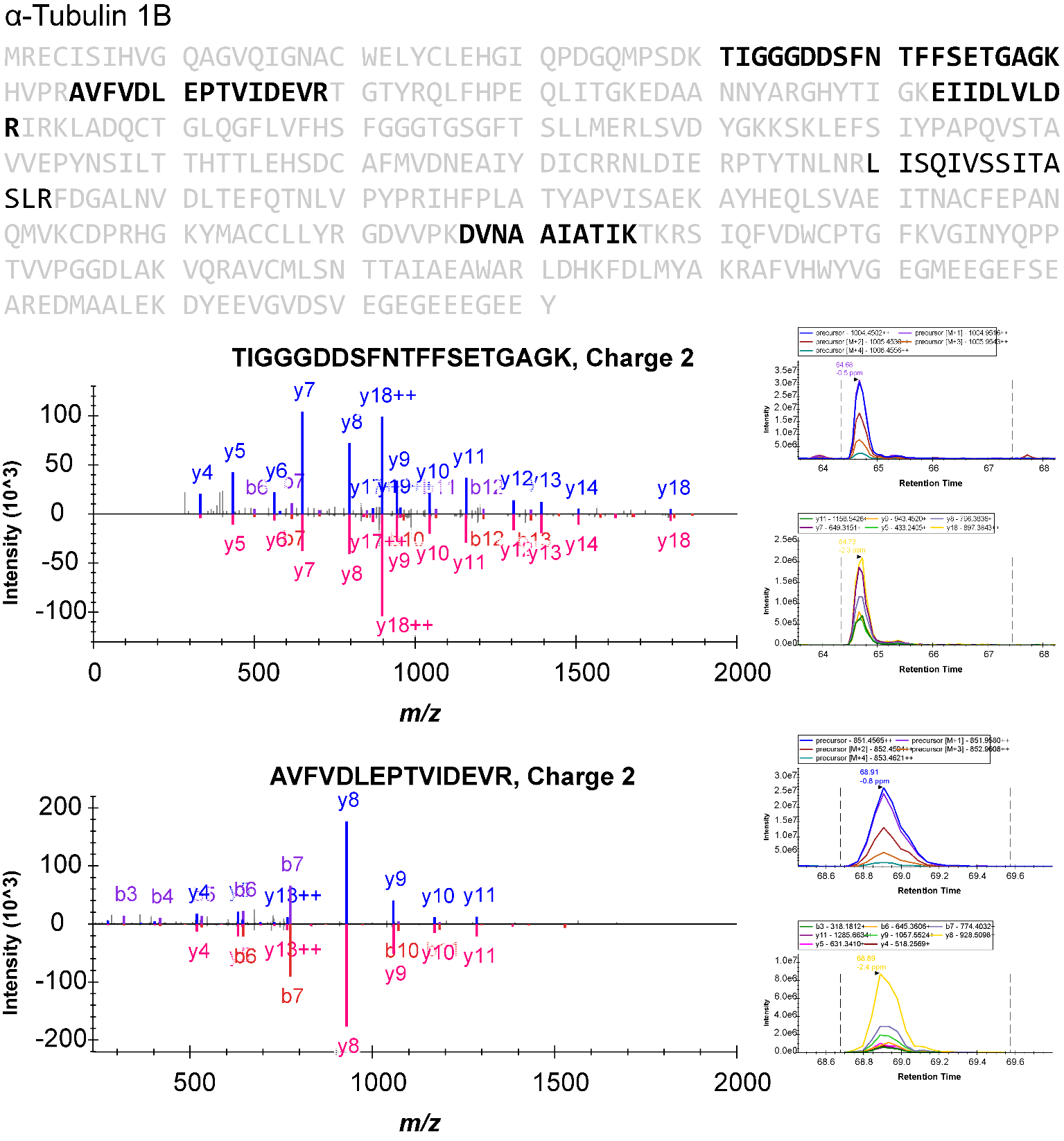

**Supplemental Comments:** This peptide is shared across α-Tubulin 1A/B/C and 3C/D/E.
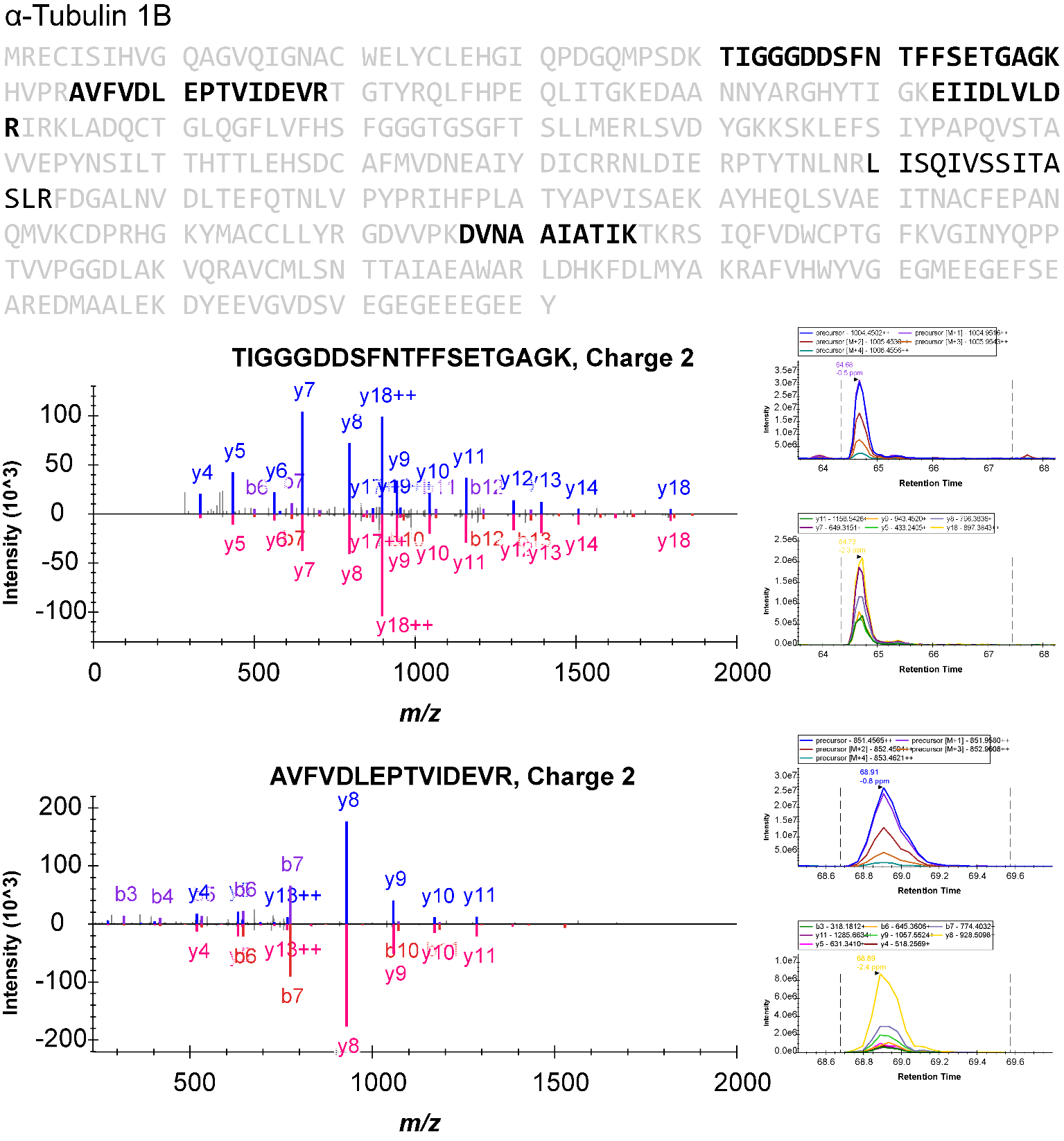

**Supplemental Comments:** This peptide is shared across α-Tubulin 1A/B/C.

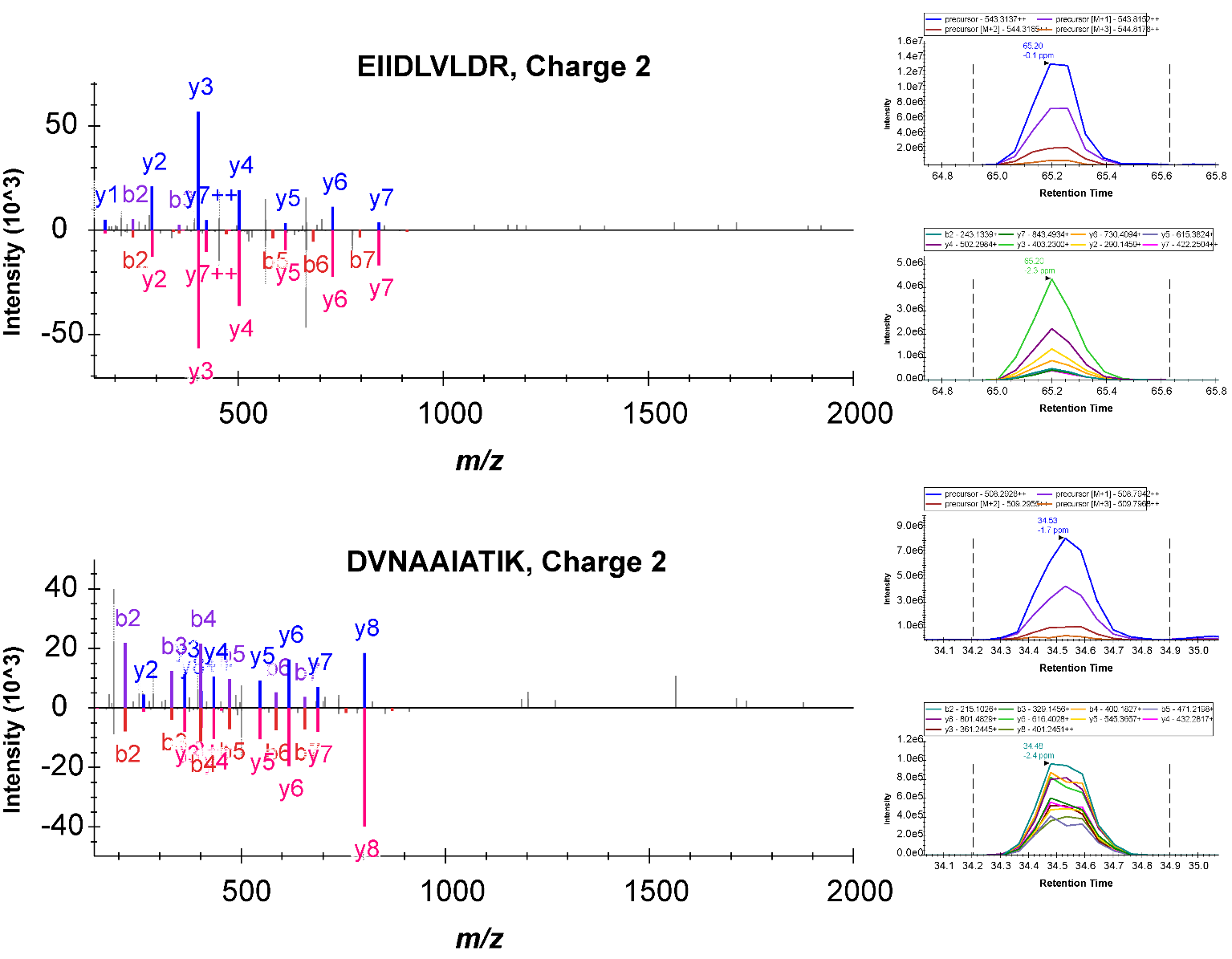

**Supplemental Comments:** This peptide is shared across α-Tubulin 1A/B/C.

**Figure S5c** (continued, assayed peptide)

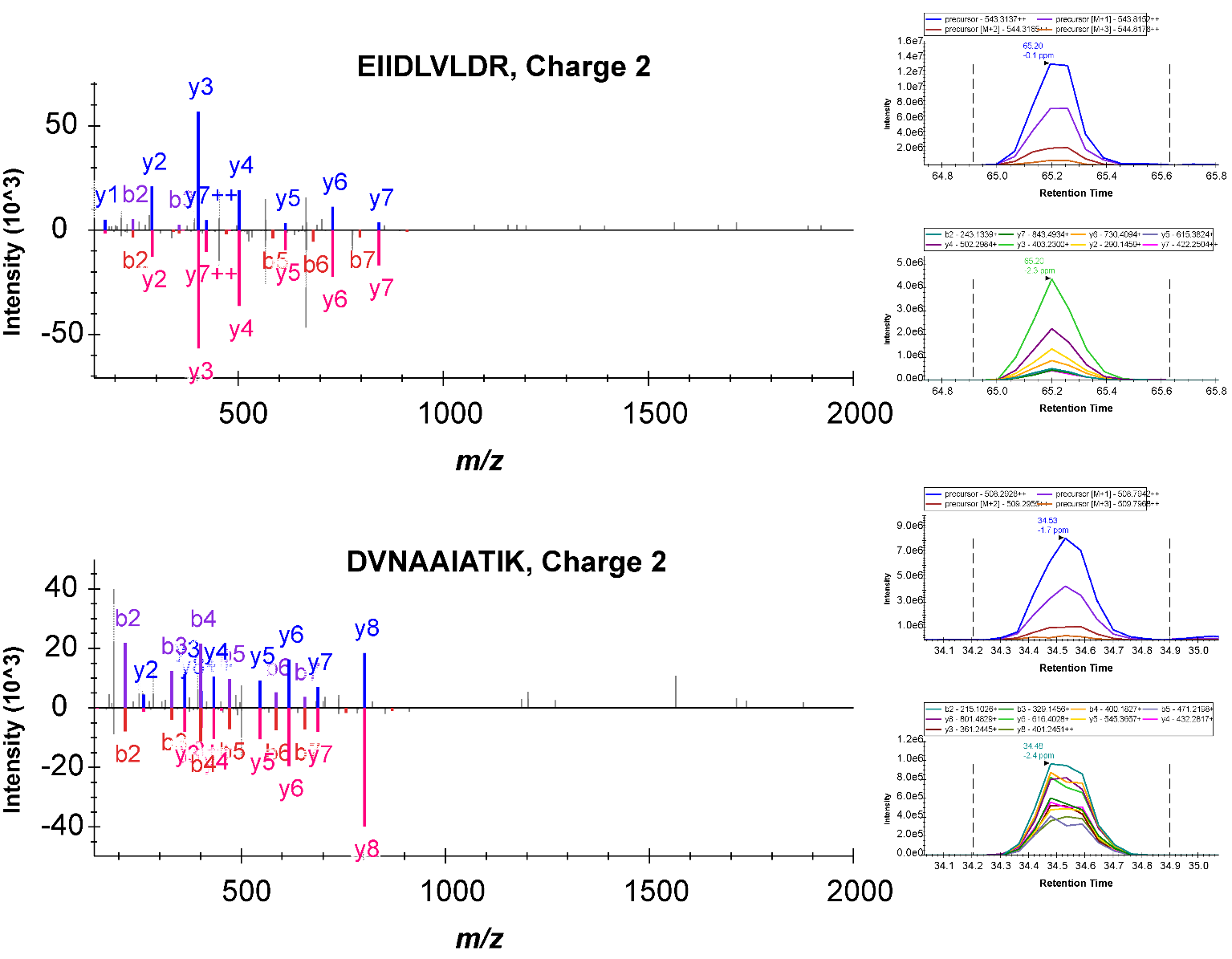

**Supplemental Comments:** This peptide is shared across α-Tubulin 1A/B/C and 3C/D/E.

**Figure S5d** Targeted β/γ-actin peptide, MS^2^ spectrum (top) and NIST CID MS^2^ (bottom)

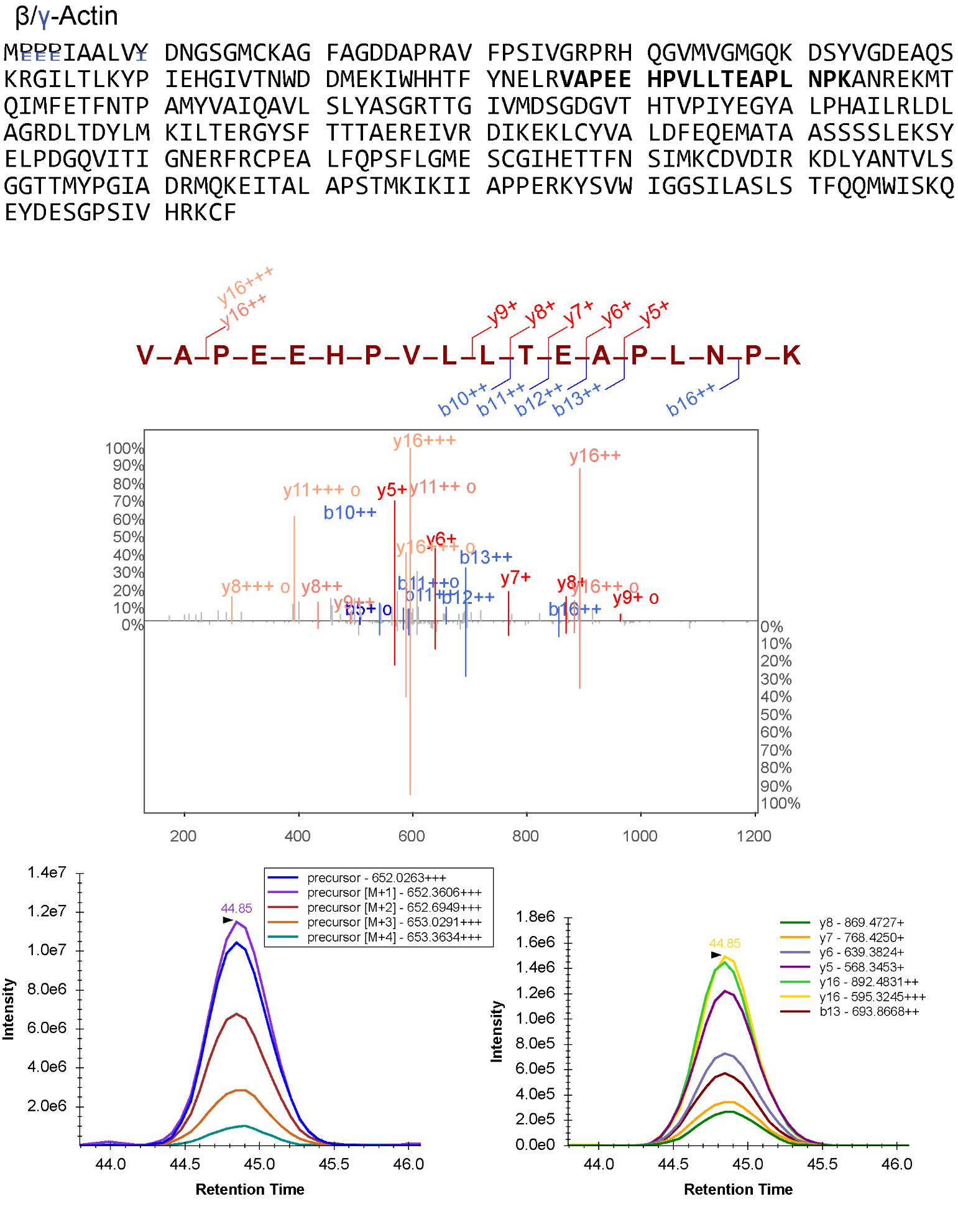

**Figure S5e** Targeted GAPDH peptide, MS^2^ spectra (top) and NIST CID MS^2^ (bottom)

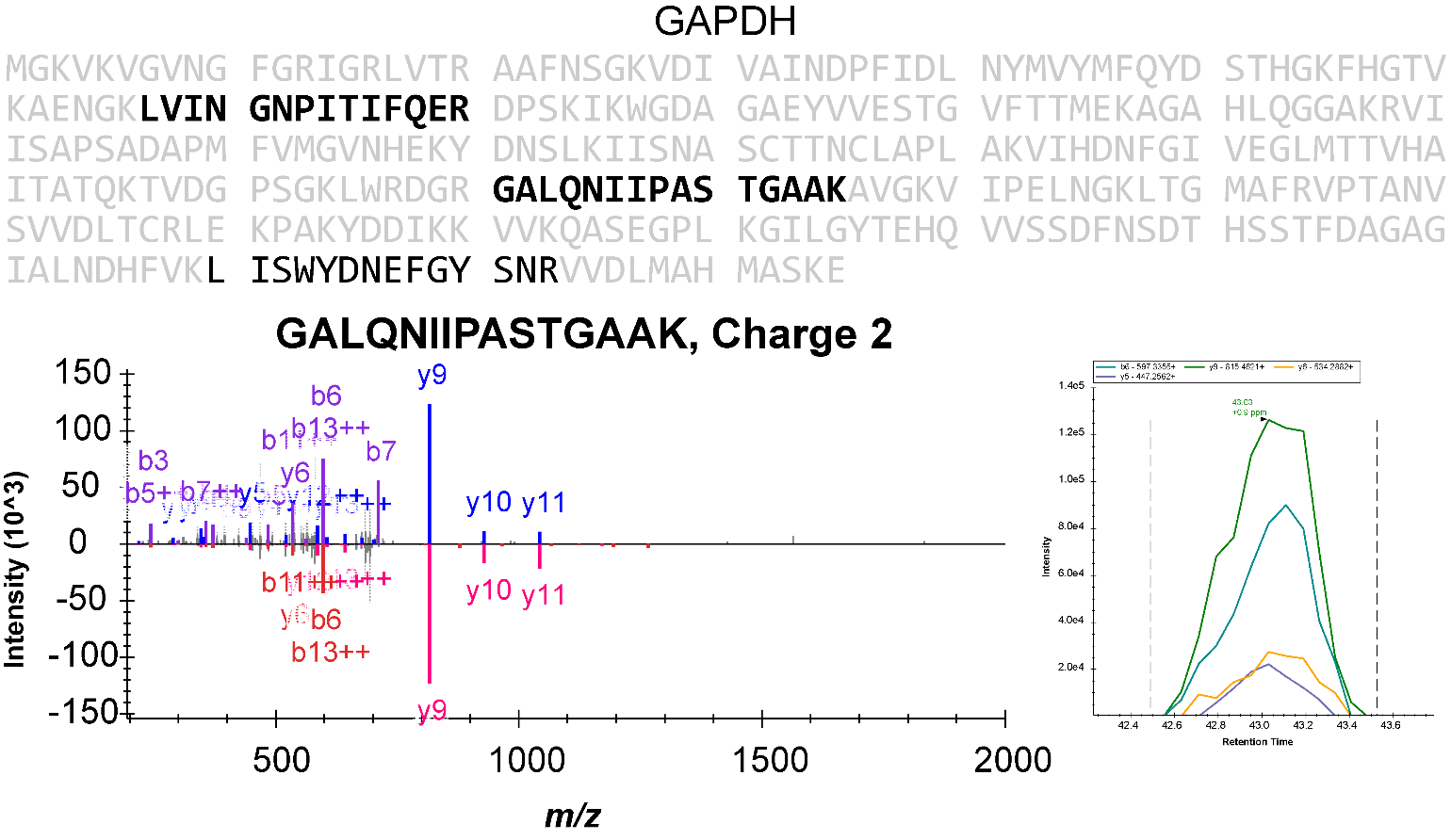

**Figure S5f** Targeted HSPA1A/B peptide, MS^2^ spectra (top) and NIST CID MS^2^ (bottom)

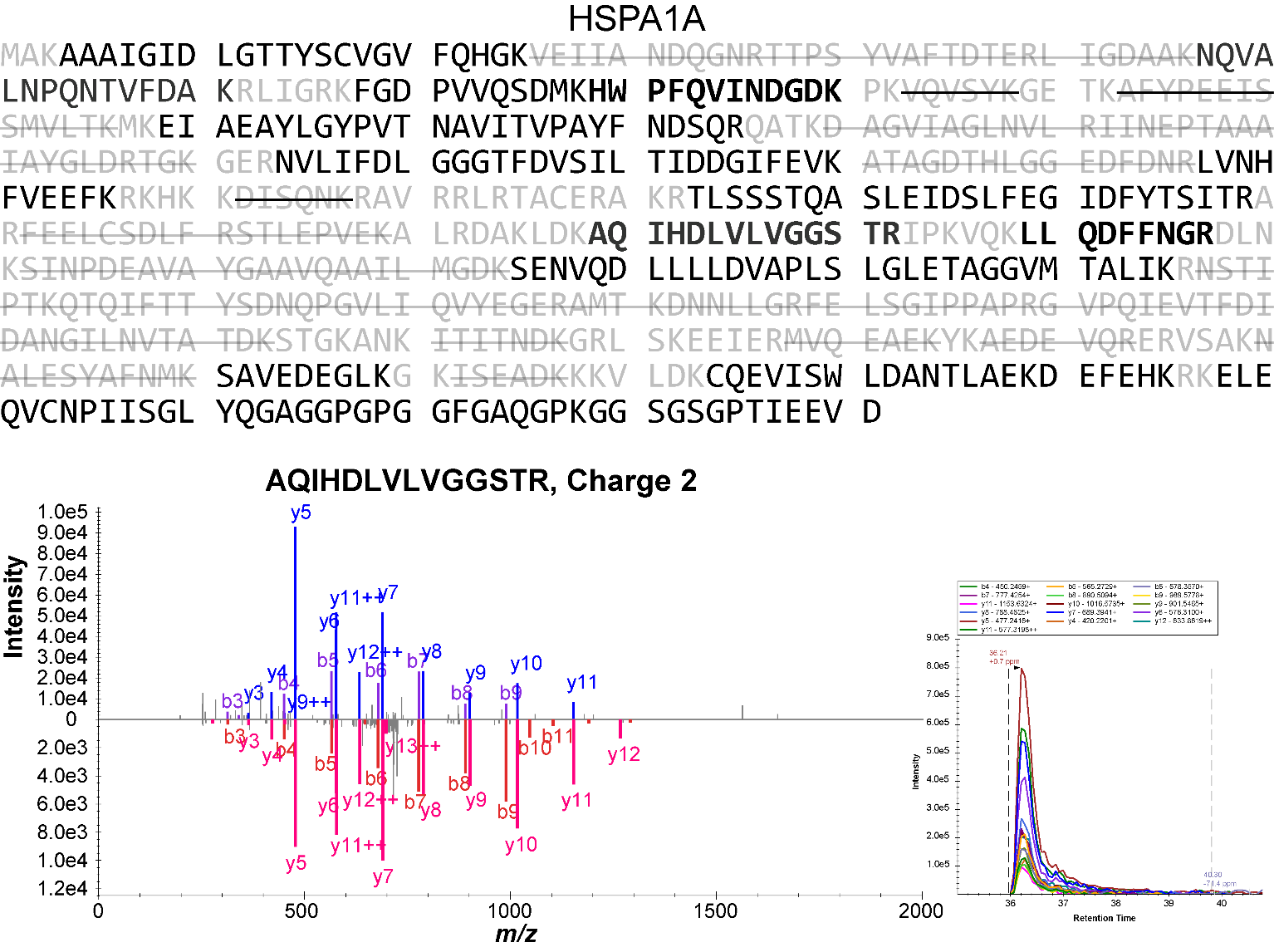

**Figure S5g** Targeted BiP peptides, MS^2^ spectra (top) and NIST CID MS^2^ (bottom)

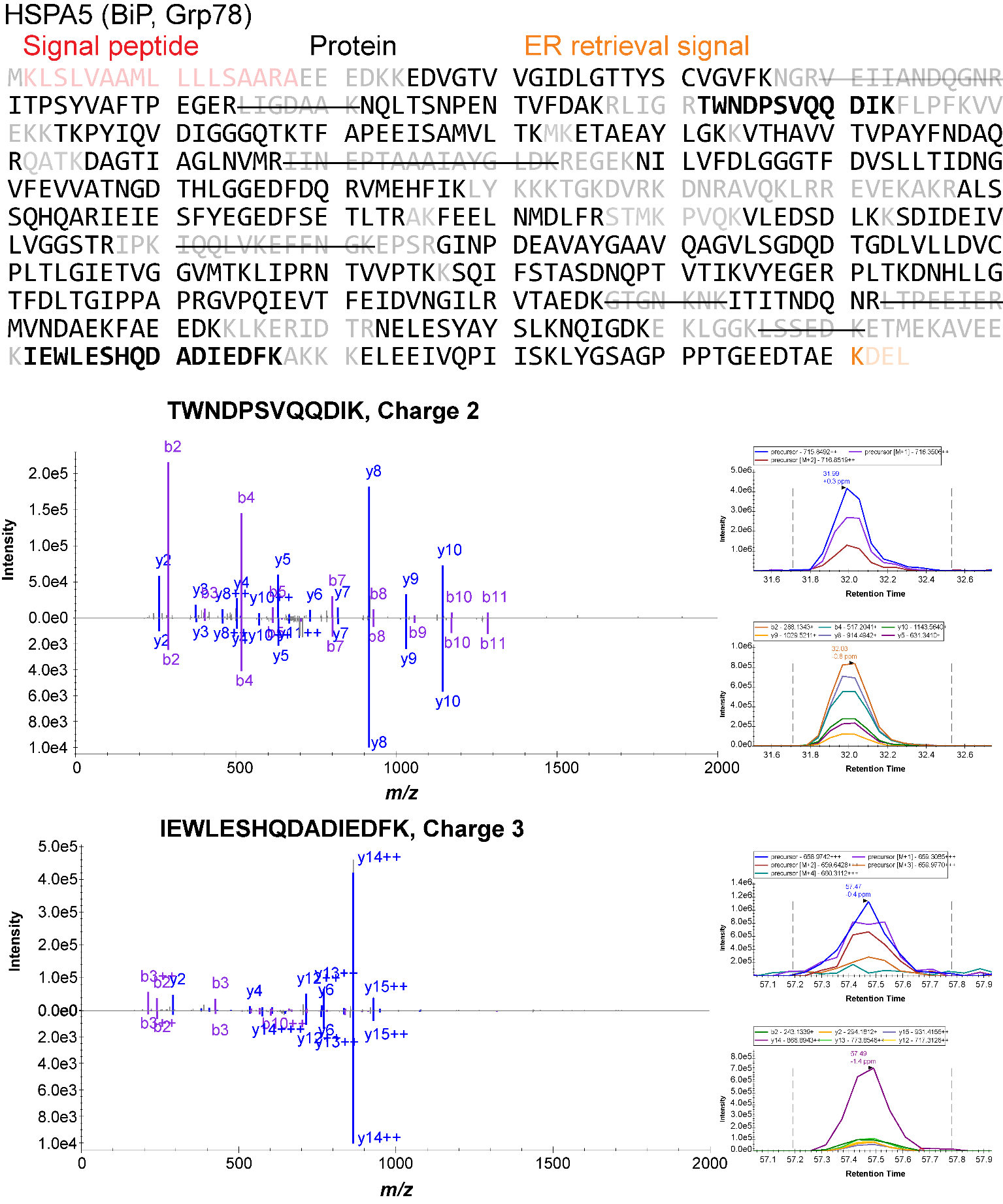

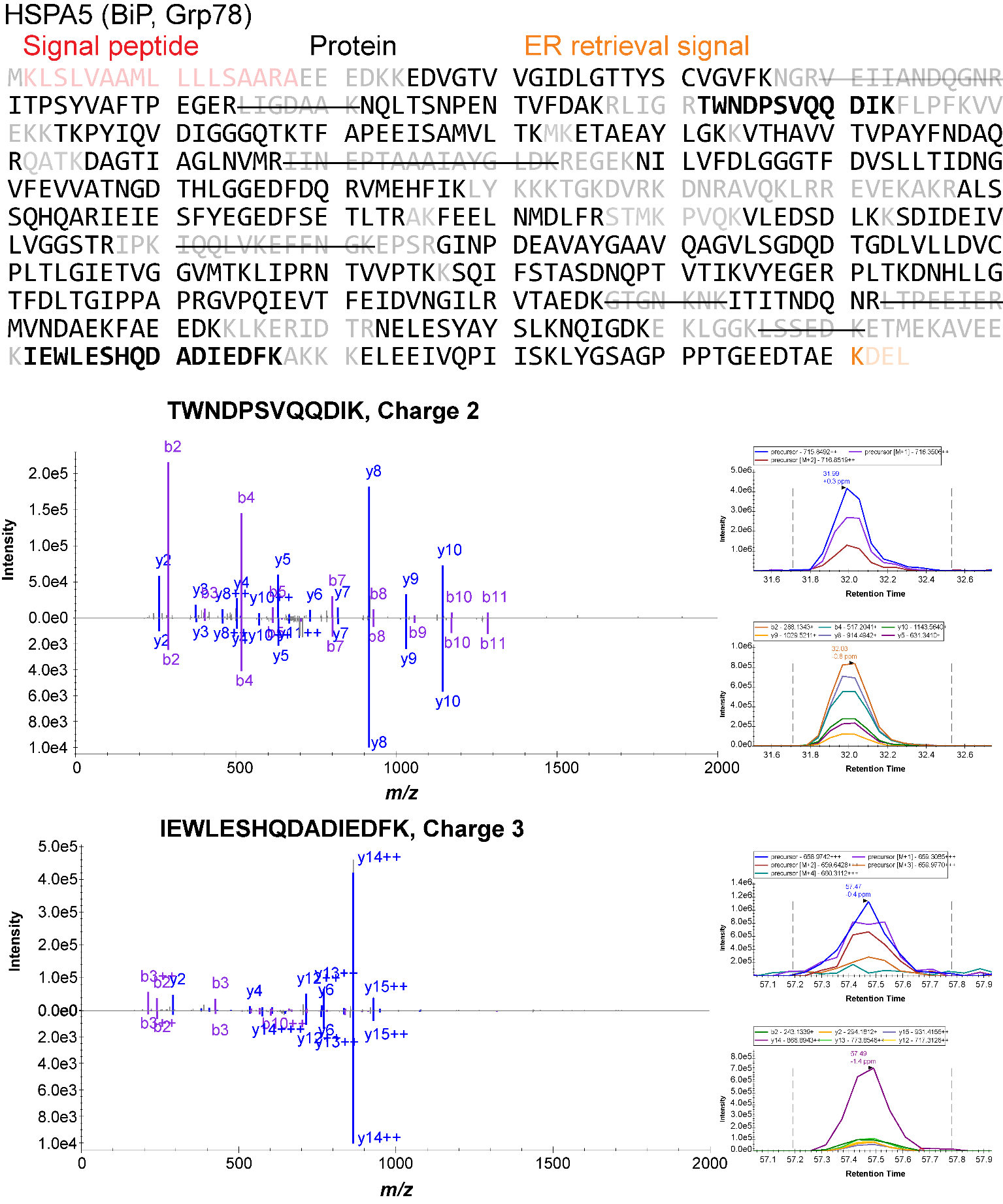

**Figure S5h** Targeted avidin peptide, MS^2^ spectra (top) and NIST CID MS^2^ (bottom)

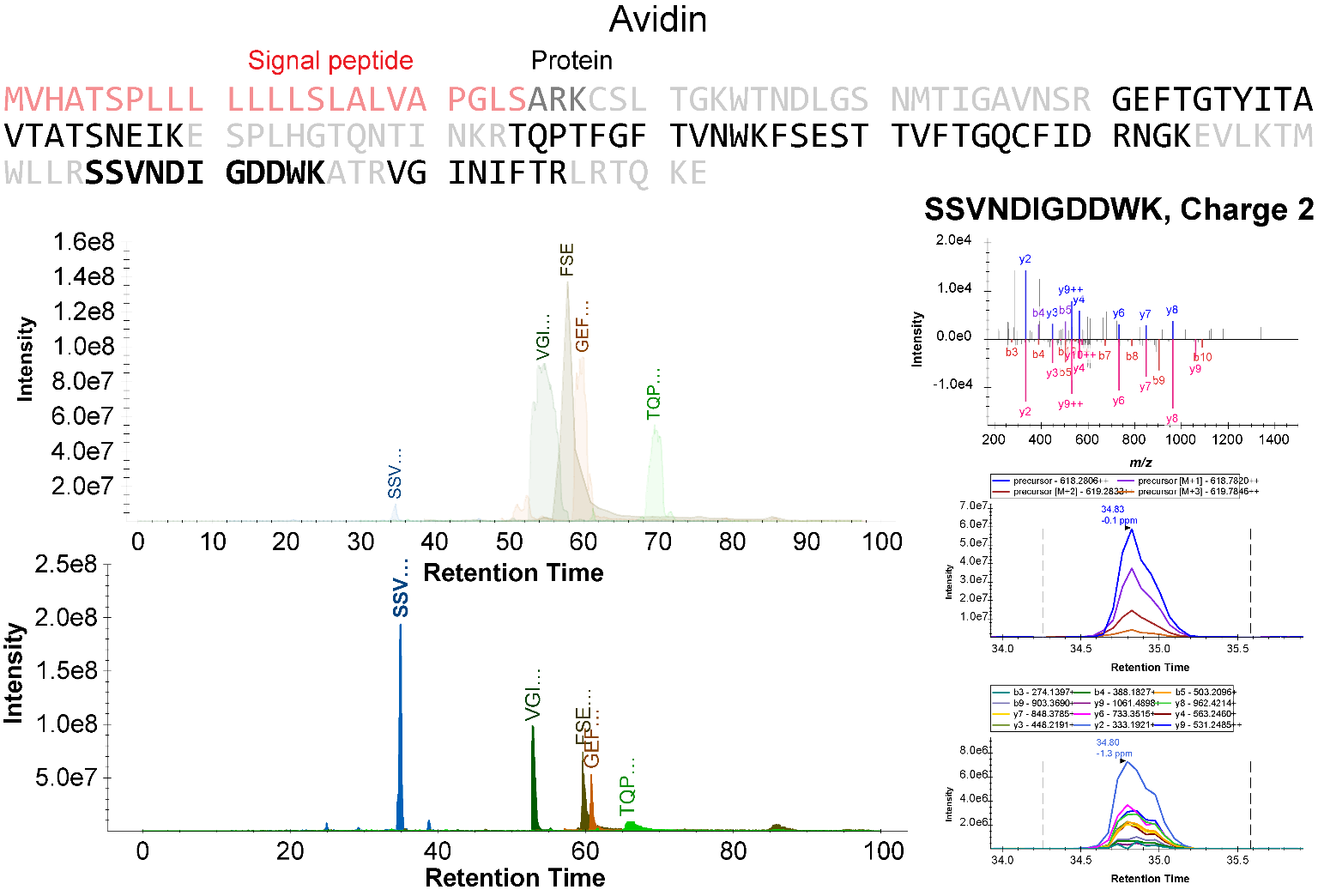

**Supplemental Comments:** The MS^1^ chromatograms of found peptides on the top were from data dependent analysis of avidin purified cell lysate. The bottom was from scheduled PRM of SSVNDIGDDWK^2+^ and MS^1^ filtering of the other peptides. Only SSVNDIGDDWK^2+^ was included in the PRM methods because it is less overlapped with other scheduled targets.

**Figure S5i** Targeted pyruvate carboxylase peptides, MS^2^ spectra (top) and NIST CID MS^2^ (bottom)

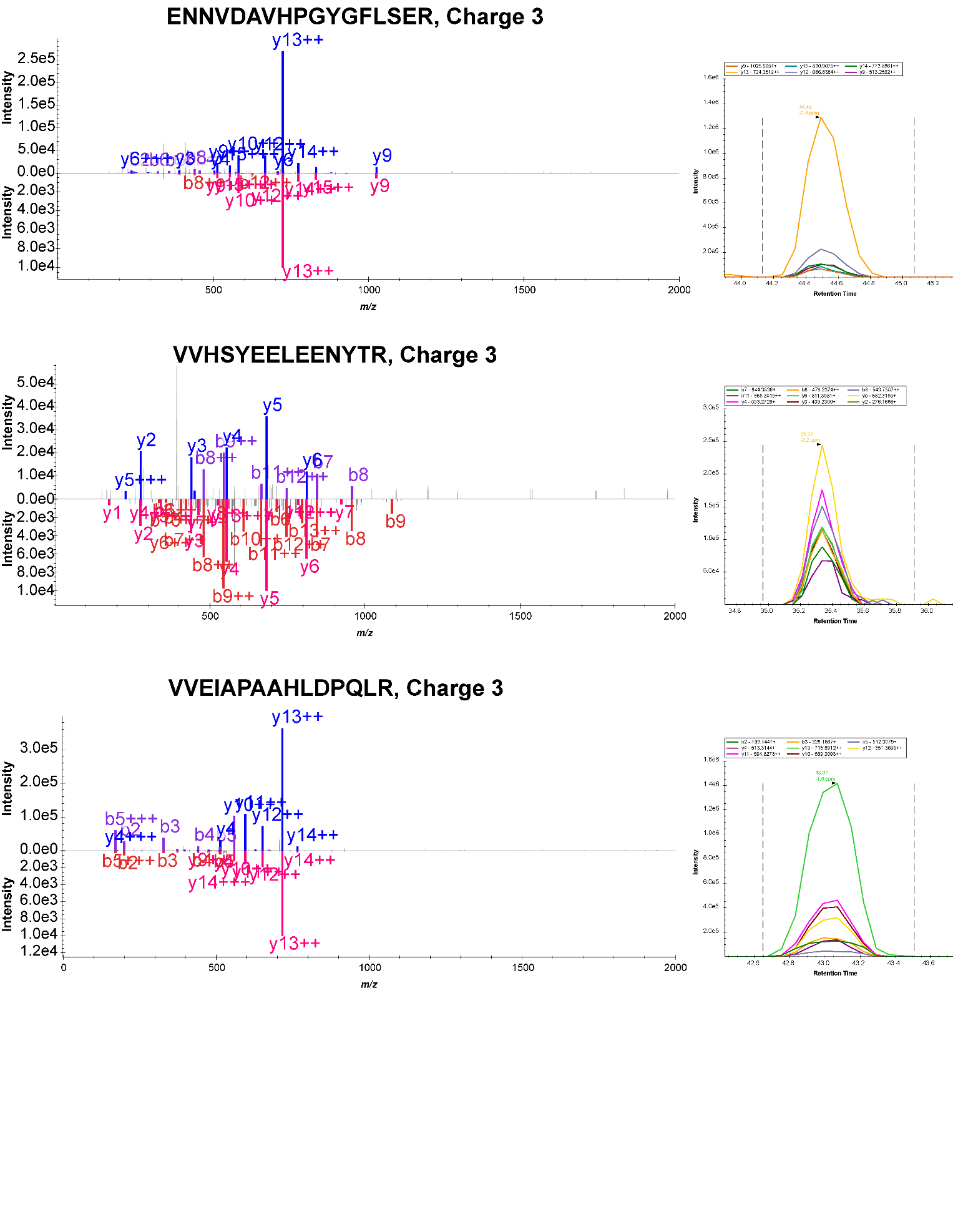

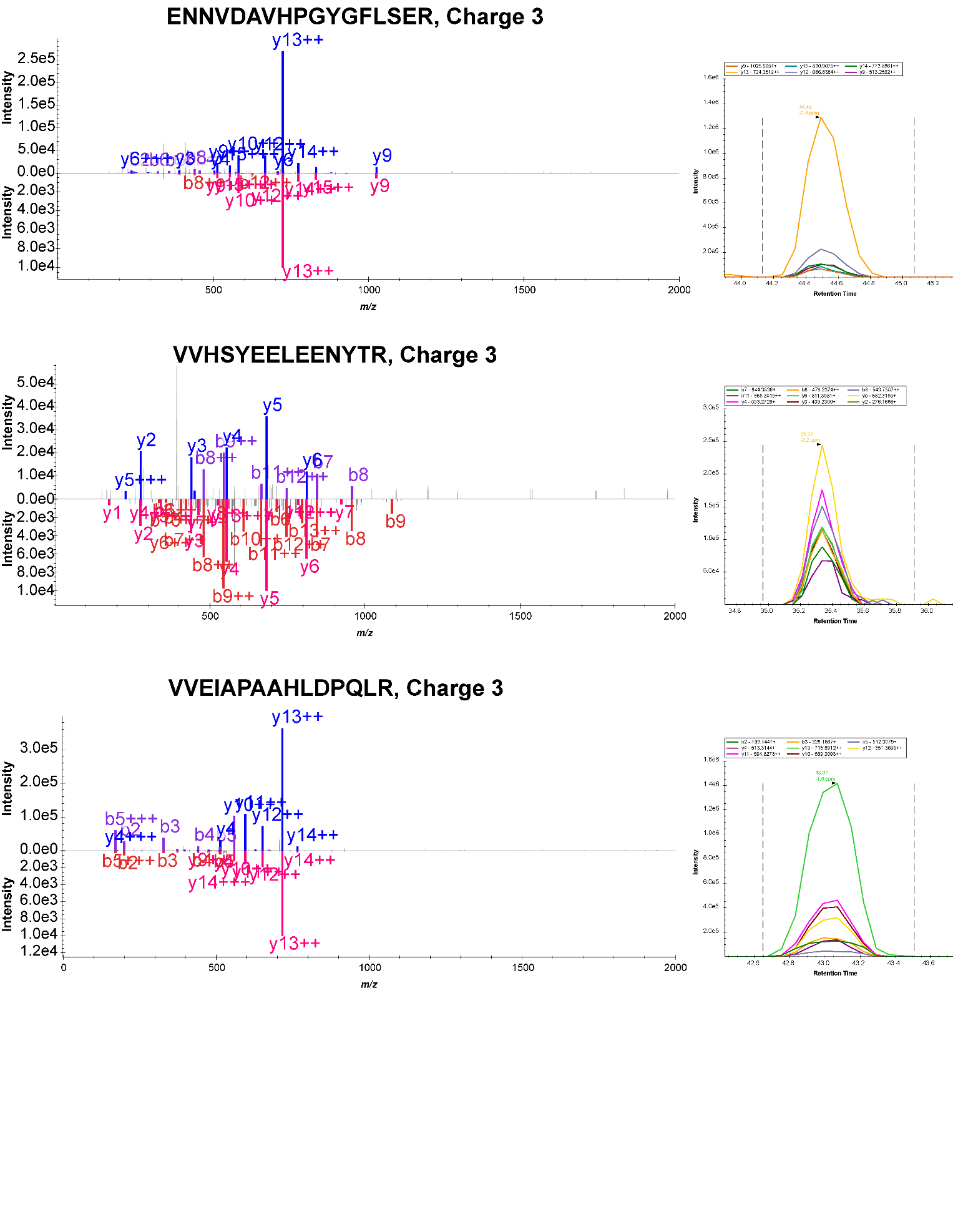

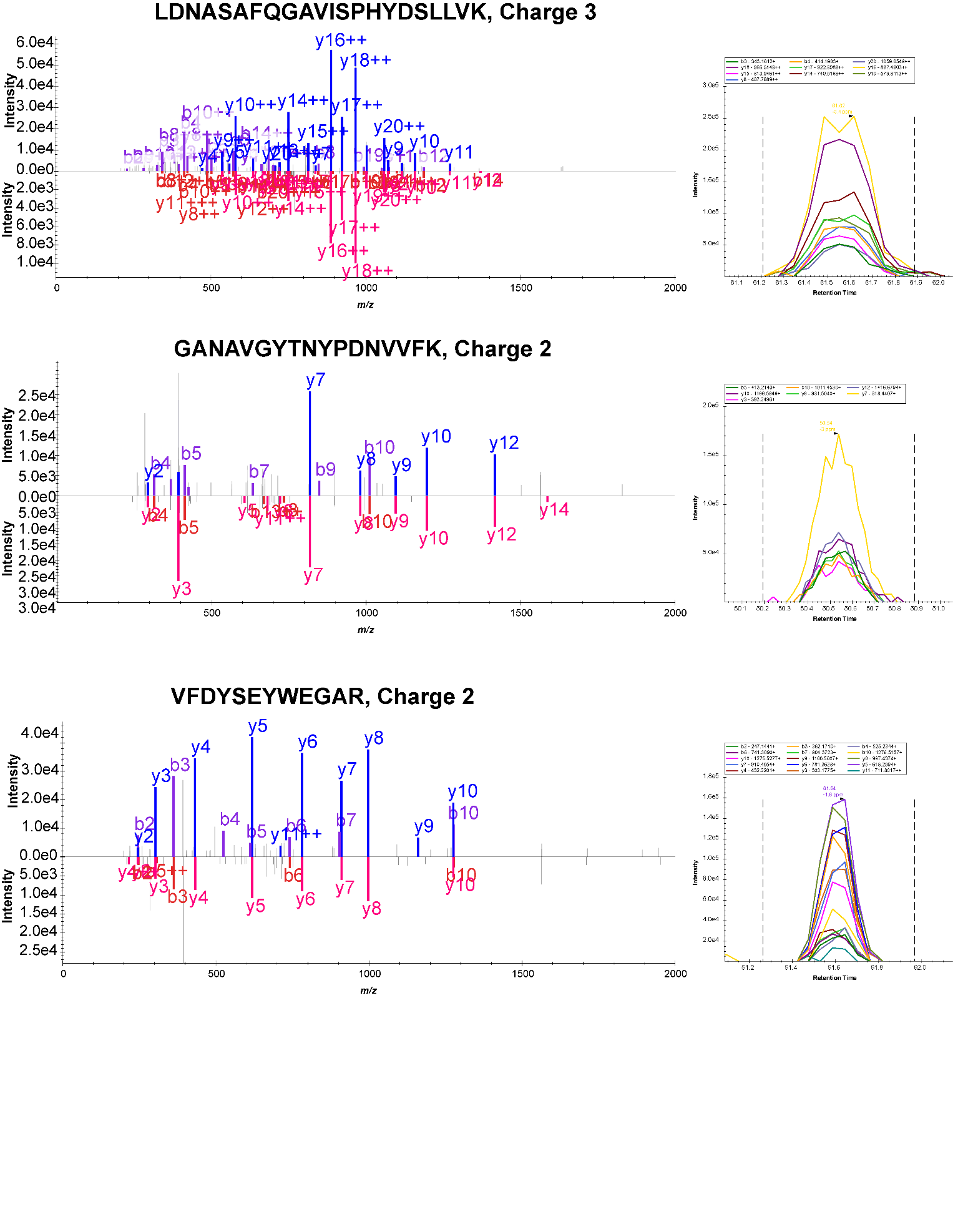

**Figure S5i** (continued, assayed peptides)

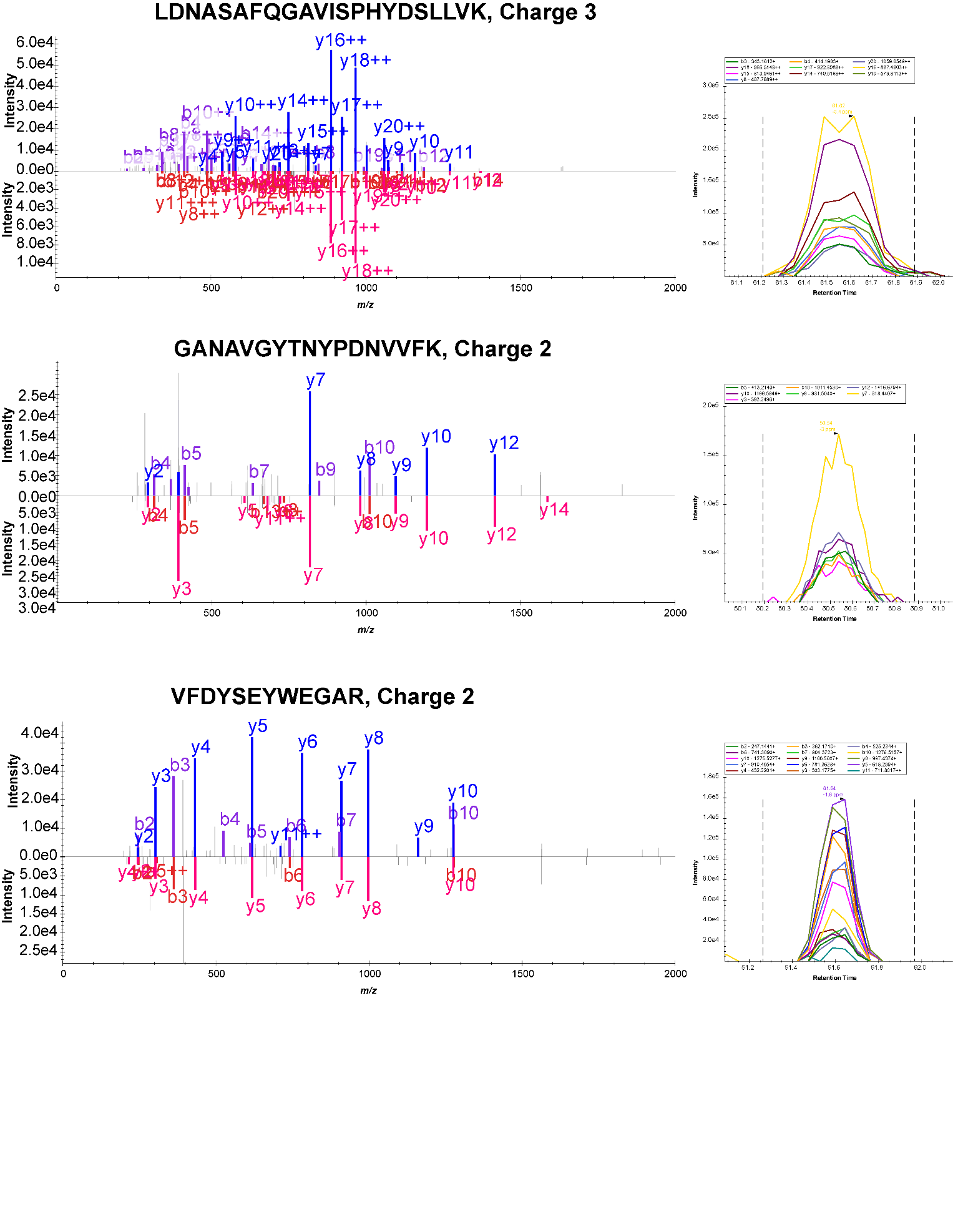

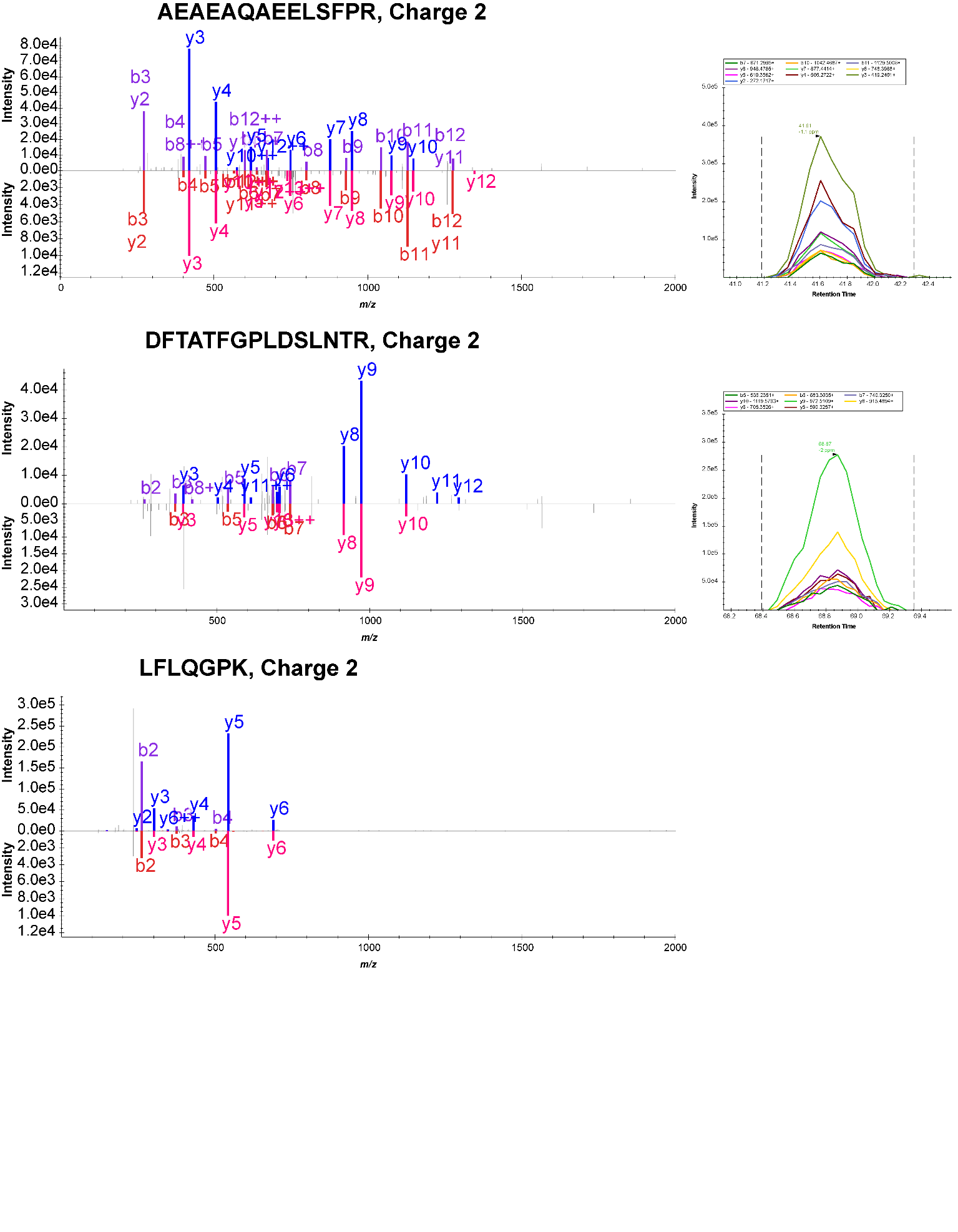

**Figure S5.** Selected PRM targets, MS^2^ spectra (top) and reference CID spectra (bottom).

**Figure S6a**

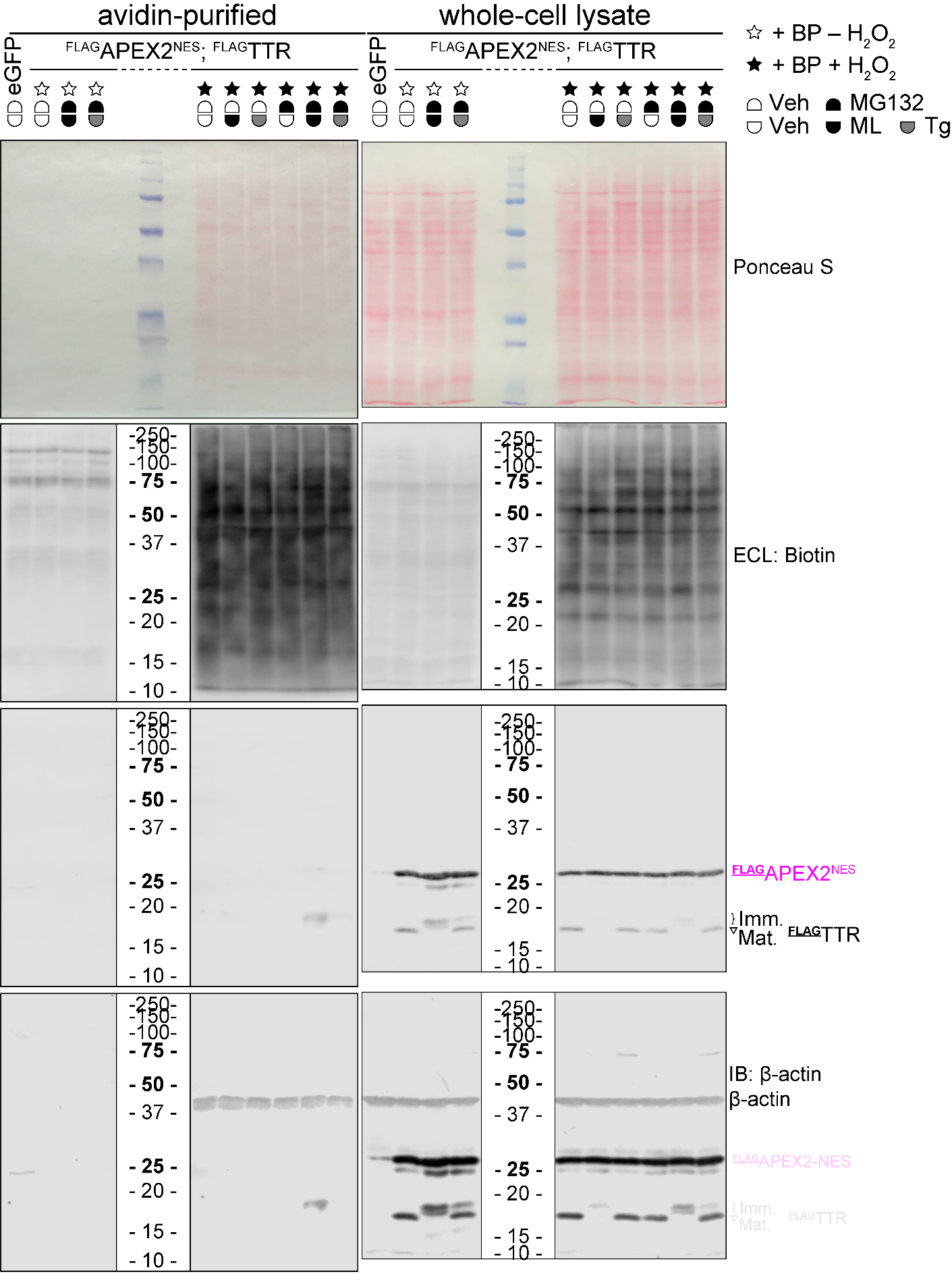

**Figure S6a (continued)**

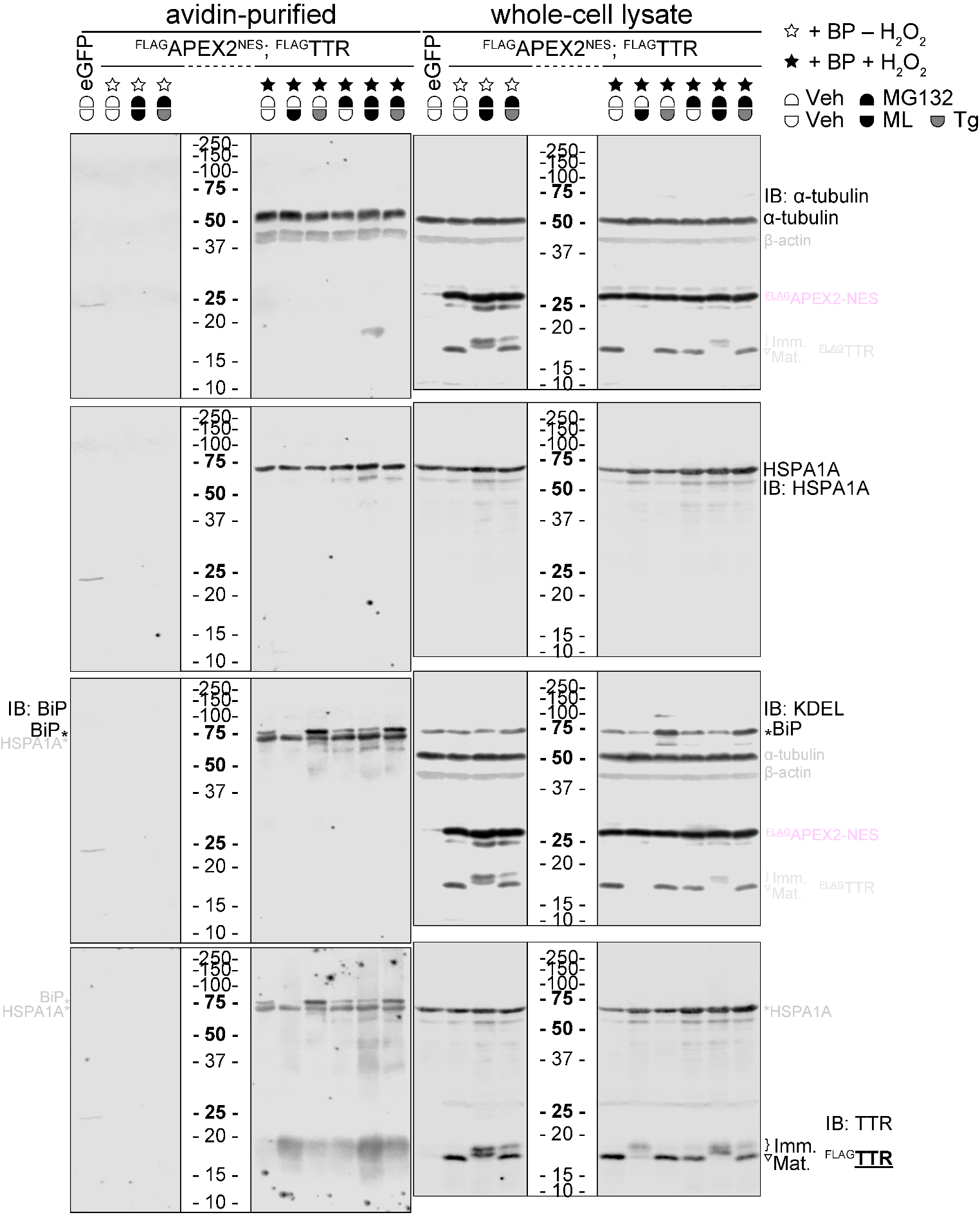

**Figure S6b**

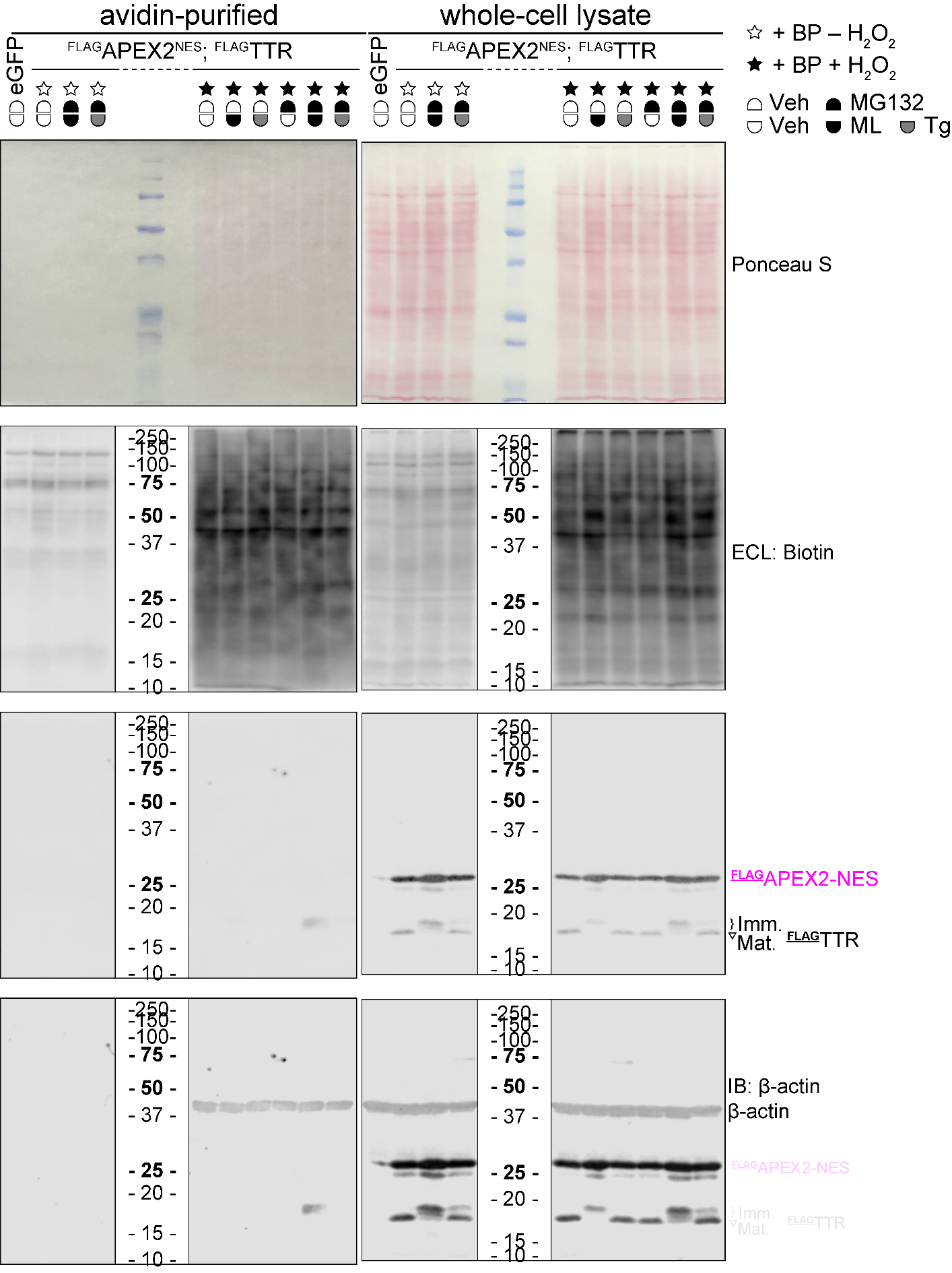

**Figure S6b (continued)**

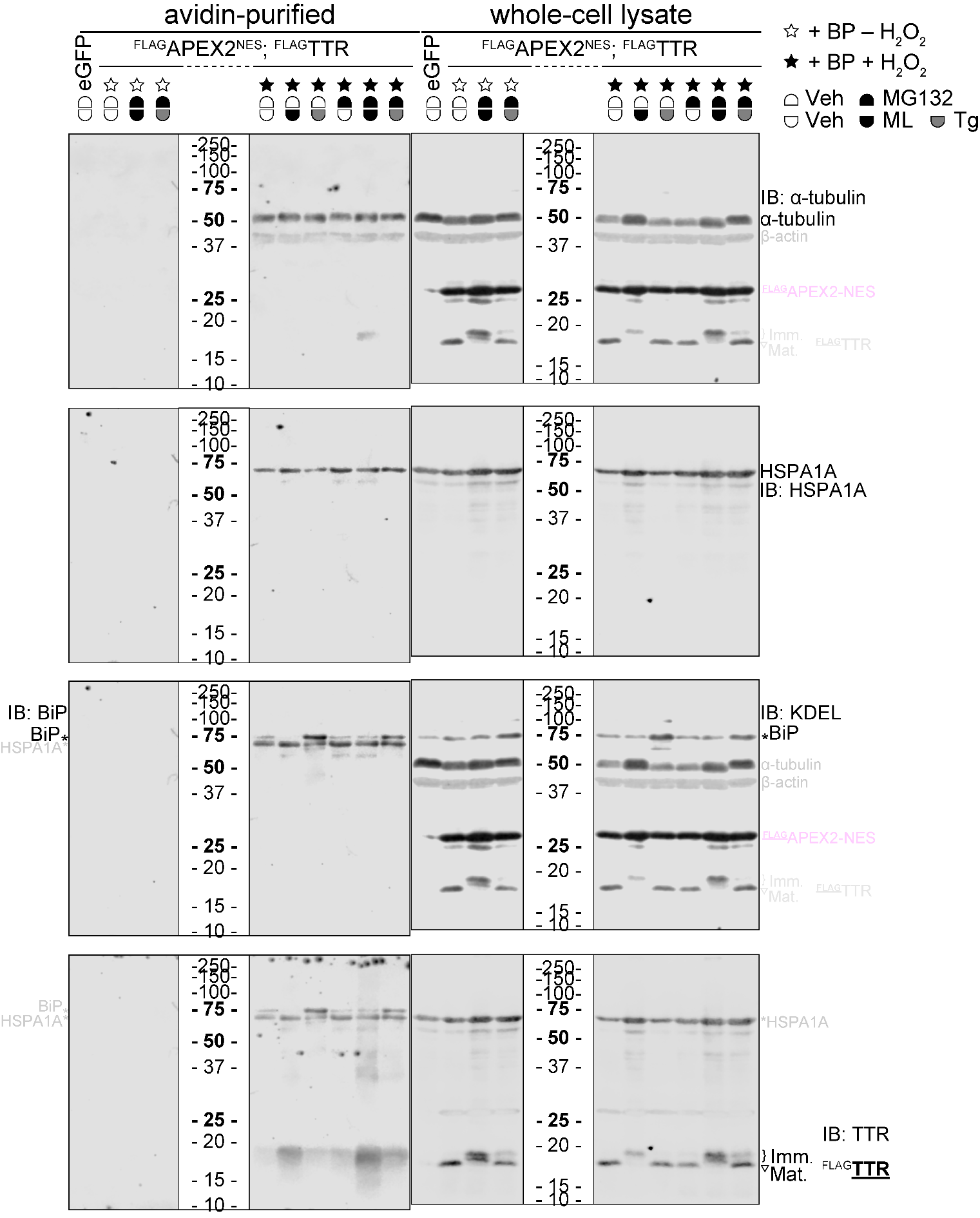

**Figure S6.** Ponceau S stain, ECL of biotin and IB Full blots related to **Figure 3**. HEK293T cells received 16-h 1 μM MG132, 25 nM ML or 50 nM Tg treatment as indicated. Ponceau S stains are displayed for protein loading control, and ECL of biotin is for biotin-phenol labeling efficiency given the same amount of total protein level. eGFP and some unlabeled samples were included to show background. IBs of assayed proteins in avidin-purifications (one half of each eluate sample) and corresponding whole cell lysates are presented in blotting order. BiP level in avidin-purifications were detected by rabbit anti-BiP polyclonal antibody and goat anti-rabbit IRDye800, instead of mouse anti-KDEL monoclonal antibody and goat anti-mouse IRDye680. The other half of each eluate sample was prepared for PRM analysis. **a**) Full blots of replicate 1. **b**) Full blots of replicate 2.

**Figure S7a**

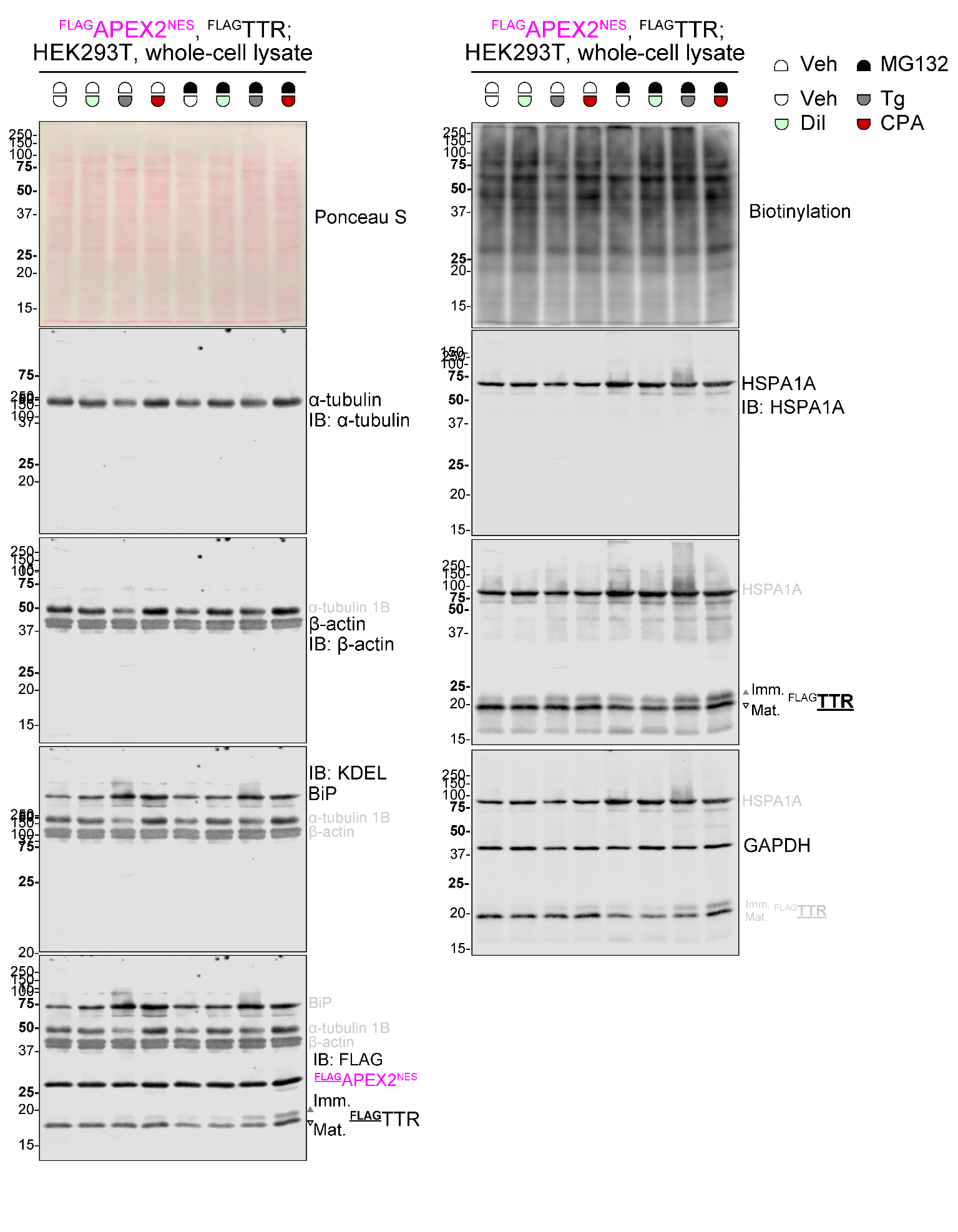

**Figure S7b**
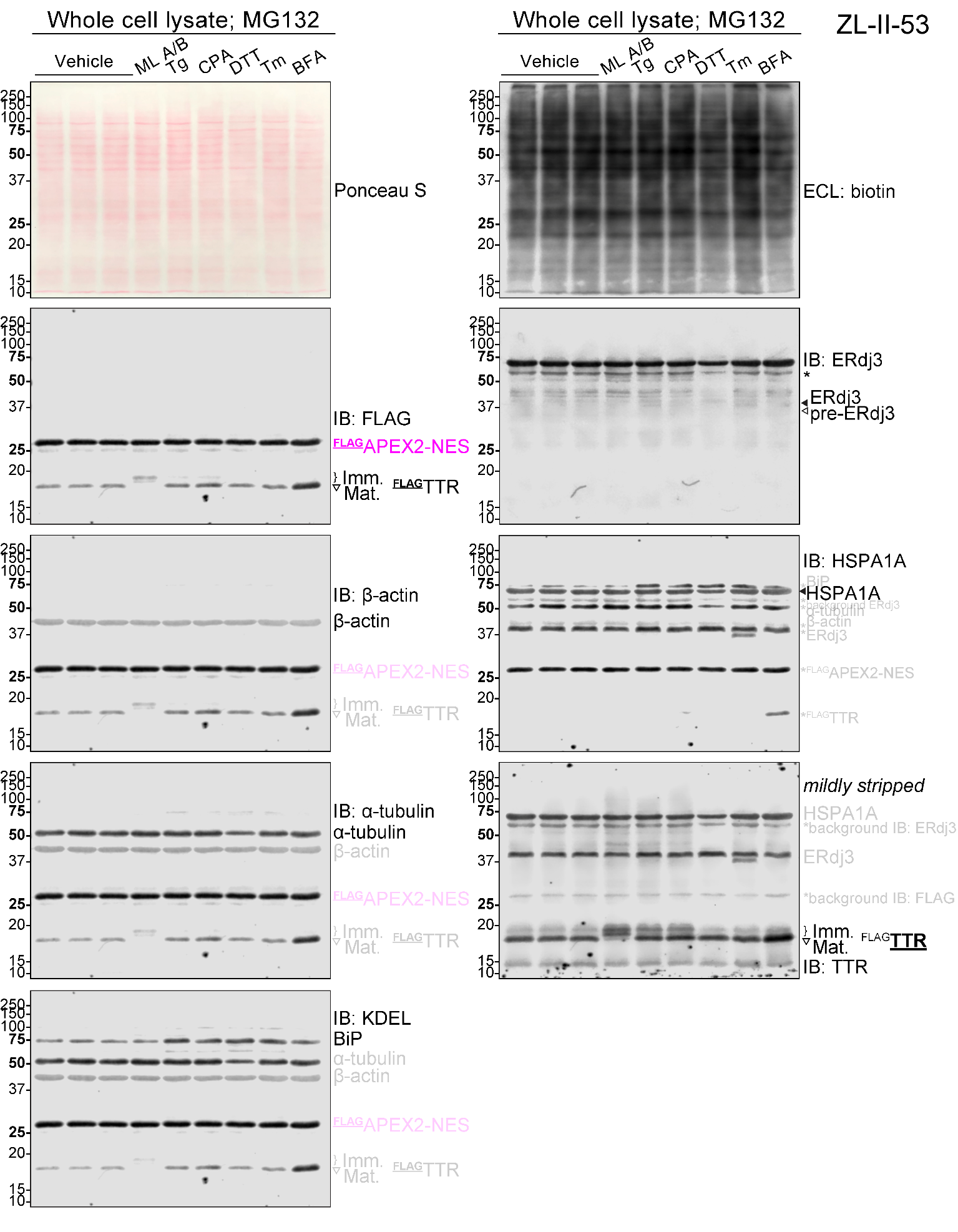
 **Figure S7.** Representative ponceau S stain, ECL of biotin and IB Full blots related to **Figures 4** and **5**. HEK293T cells received 16-h MG132, ER stressors or mycolactone A/B treatment as indicated, prior to proximity labeling. ER stress was confirmed by the upregulation of BiP (IB: KDEL). The inhibition of *N*-glycan synthesis during Tm treatment was confirmed by the appearance of the pre-ERdj3 band (*N*-glycan naïve form) from IB (IB: ERdj3, lane Tm). The inhibition of secretion by BFA was indicated by increased steady state ^FLAG^TTR level in lysate (IB: TTR, lane BFA). In these experiments, Ponceau S stains are displayed for protein loading control, and ECL of biotin is for BP labeling efficiency given the same amount of total protein level. IBs of assayed proteins in corresponding whole cell lysates are presented in blotting order. **a**) One of the two duplicates involving 30 μM diltiazem (Dil.), 50 nM Tg and 100 μM CPA. **b**) One experiment involving 1 μM MG132, 25 nM ML and ER stressors: Tg, 50 nM; CPA, 100 μM; DTT, 3 mM; Tm, 200 nM; BFA, 400 ng mL^–1^.

### **Table S1**. Drug treatment conditions (16 hours)

| **Drugs** | **Purpose** | **Final conc.** |
| --- | --- | --- |
| Mycolactone A/B (ML A/B,  0.2 mg mL^–1^ stock in EtOAc) | Sec61 inhibitor | 25 nM |
| MG132  (Selleckchem) | Proteasomal inhibitor | 1 μM |
| Thapsigargin  (Tg, Adipogen) | SERCA inhibitor | 50 nM |
| Tunicamycin (Tm, MP Biomedicals) | *N*-glycosylation inhibitor | 200 nM |
| Brefeldin A (BFA, Adipogen) | Secretion inhibitor | 400 ng mL^–1^ |
| 1,4-Dithiothreitol (DTT, Fisher) | Disulfide bond cleavage | 3 mM |
| 2-Deoxy-d-glucose (2-DG, CHEM-IMPEX INT’L INC.) | *N*-glycosylation inhibitor | 15 μM |
| Cyclopiazonic acid (CPA, Millipore) | SERCA inhibitor | 100 μM |
| Diltiazem HCl (Dil., Sigma) | Calcium channel blocker | 30 μM |
